# Charting Brain Growth and Aging at High Spatial Precision

**DOI:** 10.1101/2021.08.08.455487

**Authors:** Saige Rutherford, Charlotte Fraza, Richard Dinga, Seyed Mostafa Kia, Thomas Wolfers, Mariam Zabihi, Pierre Berthet, Amanda Worker, Serena Verdi, Derek Andrews, Laura Han, Johanna Bayer, Paola Dazzan, Phillip McGuire, Roel T. Mocking, Aaart Schene, Brenda W. Pennix, Chandra Sripada, Ivy F. Tso, Elizabeth R. Duval, Soo-Eun Chang, Mary Heitzeg, S. Alexandra Burt, Luke Hyde, David Amaral, Christine Wu Nordahl, Ole A. Andreasssen, Lars T. Westlye, Roland Zahn, Henricus G. Ruhe, Christian Beckmann, Andre F. Marquand

## Abstract

Defining reference models for population variation, and the ability to study individual deviations is essential for understanding inter-individual variability and its relation to the onset and progression of medical conditions. In this work, we assembled a reference cohort of neuroimaging data from 82 sites (N=58,836; ages 2-100) and use normative modeling to characterize lifespan trajectories of cortical thickness and subcortical volume. Models are validated against a manually quality checked subset (N=24,354) and we provide an interface for transferring to new data sources. We showcase the clinical value by applying the models to a transdiagnostic psychiatric sample (N=1,985), showing they can be used to quantify variability underlying multiple disorders whilst also refining case-control inferences. These models will be augmented with additional samples and imaging modalities as they become available. This provides a common reference platform to bind results from different studies and ultimately paves the way for personalized clinical decision making.

## Introduction

Since their introduction more than a century ago, normative growth charts have become fundamental tools in pediatric medicine and also in many other areas of anthropometry (***Cole (2012***)). They provide the ability to quantify individual variation against centiles of variation in a reference population, which shifts focus away from group-level (e.g., case-control) inferences to the level of the individual. This idea been adopted and generalized in clinical neuroimaging and normative modelling is now established as an effective technique for providing inferences at the level of the individual in neuroimaging studies (***Marquand et al. (2016***, 2019)).

Although normative modelling can be used to estimate many different kinds of mappings – for example between behavioral scores and neurobiological readouts – normative models of brain development and aging are appealing considering that many brain disorders are grounded in atypical trajectories of brain development (***Insel (2014***)) and the association between cognitive decline and brain tissue in ageing and neurodegenerative diseases (***Jack et al. (2010***); ***Karas et al. (2004***)). Indeed, normative modelling has been applied in many different clinical contexts, including charting the development of infants born pre-term (***Dimitrova et al. (2020***)) and dissecting the biological heterogeneity across cohorts of individuals with different brain disorders including schizophrenia, bipolar disorder, autism and attention deficit/hyperactivity disorder (***Bethlehem et al. (2020***); ***Wolfers et al. (2021***); ***Zabihi et al. (2019***)).

A hurdle to the widespread application of normative modelling is a lack of well-defined reference models to quantify variability across the lifespan and to compare results from different studies. Such models should: (i) accurately model population variation across large samples; (ii) be derived from widely accessible measures; (iii) provide the ability to be updated as additional data come on-line and (vi) be supported by easy-to-use software tools. In addition, they should quantify brain development and ageing at a high spatial resolution, so that different patterns of atypicality can be used to stratify cohorts and predict clinical outcomes with maximum precision. The purpose of this work is to introduce a set of reference models that satisfy these criteria.

To this end, we assemble a large neuroimaging dataset (Table 1) from 58,836 individuals across 82 scan sites covering the human lifespan (aged 2-100, Figure 1A) and fit normative models for cortical thickness and subcortical volumes derived from Freesurfer (version 6.0). We show the clinical utility of these models in a large transdiagnostic psychiatric sample (N=1,985). To maximize the utility of this contribution, we distribute model coefficients freely along with a set of [software](https://github.com/predictive-clinical-neuroscience/braincharts) tools to enable researchers to derive subject-level predictions for new datasets against a set of common reference models.

**Table 1.** Sample Description and Demographics. mQC refers to the manual quality checked subset of the full sample. ‘All’ rows = Train + Test. Clinical refers to the transdiagnostic psychiatric sample (diagnostic details in Figure 2A).

|  |  | N (subjects) | N (sites) | Sex (%F, %M) | Age (Mean, s.d) |
| --- | --- | --- | --- | --- | --- |
| **Full** | All | 58,836 | 82 |  |  |
|  | Training set | 29,418 | 82 | 51.1, 48.9 | 46.9, 24.4 |
|  | Test set | 29,416 | 82 | 50.9, 49.1 | 46.9, 24.4 |
| **mQC** | All | 24,354 | 59 |  |  |
|  | Training set | 12,177 | 59 | 50.2, 49.8 | 30.2, 24.1 |
|  | Test set | 12,177 | 59 | 50.4, 49.4 | 30.1, 24.2 |
| **Clinical** | Test set | 1,985 | 24 | 38.9, 61.1 | 30.5, 14.1 |

**Figure 1.**
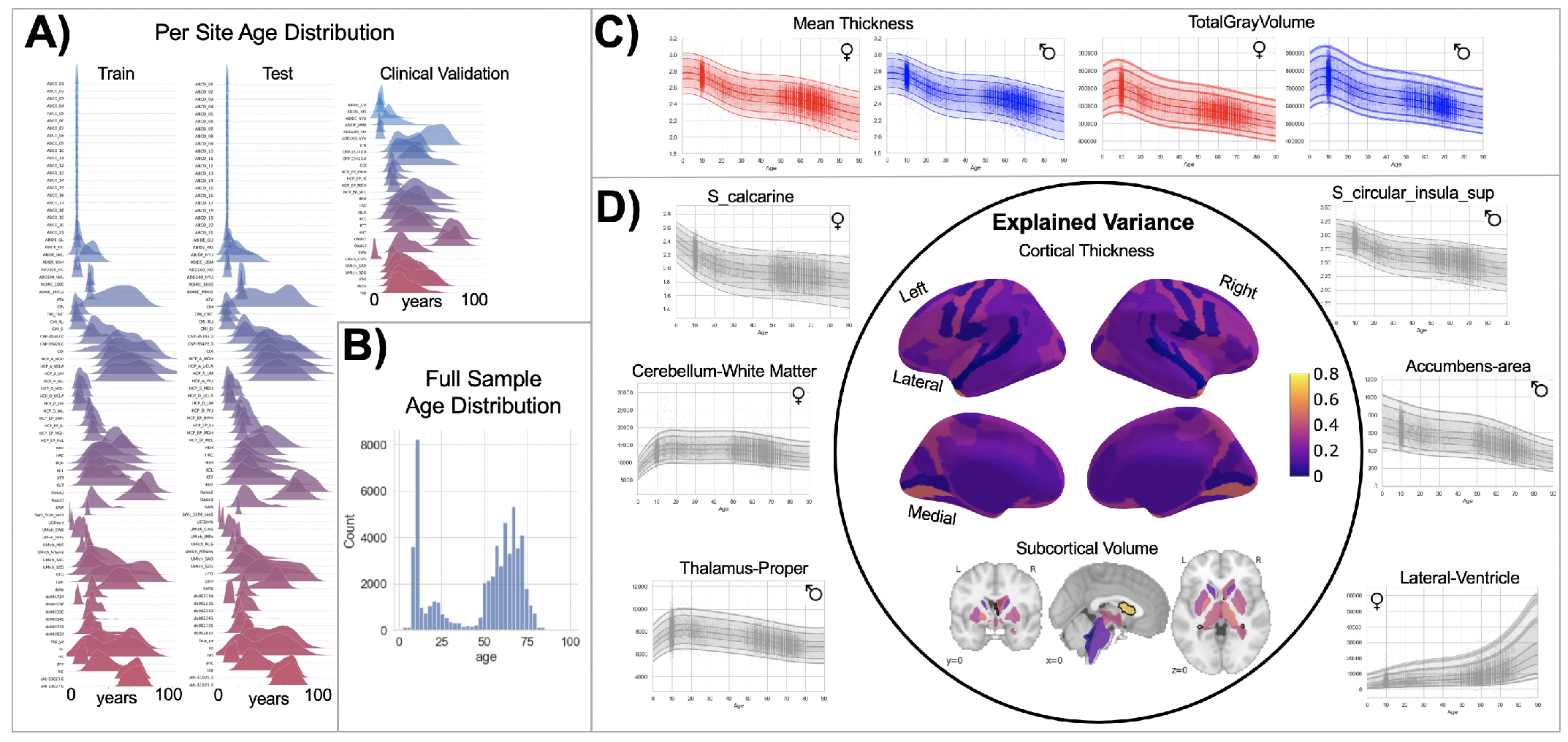
Normative Model Overview. A) Age distribution (x-axis) of each site (y-axis) in the full model train and test sets. B) Age distribution of each site in the clinical test set. C-D) Examples of lifespan trajectories of brain regions. Age is shown on x-axis and predicted thickness (or volume) values are on the y-axis. Centiles of variation are plotted for each region. In panel C, we show that sex differences between females (red) and males (blue) are most pronounced when modeling large scale features such as mean cortical thickness across the entire cortex or total gray matter volume. These sex differences manifest as a shift in the mean in that the shape of these trajectories is the same for both sexes, as determined by sensitivity analyses where separate normative models were estimated for each sex. The explained variance (in full test set) of the whole cortex and subcortex is highlighted inside the circle of panel D. All plots within the circle share the same color scale. E) The distribution of evaluation metrics in the full test set, separated into left and right hemispheres and subcortical regions, with the skew and excess kurtosis being measures that depict the accuracy of the estimated shape of the model, ideally both of these would be around zero.

## Results

We split the available data into training and test sets, stratifying the split by site, such that all sites are equally represented in the training and test sets (Table 1, Supplementary Table 2, Supplementary Table 3, and Supplementary Table 4). After careful automated and manual quality checking procedures (see methods), we then fit a normative model using a set of covariates (age, gender, and fixed effects for site) to predict cortical thickness and subcortical volume for each parcel in a high resolution atlas derived from the Freesurfer software package (***Destrieux et al. (2010***)). We employed a warped Bayesian linear regression model so as to accurately model both non-linear effects and non-Gaussian distribution of the imaging phenotype (***Fraza et al. (2021***)), whilst accounting for the well-known effects of different scanners on neuroimaging data (***Bayer et al. (2021***); ***Kia et al. (2021***)). These models are summarized in Figure 3, Supplementary Table 5, Supplementary Table 6, Supplementary Table 7, and Supplementary Table 8.

We validate our models with several careful procedures: first, we report out of sample metrics; second, we perform a supplementary analysis on a subset of participants for whom input data had undergone manual quality checking by an expert rater (Table 1 - mQC). Third, each model fit was evaluated using metrics (Figure 3, Supplementary Table 5, Supplementary Table 6, Supplementary Table 7, and Supplementary Table 8) that quantify central tendency and distributional accuracy (***Dinga et al. (2021***); ***Fraza et al. (2021***)). We also estimated separate models for males and females, which indicate that sex effects are adequately modeled using a global offset. Finally, to facilitate independent validation, we packaged pretrained models and code for transferring to new samples into an [open resource](https://pcntoolkit.readthedocs.io/en/latest/pages/tutorial_braincharts_apply_nm.html) for use by the community and demonstrate how to transfer the models to new samples (i.e., data not present in the initial training set).

Our models provide the opportunity for mapping the diverse trajectories of different brain areas. Several examples are shown in Figure 1 C and D which align with known patterns of development and aging (***Ducharme et al. (2016***); ***Gogtay et al. (2004***); ***Tamnes et al. (2010***)). Moreover, across the cortex and subcortex our model fits well, explaining up to 80% of the variance (minimum 12%) out of sample (Figure 3, Supplementary Table 5, Supplementary Table 6, Supplementary Table 7, and Supplementary Table 8 for full details).

A goal of this work is to develop normative models that can be applied to many different clinical conditions. To showcase this, we apply the model to a transdiagnostic psychiatric cohort (Table 1 – Clinical; Figure 2A) resulting in personalized, whole-brain deviation maps that can be used to understand inter-individual variability (e.g., for stratification) and to quantify group separation (e.g., case-control effects). To demonstrate this, for each clinical group, we summarized the individual deviations within that group by computing the proportion of subjects that have deviations in each region and compare to matched (same sites) controls in the test set (Figure 2B-C). Additionally, we performed case-control comparisons on the raw cortical thickness and subcortical volumes, and on the deviation maps (Figure 2D), again against a matched sample from the test set. This demonstrates the advantages of using normative models for investigating individual differences in psychiatry, i.e., quantifying clinically relevant information at the level of each individual. For most diagnostic groups, the z-statistics derived from the normative deviations also provided stronger case-control effects than the raw data. This shows the importance of accurate modeling of population variance across multiple clinically relevant dimensions. The individual-level deviations provide complimentary information to the group effects, which aligns with previous work (***Wolfers et al. (2020***); ***Zabihi et al. (2020***); ***Wolfers et al. (2018***)). We note that a detailed description of the clinical significance of our findings is beyond the scope of this work and will be presented separately.

**Figure 2.**
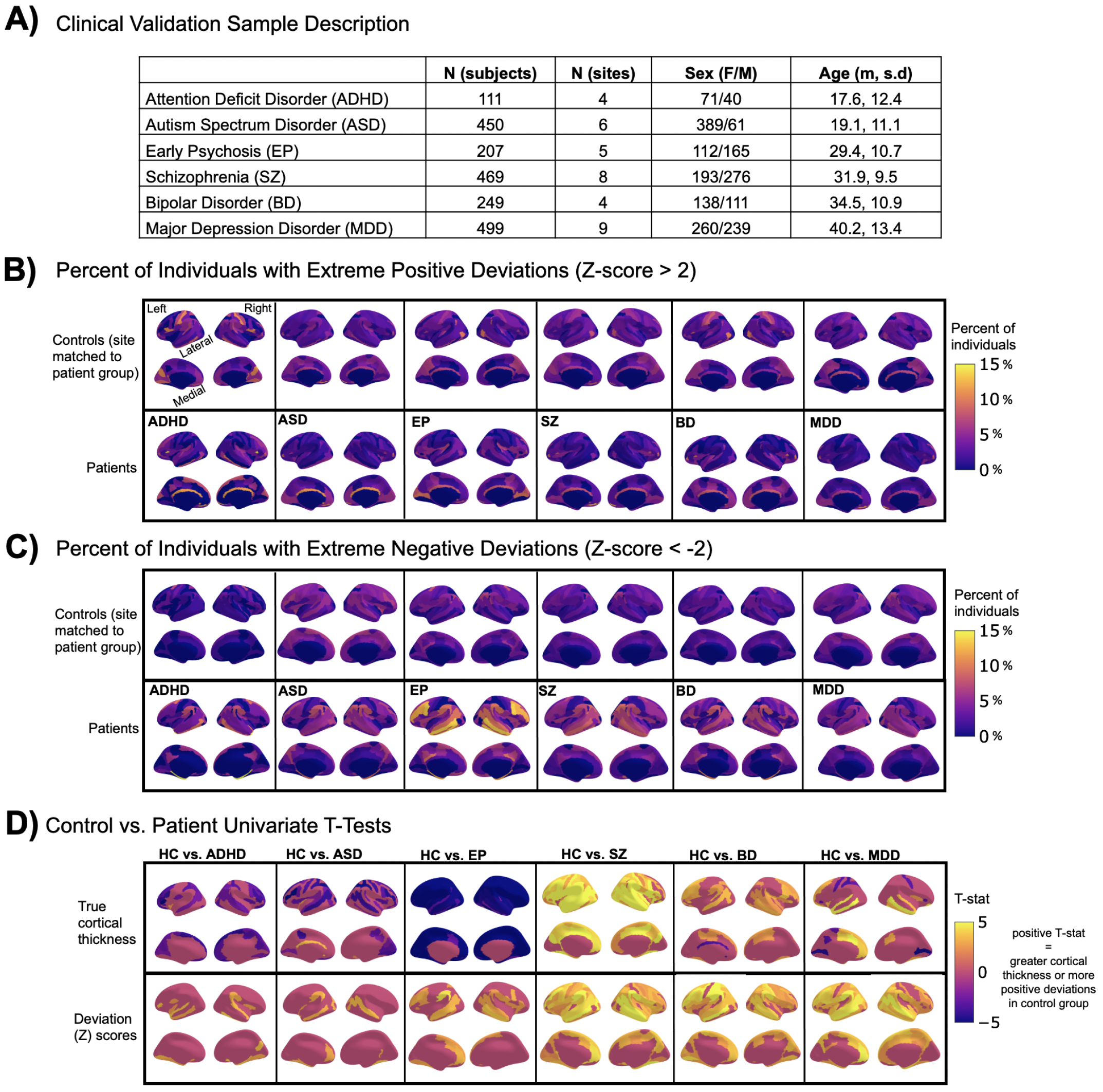
Normative Modeling in Clinical Cohorts. Reference brain charts were transferred to several clinical samples (panel A). Patterns of extreme deviations were summarized for each clinical group and compared to matched control groups (from the same sites). Panel B) shows extreme positive deviations (thicker/larger than expected) and panel C) shows the extreme negative deviation (thinner/smaller than expected) patterns. Panel D) shows the significant (FDR corrected p<0.05) results of classical case-control methods (mass-univariate t-tests) on the true cortical thickness data (top row) and on the deviations scores (bottom row). There is unique information added by each approach which becomes evident when noticing the maps in panels B-D are not identical. ADHD=Attention Deficit Hyperactive Disorder, ASD=Autism Spectrum Disorder, EP=Early Psychosis, SZ=Schizophrenia, BD=Bipolar Disorder, MDD=Major Depressive Disorder.

## Discussion

In this work we create lifespan brain charts of cortical thickness and subcortical volume derived from structural MRI, to serve as reference models. Multiple datasets were joined to build a mega-site lifespan reference cohort to provide good coverage of the lifespan. We applied the reference cohort models to clinical datasets and demonstrate the benefits of normative modeling in addition to standard case-control comparisons. All models, including [documentation](https://pcntoolkit.readthedocs.io/en/latest/pages/tutorial_braincharts_apply_nm.html) and [code](https://github.com/predictive-clinical-neuroscience/braincharts), are made available to the research community.

We identify three main strengths of our approach. First, our large lifespan dataset provides high anatomical specificity, necessary for discriminating between conditions, predicting outcomes, and stratifying subtypes. Second, our models are flexible in that they can model non-Gaussian distributions, can easily be transferred to new sites, and are built on validated analytical techniques and software tools (***Fraza et al. (2021***); ***Kia et al. (2021***); ***Marquand et al. (2019***)). Third, we show the general utility of this work in that it provides the ability to map individual variation whilst also improving case control inferences across multiple disorders.

In recent work, a large consortium established lifespan brain charts that are complementary to our approach (***Bethlehem et al. (2021***)). Benefits of their work include the use of a large sample and good coverage of the peri-natal life stages. This was used to provide estimates of brain growth in terms of global features (e.g., brain volume), which is valuable for applications where quantifying global brain development or ageing is of interest but has limited spatial precision. In contrast, we focus on providing spatially specific estimates across the post-natal lifespan which provides fine-grained anatomical estimates of deviation that may be valuable for understanding the biological basis for mental disorders where individual patterns are widespread (e.g., not all individuals deviate in the same regions).

We also identify limitations of this work. First, we view the word “normative” as problematic. This language implies that there are normal and abnormal brains, a potentially problematic assumption. As indicated in Figure 2, there is considerable individual variability and heterogeneity among trajectories. We encourage the use of the phrase ‘reference cohort’ over ‘normative model’. Second, to provide coverage of the lifespan the curated dataset is based on aggregating existing data, meaning there is unavoidable sampling bias. Race, education, and socioeconomic variables were not fully available for all included datasets, however, given that data were compiled from research studies, they are likely samples drawn predominantly from Western, Educated, Industrialized, Rich, and Democratic (WEIRD) societies (***Henrich et al. (2010***)) and future work should account for these factors. By sampling both healthy population samples and case-control studies, we achieve a reasonable estimate of variation across individuals, however, downstream analyses should consider the nature of the reference cohort and whether it is appropriate for the target sample. Finally, although the models presented in this study are comprehensive, they are only the first step, and we will augment our [repository](https://github.com/predictive-clinical-neuroscience/braincharts/tree/master/models) with more diverse data, different features and modelling advances as these become available.

## Methods and Materials

Data from 82 sites were combined to create the initial full sample. These sites are described in detail in Supplemental Table 5, including the sample size, age (mean and standard deviation), and sex distribution of each site. Many sites were pulled from publicly available data sets including [ABCD](https://abcdstudy.org/), [ABIDE](http://fcon_1000.projects.nitrc.org/indi/abide/), ADHD200, [CAMCAN](https://www.cam-can.org/index.php?content=dataset), [CMI-HBN](http://fcon_1000.projects.nitrc.org/indi/cmi_healthy_brain_network/), [HCP-Aging](https://www.humanconnectome.org/study/hcp-lifespan-aging), [HCP-Development](https://www.humanconnectome.org/study/hcp-lifespan-development), [HCP-Early Psychosis](https://www.humanconnectome.org/study/human-connectome-project-for-early-psychosis), HCP-Young Adult, [IXI](https://brain-development.org/ixi-dataset/), [NKI-RS](http://fcon_1000.projects.nitrc.org/indi/enhanced/), [Oasis](https://www.oasis-brains.org/), [OpenNeuro](https://openneuro.org/), [PNC](https://www.med.upenn.edu/bbl/philadelphianeurodevelopmentalcohort.html), [SRPBS](https://bicr-resource.atr.jp/srpbsopen/), and [UK Biobank](https://www.ukbiobank.ac.uk/). Other included data come from studies conducted at the University of Michigan (***Duval et al. (2018***); ***Rutherford et al. (2020***); ***Tomlinson et al. (2020***); ***Tso et al. (2021***); ***Weigard et al. (2021***); ***Zucker et al. (1996***)), University of California Davis (***Nordahl et al. (2020***)), University of Oslo (***Nesvåg et al. (2017***)), King’s College London (***Green et al. (2012***); ***Lythe et al. (2015***)), and Amsterdam University Medical Center (***Mocking et al. (2016***)). Full details regarding sample characteristics, diagnostic procedures and acquisition protocols can be found in the publications associated with each of the studies. Equal sized training and testing data sets (split half) were created using scikit-learn’s train_test_split function, stratifying on the site variable. It is important to stratify based on site, not only study (***Bethlehem et al. (2021***)), as many of the public studies (i.e., ABCD) include several sites, thus modeling study does not adequately address MRI scanner confounds.

The clinical validation sample consisted of a subset of the full data set (described in detail in Figure 2A and Supplemental Table 3). Studies (sites) contributing clinical data included: Autism Brain Imaging Database Exchange (ABIDE GU, KKI, NYU, USM), ADHD200 (KKI, NYU), CNP, SRPBS (CIN, COI, KTT, KUT, HKH, HRC, HUH, SWA, UTO), Delta (AmsterdamUMC), Human Connectome Project Early Psychosis (HCP-EP BWH, IU, McL, MGH), KCL, University of Michigan Schizophrenia Gaze Processing (UMich_SZG), and TOP (University of Oslo).

In addition to the sample-specific inclusion criteria, inclusion criteria for the full sample was based on participants having basic demographic information (age and sex), a T1-weighted MRI volume, and Freesurfer output directories that include summary files which represent left and right hemisphere cortical thickness values of the Destrieux parcellation and subcortical volumetric values (aseg.stats, lh.aparc.a2009s.stats, rh.aparc.a2009s.stats). Freesurfer image analysis suite (version 6.0) was used for cortical reconstruction and volumetric segmentation for all studies. The technical details of these procedures are described in prior publications (***Dale et al. (1999***); ***Fischl and Dale (2000***); ***Fischl et al. (2002***)). UK Biobank was the only study for which Freesurfer was not run by the authors. Freesurfer functions aparcstats2table and asegstats2table were run to extract cortical thickness from the Destrieux parcellation (***Destrieux et al. (2010***)) and subcortical volume for all participants into CSV files. These files were inner merged with the demographic files, using Pandas, and NaN rows were dropped.

Quality control (QC) is an important consideration for large samples and is an active research area (***Alfaro-Almagro et al. (2018***); ***Klapwijk et al. (2019***); ***Rosen et al. (2018***)). We consider manual quality checking of images both prior to and after preprocessing to be the gold standard. However, this is labor intensive and prohibitive for very large samples. Therefore, in this work we adopt a pragmatic and multi-pronged approach to QC. First, a subset of the full data set underwent manual quality checking (mQC, described in Supplemental Table 4) by author S.R. using Papaya, a JavaScript based image viewer. Each subject’s T1w volume was viewed in 3D volumetric space, with the Freesurfer brain.finalsurfs file as an overlay, to check for obvious quality issues such as excessive motion, ghosting or ringing artifacts. Example scripts used for quality checking can be found on GitHub. The ABCD study data distributes a variable (freesqc01.txt; fsqc_qc==1/0) that represents manual quality checking (pass/fail) of the T1w volume and Freesurfer data, thus this data set was added into our manual quality checked data set bringing the sample size to 24,354 individuals passing manual quality checks. Although this has a reduced sample, we consider this to be a gold standard sample in that every single scan has been checked manually. All inferences reported in this manuscript were validated against this sample. Second, for the full sample, we adopted an automated QC procedure that quantifies image quality based on the Freesurfer Euler Characteristic (EC), which has been shown to be an excellent proxy for manual labelling of scan quality (***Monereo-Sánchez et al. (2021***); ***Rosen et al. (2018***)) and is the most important feature in automated scan quality classifiers (***Klapwijk et al. (2019***)). Since the distribution of the EC varies across sites, we adopt a simple approach that involves scaling and centering the distribution over the EC across sites and removing samples in the tail of the distribution (see (***Kia et al. (2021***)) for details).

Normative modeling was run using python 3.8 and the [PCNtoolkit](https://pcntoolkit.readthedocs.io/) package (version 0.20). Bayesian Linear Regression (BLR) with likelihood warping was used to predict cortical thickness and subcortical volume from a vector of covariates (age, sex, site). For a complete mathematical description and explanation of this implementation see (***Fraza et al. (2021***)). Briefly, for each brain region of interest (cortical thickness or subcortical volume), y is predicted as:

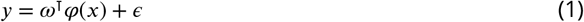

Where *ω* ^T^ is the estimated weight vector, *φ* (*x*) is a basis expansion of the of covariate vector *x*, consisting of a B-spline basis expansion (cubic spline with 5 evenly spaced knots) to model nonlinear effects of age, and *ϵ* = *η*(*θ, β*) a Gaussian noise distribution with mean zero and noise precision term *β)* (the inverse variance). A likelihood warping approach (***Rios and Tobar (2019***); ***Snelson et al. (2003***)) was used to model non-Gaussian effects. This involves applying a bijective nonlinear warping function to the non-Gaussian response variables to map them to a Gaussian latent space where inference can be performed in closed form. We employed a ‘sinarcsinsh’ warping function, which is equivalent to the SHASH distribution commonly used in the generalized additive modeling literature (***Jones and Pewsey (2009***)) and which we have found to perform well in prior work (***Dinga et al. (2021***); ***Fraza et al. (2021***)). Site variation was modeled using fixed effects, which we have shown in prior work provides relatively good performance (***Kia et al. (2021***)), although random effects for site may provide additional flexibility at higher computational cost. A fast numerical optimization algorithm was used to optimize hyperparameters (L-BFGS). Computational complexity of hyperparameter optimization was controlled by minimizing the negative log likelihood. Deviation scores (Z-scores) are calculated for the 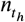subject, and 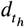 *s*brain area, in the test set as:

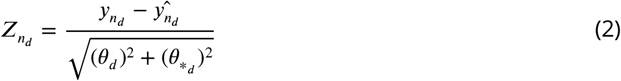

Where 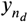 is the true response, 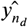 is the predicted mean, 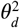 is the estimated noise variance (reflecting uncertainty in the data), and 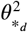 is the variance attributed to modeling uncertainty. Model fit for each brain region was evaluated by calculating the explained variance (which measures central tendency), the mean squared log-loss (MSLL, central tendency and variance) plus skew and kurtosis of the deviation scores (equation 2) which measures how well the shape of the regression function matches the data (***Dinga et al. (2021***)). Note that for all models, we report out of sample metrics (Figure 3, Supplementary Table 5, Supplementary Table 6, Supplementary Table 7, and Supplementary Table 8).

**Figure 3.**
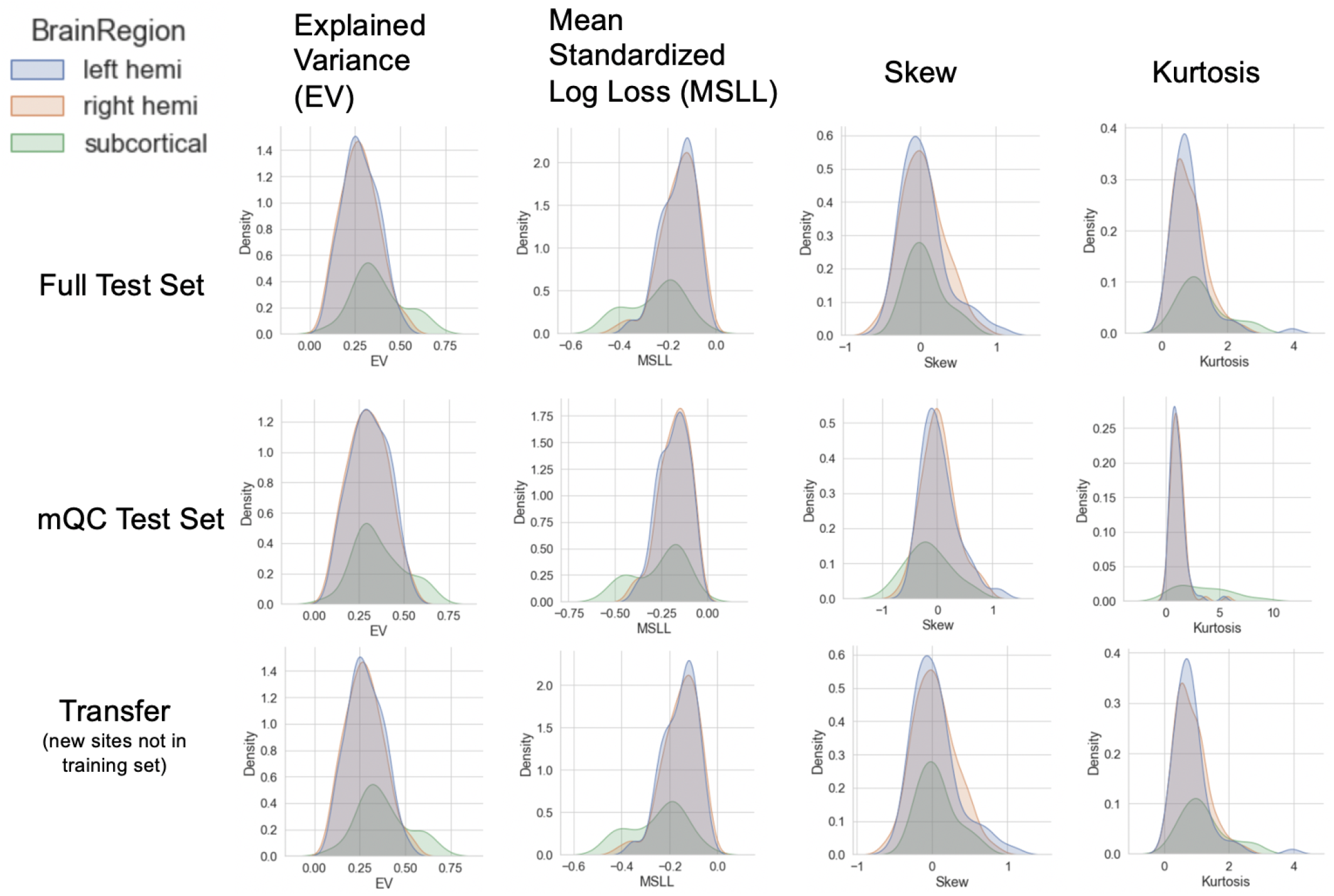
Evaluation Metrics Across All Test Sets. The distribution of evaluation metrics in 3 different test sets (full, mQC, and Transfer) separated into left and right hemispheres and subcortical regions, with the skew and excess kurtosis being measures that depict the accuracy of the estimated shape of the model, ideally both of these would be around zero.

To provide a summary of individual variation within each clinical group, deviation scores were summarized for each clinical group (Figure 2B-C) by first separating them into positive and negative deviations, counting how many subjects had an extreme deviation (positive extreme deviation defined as *z >* 2, negative extreme deviation as *z <* −2) at a given ROI, and then dividing by the group size to show the percentage of individuals with extreme deviations at that brain area. Controls from the same sites as the patient groups were summarized in the same manner for comparison. We also performed classical case vs. control group difference testing on the true data and on the deviation scores (Figure 2D) and thresholded results at a Benjamini-Hochberg false discovery rate (FDR) of *p <* 0.05. Note that in both cases, we directly contrast each patient group to their matched controls to avoid nuisance variation confounding any reported effects (e.g., sampling characteristics, demographic differences).

All pretrained models and code are shared online with straightforward directions for transferring to new sites. Given a new set of data (e.g., sites not present in the training set), this is done by first applying the warp parameters estimating on the training data to the new dataset, adjusting the mean and variance in the latent Gaussian space, then (if necessary) warping the adjusted data back to the original space, which is similar to the approach outlined in (***Dinga et al. (2021***)). Note that to remain unbiased, this should be done on a held-out calibration dataset. To illustrate this procedure, we apply this approach to predicting a subset of sites in the 1000 functional connectomes project (***Biswal et al. (2010***)) that were not used during the model estimation step. These results are reported in Supplemental Figure 4 (bottom row). We also distribute scripts for this purpose in the GitHub Repository associated with this manuscript. Furthermore, to promote the use of these models and remove barriers to using them, we have set up access to the pretrained models and code for transferring to users’ own data, using [Google Colab](https://colab.research.google.com/github/predictive-clinical-neuroscience/braincharts/blob/master/scripts/apply_normative_models.ipynb), a free, cloud-based platform for running python notebooks. This eliminates the need to install python/manage package versions and only requires users to have a personal computer with stable internet connection.

## Acknowledgments

This research was supported by grants from the European Research Council (ERC, grant “MENTAL-PRECISION” 10100118 and “BRAINMINT” 802998), the Wellcome Trust under an Innovator award (“BRAINCHART”, 215698/Z/19/Z) and a Strategic Award (098369/Z/12/Z), the Dutch Organisation for Scientific Research (VIDI grant 016.156.415) the Research Council of Norway (223273, 249795, 298646, 300768, 276082), the South-Eastern Norway Regional Health Authority (2014097, 2015073, 2016083, 2019101), the KG Jebsen Stiftelsen, an Autism Center of Excellence grant awarded by the National Institute of Child Health and Development (NICHD) (P50 HD093079) as well as the National Institute of Mental Health (R01MH104438, R01MH103371). TW also gratefully acknowledges the Niels Stensen Fellowship as well as the European Union’s Horizon 2020 research and innovation programme under the Marie Sklodowska-Curie Grant agreement No. 895011. RZ was funded by Medical Research Council grant (G0902304). IFT was funded by National Institute of Mental Health K23MH108823. SC was funded by National Institute on Deafness and other Communication Disorders (NIDCD/NIH) grant R01DC011277. CS was funded by the National Institute of Mental Health R01MH107741. MTwiNS was supported by the National Institute of Mental Health and the Offce of the Director National Institute of Health, under Award Number UG3MH114249 and the Eunice Kennedy Shriver National Institute of Child Health & Human Development of the National Institutes of Health under Award Number R01HD093334 to SAB and LWH. RJTM was funded by an ABC Talent Grant. The ABCD Study is supported by the National Institutes of Health (NIH) award numbers U01DA041022, U01DA041028, U01DA041048, U01DA041089, U01DA041106, U01DA041117, U01DA041120, U01DA041134, U01DA041148, U01DA041156, U01DA041174, U24DA041123, and U24DA041147.

---

**Appendix 0 Figure 4.**
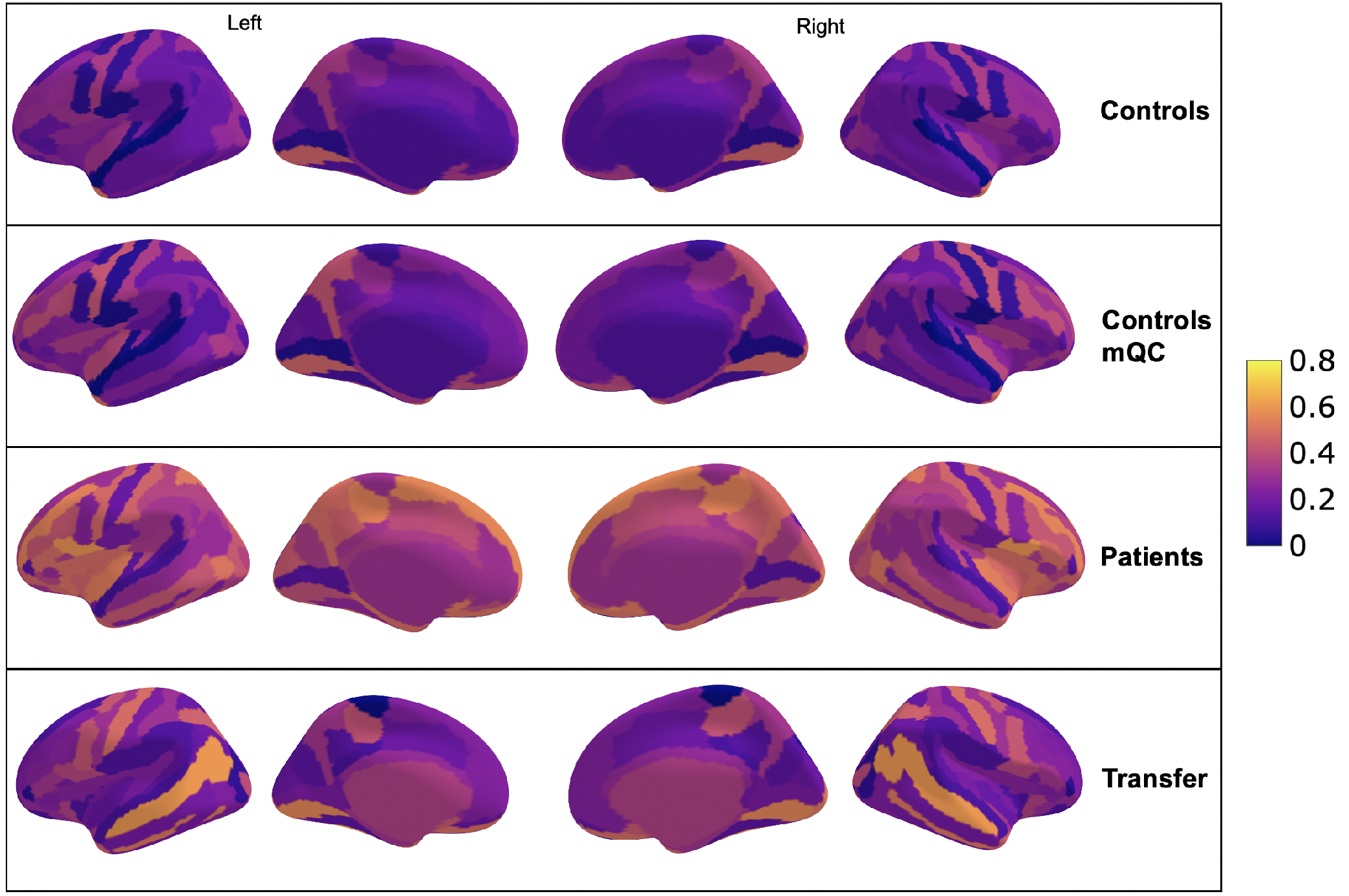
Comparison of the explained variance in cortical thickness across the different test sets. The patterns appear to be robust and consistent across the different test sets.

**Appendix 0 Table 2.**
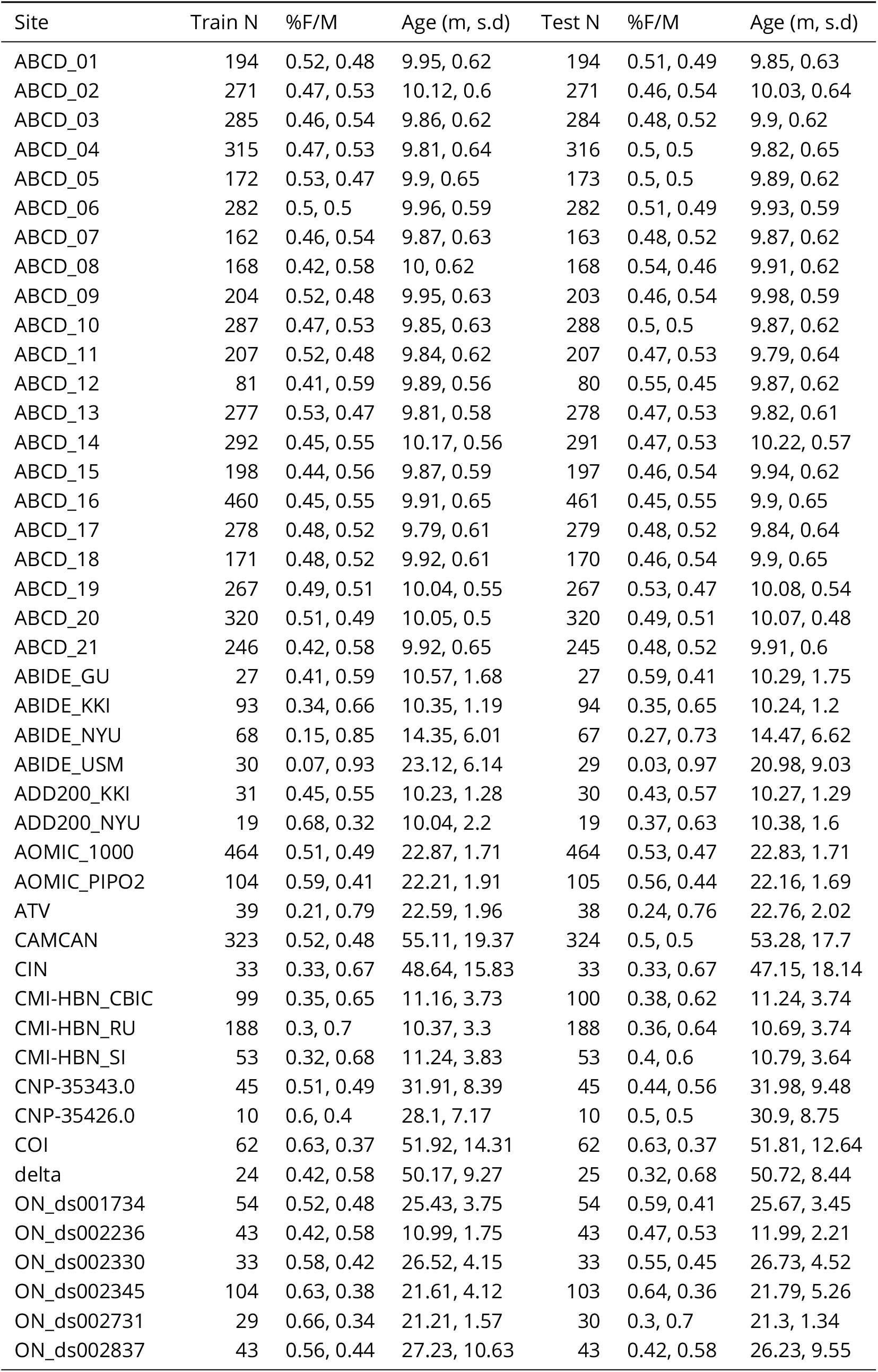
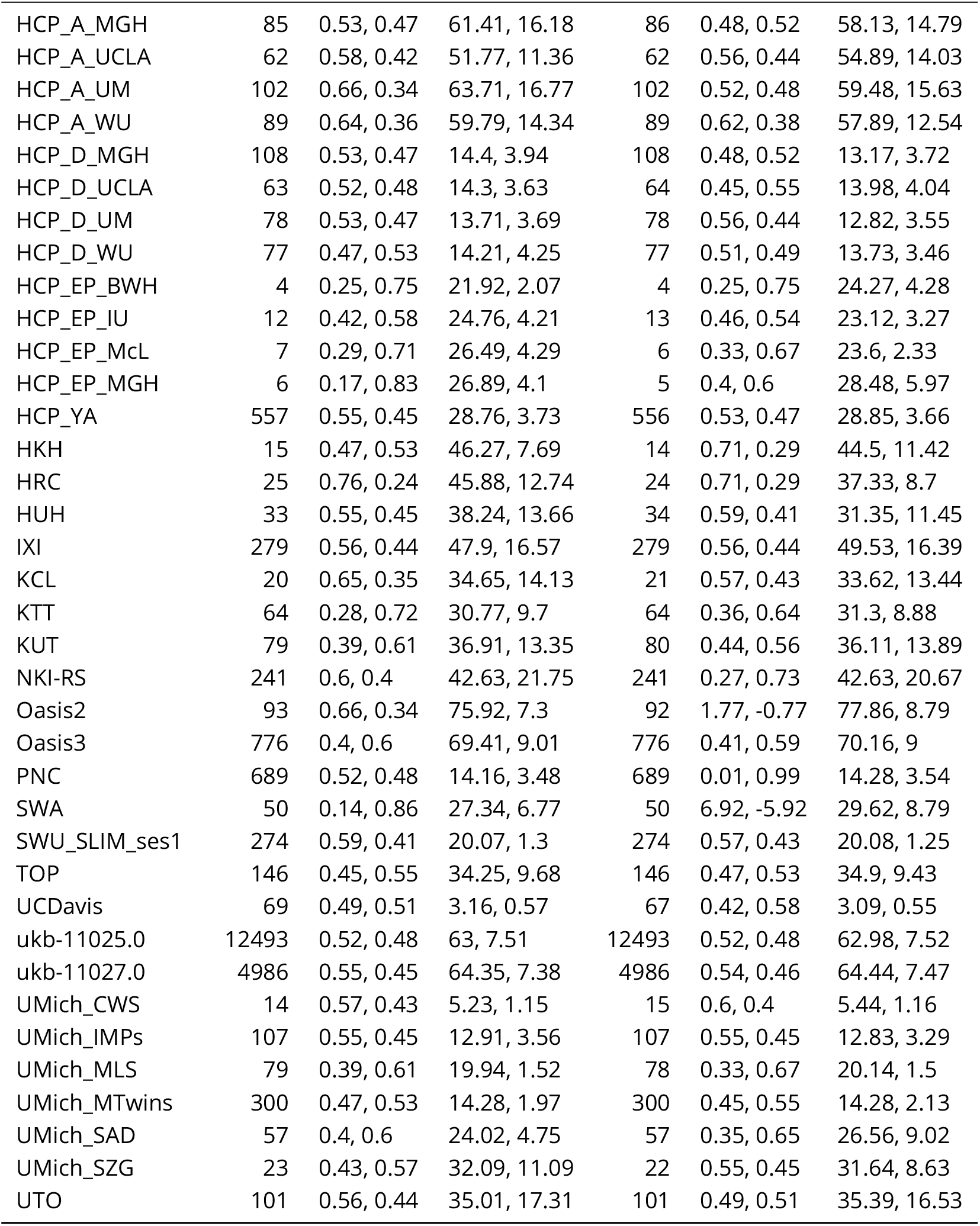
Full Sample Description, per site, in the train and test sets.

| Site | Train N | %F/M | Age (m, s.d) | Test N | %F/M | Age (m, s.d) |
| --- | --- | --- | --- | --- | --- | --- |
| ABCD_01 | 194 | 0.52, 0.48 | 9.95, 0.62 | 194 | 0.51, 0.49 | 9.85, 0.63 |
| ABCD_02 | 271 | 0.47, 0.53 | 10.12, 0.6 | 271 | 0.46, 0.54 | 10.03, 0.64 |
| ABCD_03 | 285 | 0.46, 0.54 | 9.86, 0.62 | 284 | 0.48, 0.52 | 9.9, 0.62 |
| ABCD_04 | 315 | 0.47, 0.53 | 9.81, 0.64 | 316 | 0.5, 0.5 | 9.82, 0.65 |
| ABCD_05 | 172 | 0.53, 0.47 | 9.9, 0.65 | 173 | 0.5, 0.5 | 9.89, 0.62 |
| ABCD_06 | 282 | 0.5, 0.5 | 9.96, 0.59 | 282 | 0.51, 0.49 | 9.93, 0.59 |
| ABCD_07 | 162 | 0.46, 0.54 | 9.87, 0.63 | 163 | 0.48, 0.52 | 9.87, 0.62 |
| ABCD_08 | 168 | 0.42, 0.58 | 10, 0.62 | 168 | 0.54, 0.46 | 9.91, 0.62 |
| ABCD_09 | 204 | 0.52, 0.48 | 9.95, 0.63 | 203 | 0.46, 0.54 | 9.98, 0.59 |
| ABCD_10 | 287 | 0.47, 0.53 | 9.85, 0.63 | 288 | 0.5, 0.5 | 9.87, 0.62 |
| ABCD_11 | 207 | 0.52, 0.48 | 9.84, 0.62 | 207 | 0.47, 0.53 | 9.79, 0.64 |
| ABCD_12 | 81 | 0.41, 0.59 | 9.89, 0.56 | 80 | 0.55, 0.45 | 9.87, 0.62 |
| ABCD_13 | 277 | 0.53, 0.47 | 9.81, 0.58 | 278 | 0.47, 0.53 | 9.82, 0.61 |
| ABCD_14 | 292 | 0.45, 0.55 | 10.17, 0.56 | 291 | 0.47, 0.53 | 10.22, 0.57 |
| ABCD_15 | 198 | 0.44, 0.56 | 9.87, 0.59 | 197 | 0.46, 0.54 | 9.94, 0.62 |
| ABCD_16 | 460 | 0.45, 0.55 | 9.91, 0.65 | 461 | 0.45, 0.55 | 9.9, 0.65 |
| ABCD_17 | 278 | 0.48, 0.52 | 9.79, 0.61 | 279 | 0.48, 0.52 | 9.84, 0.64 |
| ABCD_18 | 171 | 0.48, 0.52 | 9.92, 0.61 | 170 | 0.46, 0.54 | 9.9, 0.65 |
| ABCD_19 | 267 | 0.49, 0.51 | 10.04, 0.55 | 267 | 0.53, 0.47 | 10.08, 0.54 |
| ABCD_20 | 320 | 0.51, 0.49 | 10.05, 0.5 | 320 | 0.49, 0.51 | 10.07, 0.48 |
| ABCD_21 | 246 | 0.42, 0.58 | 9.92, 0.65 | 245 | 0.48, 0.52 | 9.91, 0.6 |
| ABIDE_GU | 27 | 0.41, 0.59 | 10.57, 1.68 | 27 | 0.59, 0.41 | 10.29, 1.75 |
| ABIDE_KKI | 93 | 0.34, 0.66 | 10.35, 1.19 | 94 | 0.35, 0.65 | 10.24, 1.2 |
| ABIDE_NYU | 68 | 0.15, 0.85 | 14.35, 6.01 | 67 | 0.27, 0.73 | 14.47, 6.62 |
| ABIDE_USM | 30 | 0.07, 0.93 | 23.12, 6.14 | 29 | 0.03, 0.97 | 20.98, 9.03 |
| ADD200_KKI | 31 | 0.45, 0.55 | 10.23, 1.28 | 30 | 0.43, 0.57 | 10.27, 1.29 |
| ADD200_NYU | 19 | 0.68, 0.32 | 10.04, 2.2 | 19 | 0.37, 0.63 | 10.38, 1.6 |
| AOMIC_1000 | 464 | 0.51, 0.49 | 22.87, 1.71 | 464 | 0.53, 0.47 | 22.83, 1.71 |
| AOMIC_PIPO2 | 104 | 0.59, 0.41 | 22.21, 1.91 | 105 | 0.56, 0.44 | 22.16, 1.69 |
| ATV | 39 | 0.21, 0.79 | 22.59, 1.96 | 38 | 0.24, 0.76 | 22.76, 2.02 |
| CAMCAN | 323 | 0.52, 0.48 | 55.11, 19.37 | 324 | 0.5, 0.5 | 53.28, 17.7 |
| CIN | 33 | 0.33, 0.67 | 48.64, 15.83 | 33 | 0.33, 0.67 | 47.15, 18.14 |
| CMI-HBN_CBIC | 99 | 0.35, 0.65 | 11.16, 3.73 | 100 | 0.38, 0.62 | 11.24, 3.74 |
| CMI-HBN_RU | 188 | 0.3, 0.7 | 10.37, 3.3 | 188 | 0.36, 0.64 | 10.69, 3.74 |
| CMI-HBN_SI | 53 | 0.32, 0.68 | 11.24, 3.83 | 53 | 0.4, 0.6 | 10.79, 3.64 |
| CNP-35343.0 | 45 | 0.51, 0.49 | 31.91, 8.39 | 45 | 0.44, 0.56 | 31.98, 9.48 |
| CNP-35426.0 | 10 | 0.6, 0.4 | 28.1, 7.17 | 10 | 0.5, 0.5 | 30.9, 8.75 |
| COI | 62 | 0.63, 0.37 | 51.92, 14.31 | 62 | 0.63, 0.37 | 51.81, 12.64 |
| delta | 24 | 0.42, 0.58 | 50.17, 9.27 | 25 | 0.32, 0.68 | 50.72, 8.44 |
| ON_ds001734 | 54 | 0.52, 0.48 | 25.43, 3.75 | 54 | 0.59, 0.41 | 25.67, 3.45 |
| ON_ds002236 | 43 | 0.42, 0.58 | 10.99, 1.75 | 43 | 0.47, 0.53 | 11.99, 2.21 |
| ON_ds002330 | 33 | 0.58, 0.42 | 26.52, 4.15 | 33 | 0.55, 0.45 | 26.73, 4.52 |
| ON_ds002345 | 104 | 0.63, 0.38 | 21.61, 4.12 | 103 | 0.64, 0.36 | 21.79, 5.26 |
| ON_ds002731 | 29 | 0.66, 0.34 | 21.21, 1.57 | 30 | 0.3, 0.7 | 21.3, 1.34 |
| ON_ds002837 | 43 | 0.56, 0.44 | 27.23, 10.63 | 43 | 0.42, 0.58 | 26.23, 9.55 |
| HCP_A_MGH | 85 | 0.53, 0.47 | 61.41, 16.18 | 86 | 0.48, 0.52 | 58.13, 14.79 |
| HCP_A_UCLA | 62 | 0.58, 0.42 | 51.77, 11.36 | 62 | 0.56, 0.44 | 54.89, 14.03 |
| HCP_A_UM | 102 | 0.66, 0.34 | 63.71, 16.77 | 102 | 0.52, 0.48 | 59.48, 15.63 |
| HCP_A_WU | 89 | 0.64, 0.36 | 59.79, 14.34 | 89 | 0.62, 0.38 | 57.89, 12.54 |
| HCP_D_MGH | 108 | 0.53, 0.47 | 14.4, 3.94 | 108 | 0.48, 0.52 | 13.17, 3.72 |
| HCP_D_UCLA | 63 | 0.52, 0.48 | 14.3, 3.63 | 64 | 0.45, 0.55 | 13.98, 4.04 |
| HCP_D_UM | 78 | 0.53, 0.47 | 13.71, 3.69 | 78 | 0.56, 0.44 | 12.82, 3.55 |
| HCP_D_WU | 77 | 0.47, 0.53 | 14.21, 4.25 | 77 | 0.51, 0.49 | 13.73, 3.46 |
| HCP_EP_BWH | 4 | 0.25, 0.75 | 21.92, 2.07 | 4 | 0.25, 0.75 | 24.27, 4.28 |
| HCP_EP_IU | 12 | 0.42, 0.58 | 24.76, 4.21 | 13 | 0.46, 0.54 | 23.12, 3.27 |
| HCP_EP_McL | 7 | 0.29, 0.71 | 26.49, 4.29 | 6 | 0.33, 0.67 | 23.6, 2.33 |
| HCP_EP_MGH | 6 | 0.17, 0.83 | 26.89, 4.1 | 5 | 0.4, 0.6 | 28.48, 5.97 |
| HCP_YA | 557 | 0.55, 0.45 | 28.76, 3.73 | 556 | 0.53, 0.47 | 28.85, 3.66 |
| HKH | 15 | 0.47, 0.53 | 46.27, 7.69 | 14 | 0.71, 0.29 | 44.5, 11.42 |
| HRC | 25 | 0.76, 0.24 | 45.88, 12.74 | 24 | 0.71, 0.29 | 37.33, 8.7 |
| HUH | 33 | 0.55, 0.45 | 38.24, 13.66 | 34 | 0.59, 0.41 | 31.35, 11.45 |
| IXI | 279 | 0.56, 0.44 | 47.9, 16.57 | 279 | 0.56, 0.44 | 49.53, 16.39 |
| KCL | 20 | 0.65, 0.35 | 34.65, 14.13 | 21 | 0.57, 0.43 | 33.62, 13.44 |
| KTT | 64 | 0.28, 0.72 | 30.77, 9.7 | 64 | 0.36, 0.64 | 31.3, 8.88 |
| KUT | 79 | 0.39, 0.61 | 36.91, 13.35 | 80 | 0.44, 0.56 | 36.11, 13.89 |
| NKI-RS | 241 | 0.6, 0.4 | 42.63, 21.75 | 241 | 0.27, 0.73 | 42.63, 20.67 |
| Oasis2 | 93 | 0.66, 0.34 | 75.92, 7.3 | 92 | 1.77, -0.77 | 77.86, 8.79 |
| Oasis3 | 776 | 0.4, 0.6 | 69.41, 9.01 | 776 | 0.41, 0.59 | 70.16, 9 |
| PNC | 689 | 0.52, 0.48 | 14.16, 3.48 | 689 | 0.01, 0.99 | 14.28, 3.54 |
| SWA | 50 | 0.14, 0.86 | 27.34, 6.77 | 50 | 6.92, -5.92 | 29.62, 8.79 |
| SWU_SLIM_ses1 | 274 | 0.59, 0.41 | 20.07, 1.3 | 274 | 0.57, 0.43 | 20.08, 1.25 |
| TOP | 146 | 0.45, 0.55 | 34.25, 9.68 | 146 | 0.47, 0.53 | 34.9, 9.43 |
| UCDavis | 69 | 0.49, 0.51 | 3.16, 0.57 | 67 | 0.42, 0.58 | 3.09, 0.55 |
| ukb-11025.0 | 12493 | 0.52, 0.48 | 63, 7.51 | 12493 | 0.52, 0.48 | 62.98, 7.52 |
| ukb-11027.0 | 4986 | 0.55, 0.45 | 64.35, 7.38 | 4986 | 0.54, 0.46 | 64.44, 7.47 |
| UMich_CWS | 14 | 0.57, 0.43 | 5.23, 1.15 | 15 | 0.6, 0.4 | 5.44, 1.16 |
| UMich_IMPs | 107 | 0.55, 0.45 | 12.91, 3.56 | 107 | 0.55, 0.45 | 12.83, 3.29 |
| UMich_MLS | 79 | 0.39, 0.61 | 19.94, 1.52 | 78 | 0.33, 0.67 | 20.14, 1.5 |
| UMich_MTwins | 300 | 0.47, 0.53 | 14.28, 1.97 | 300 | 0.45, 0.55 | 14.28, 2.13 |
| UMich_SAD | 57 | 0.4, 0.6 | 24.02, 4.75 | 57 | 0.35, 0.65 | 26.56, 9.02 |
| UMich_SZG | 23 | 0.43, 0.57 | 32.09, 11.09 | 22 | 0.55, 0.45 | 31.64, 8.63 |
| UTO | 101 | 0.56, 0.44 | 35.01, 17.31 | 101 | 0.49, 0.51 | 35.39, 16.53 |

**Appendix 0 Table 3.**
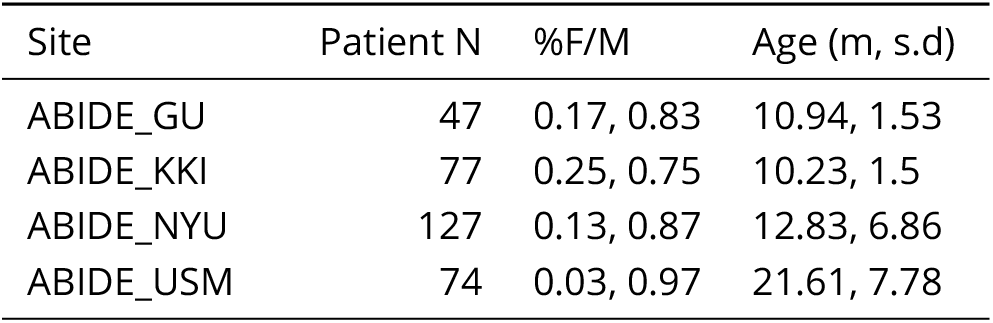
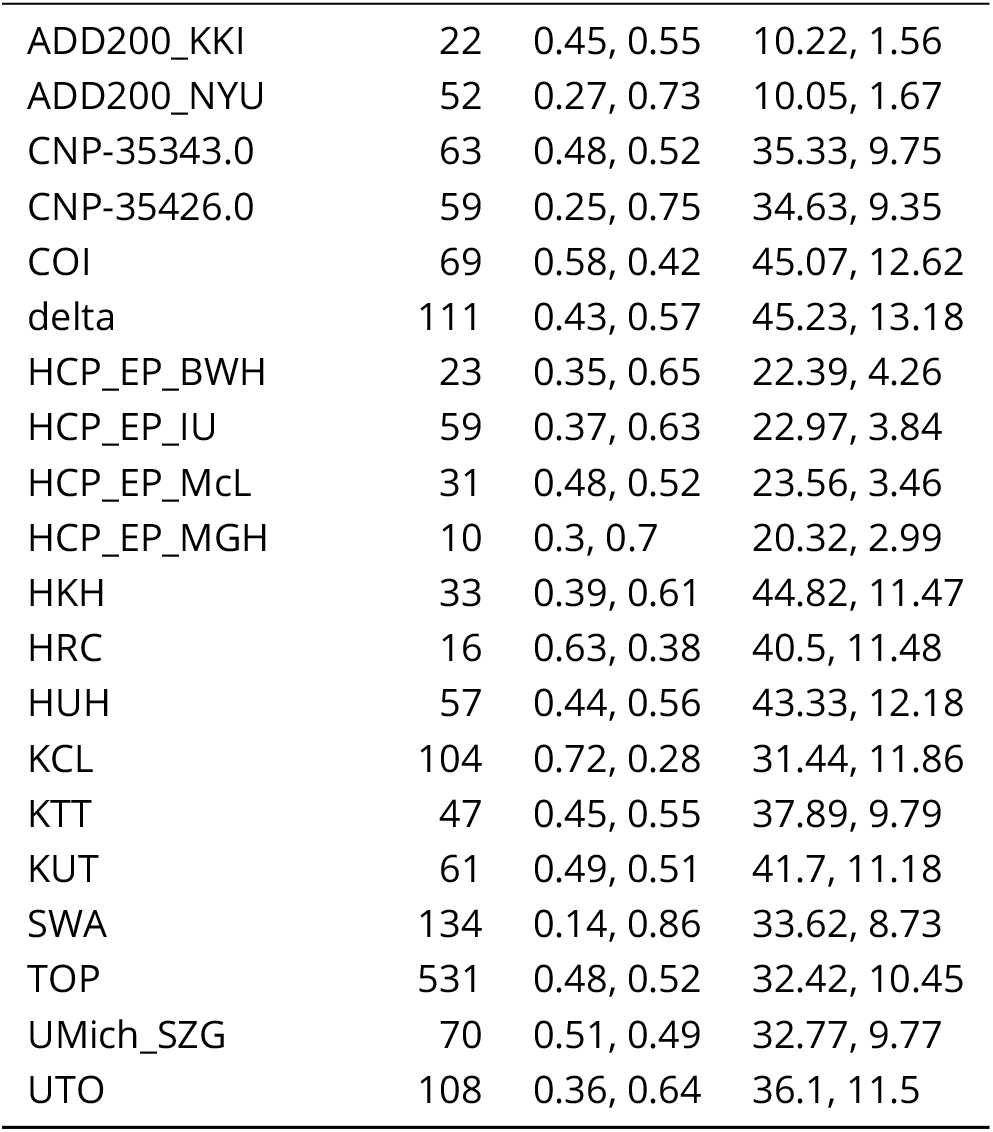
Clinical Sample Description

**Appendix 0 Table 4.**
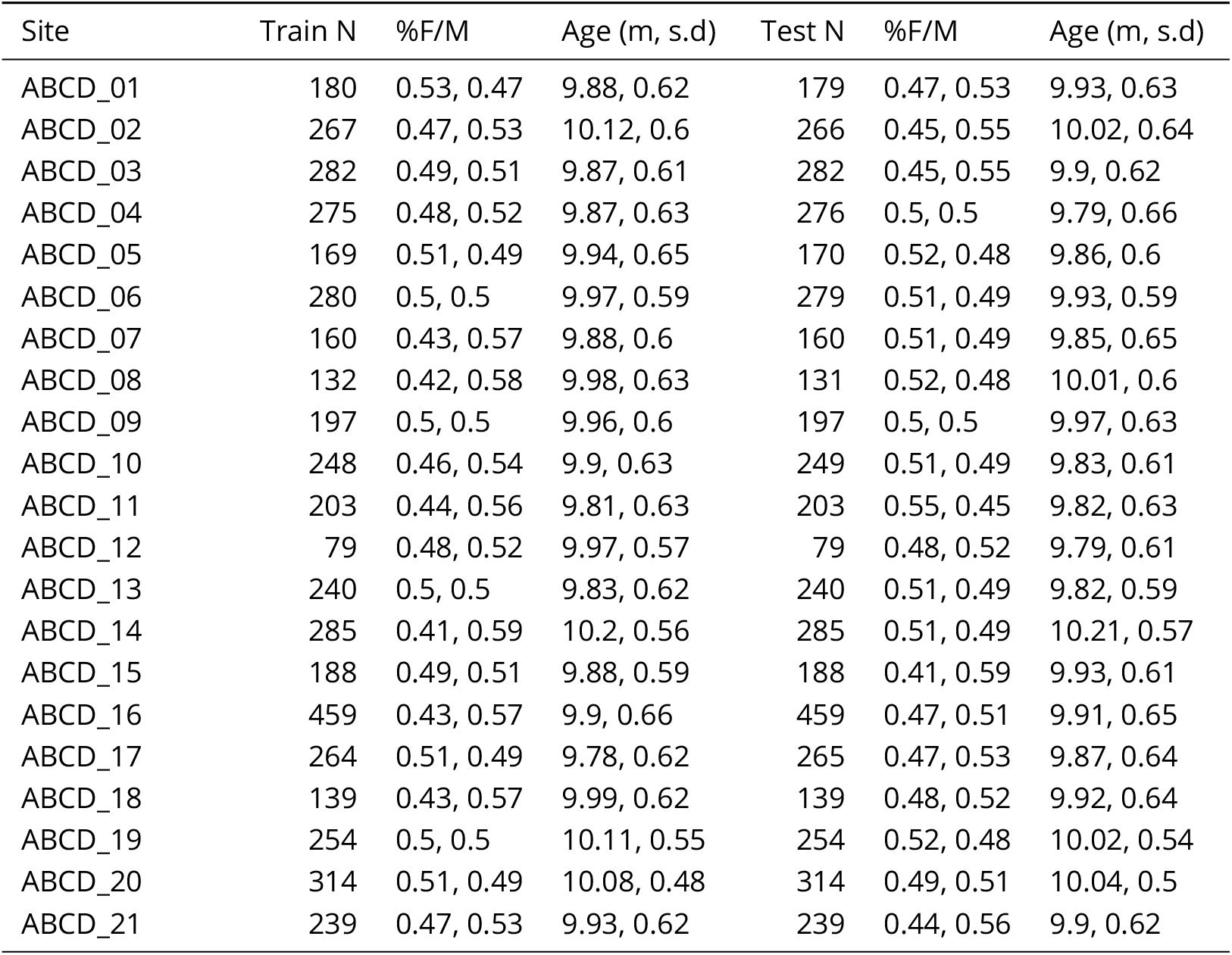
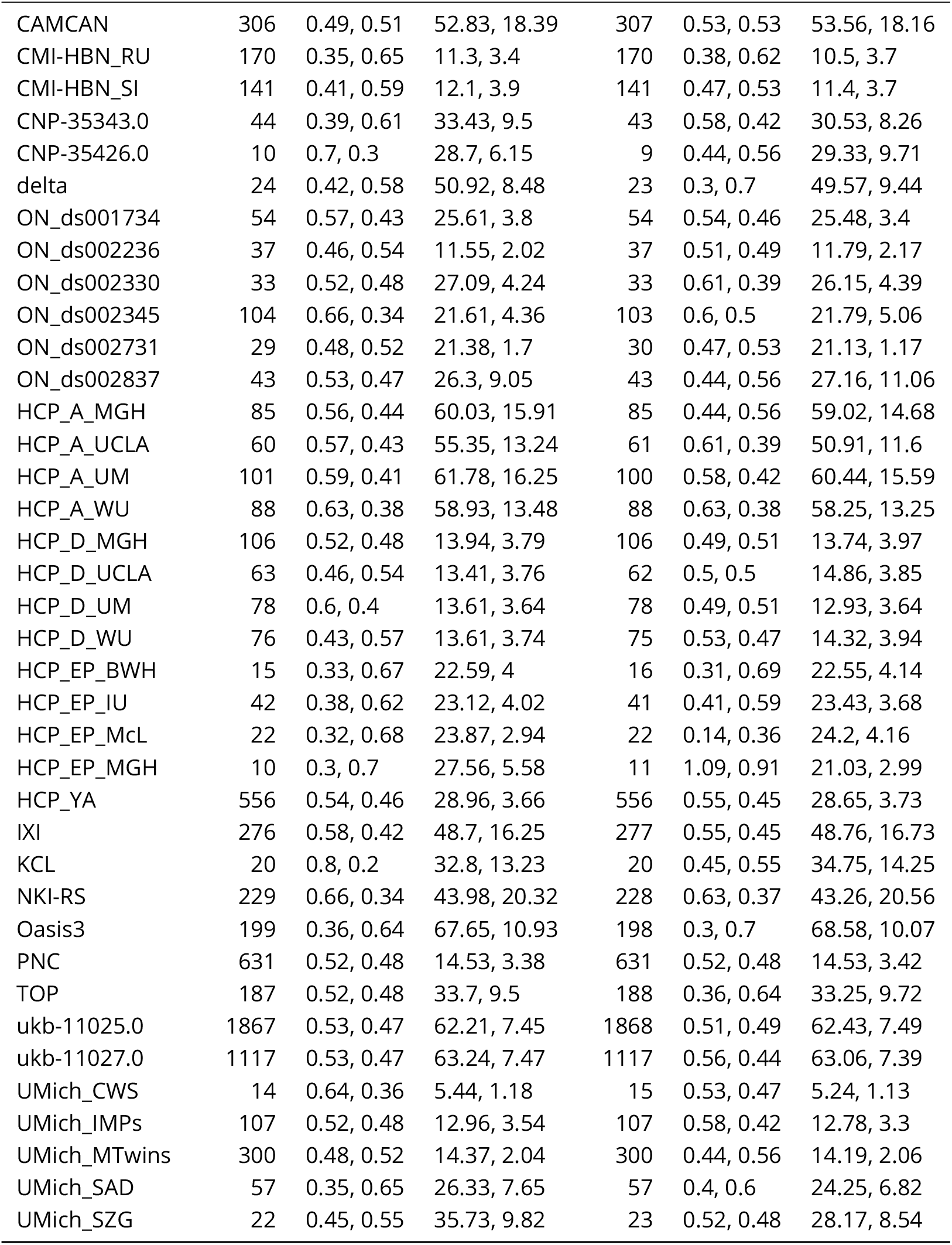
mQC Sample Description, per site, in the train and test sets.

**Appendix 0 Table 5.**
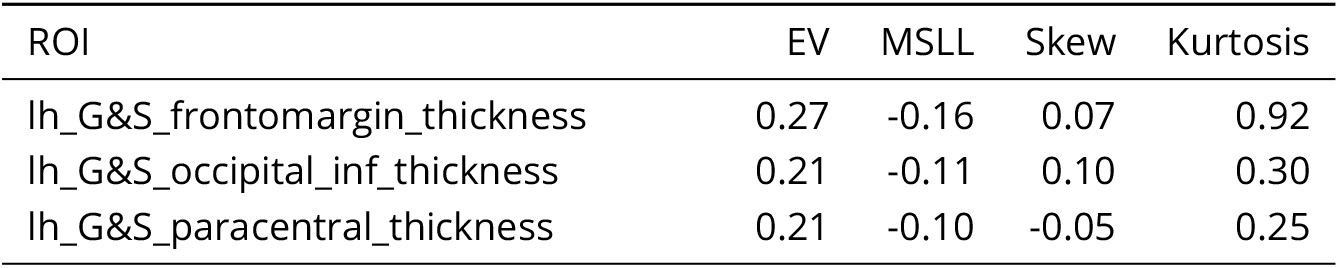
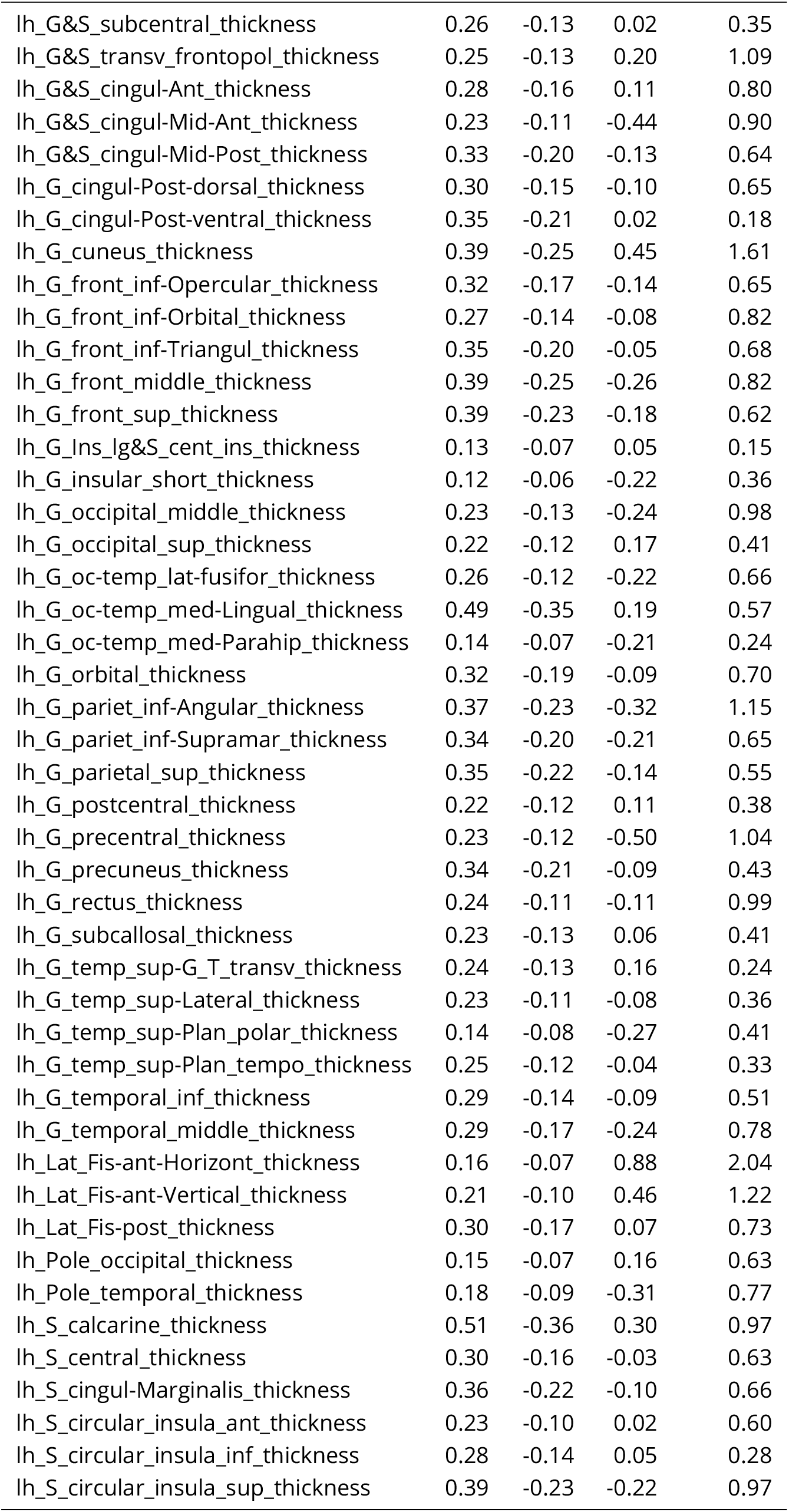
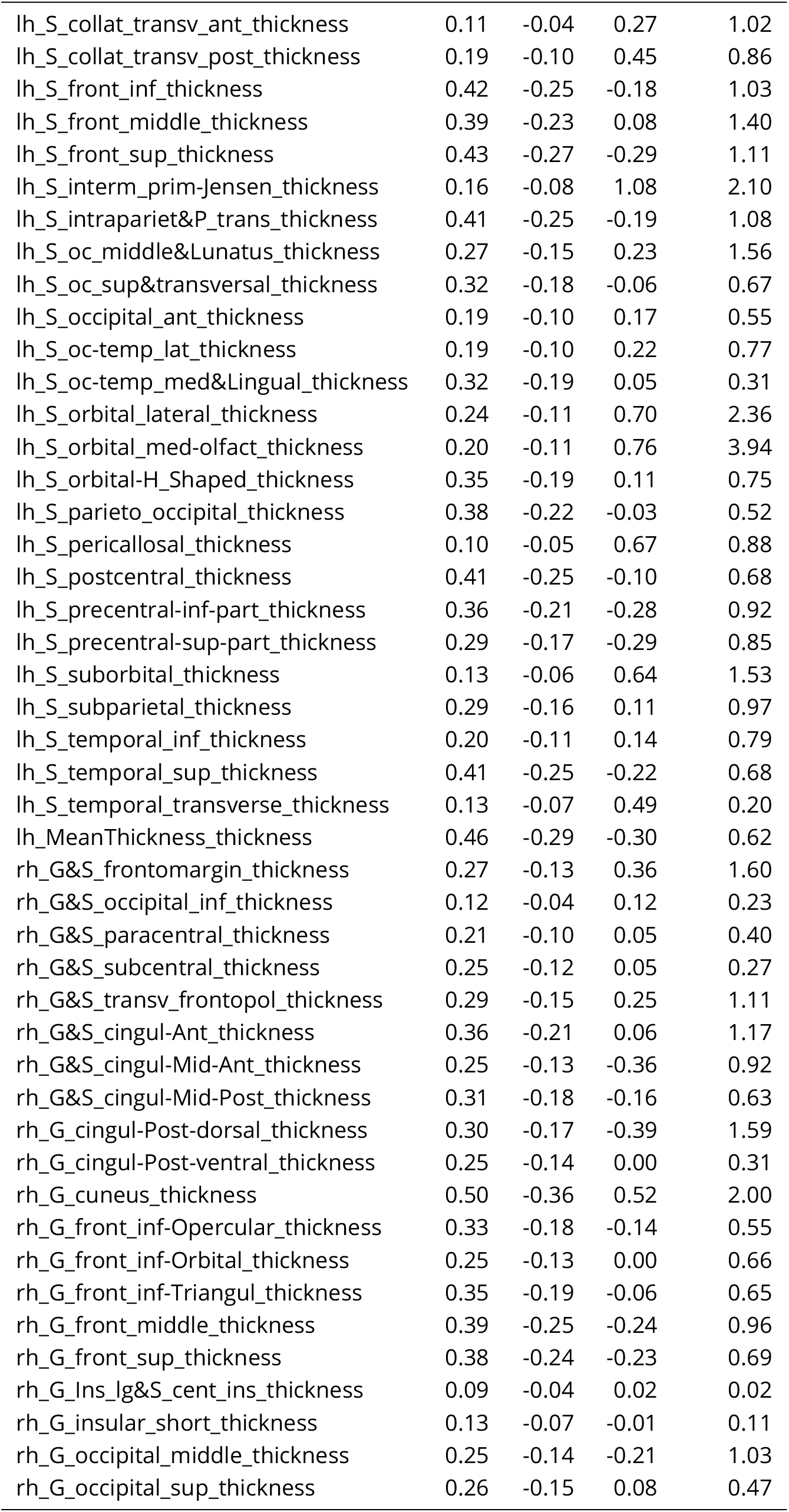
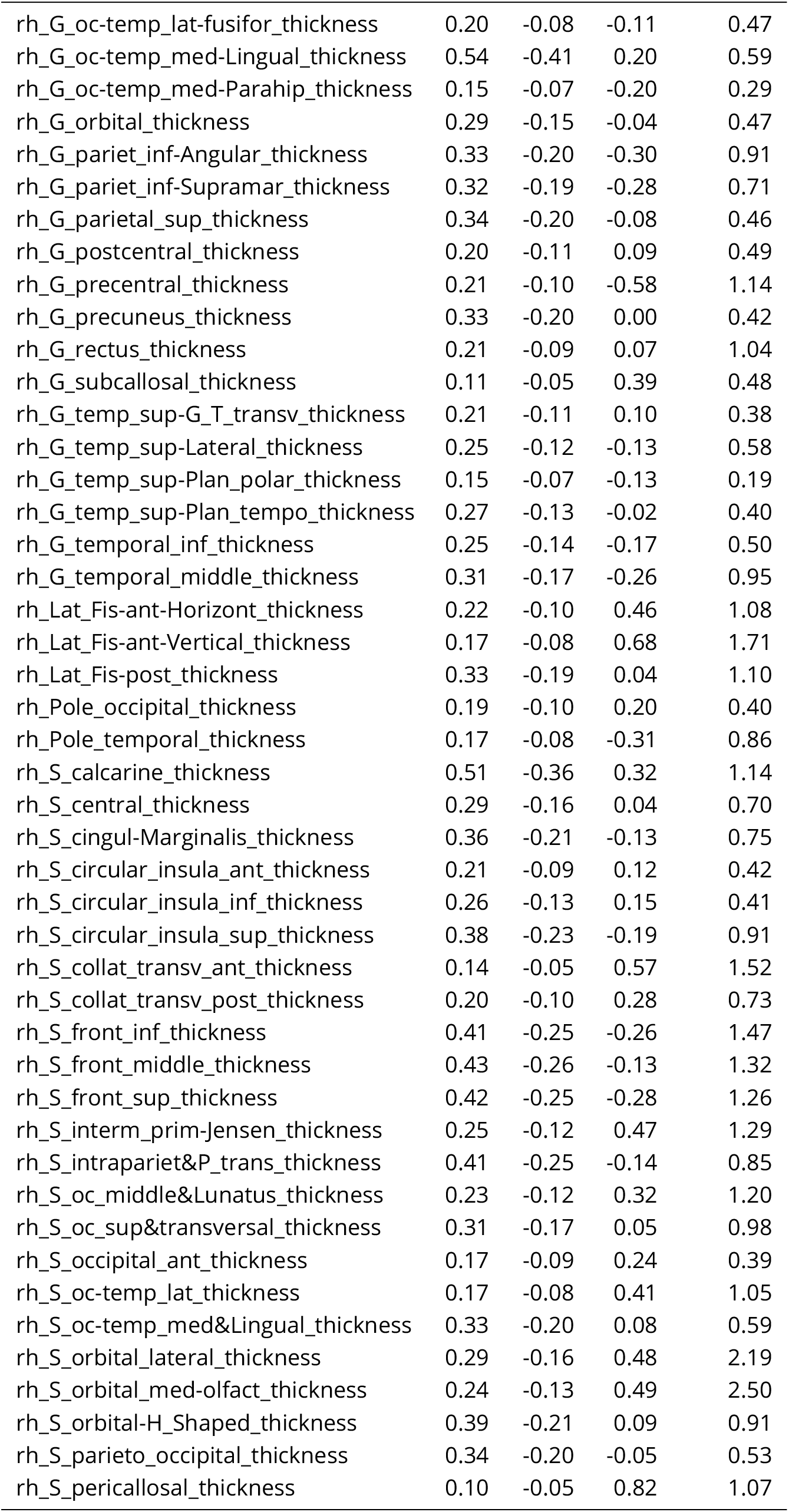
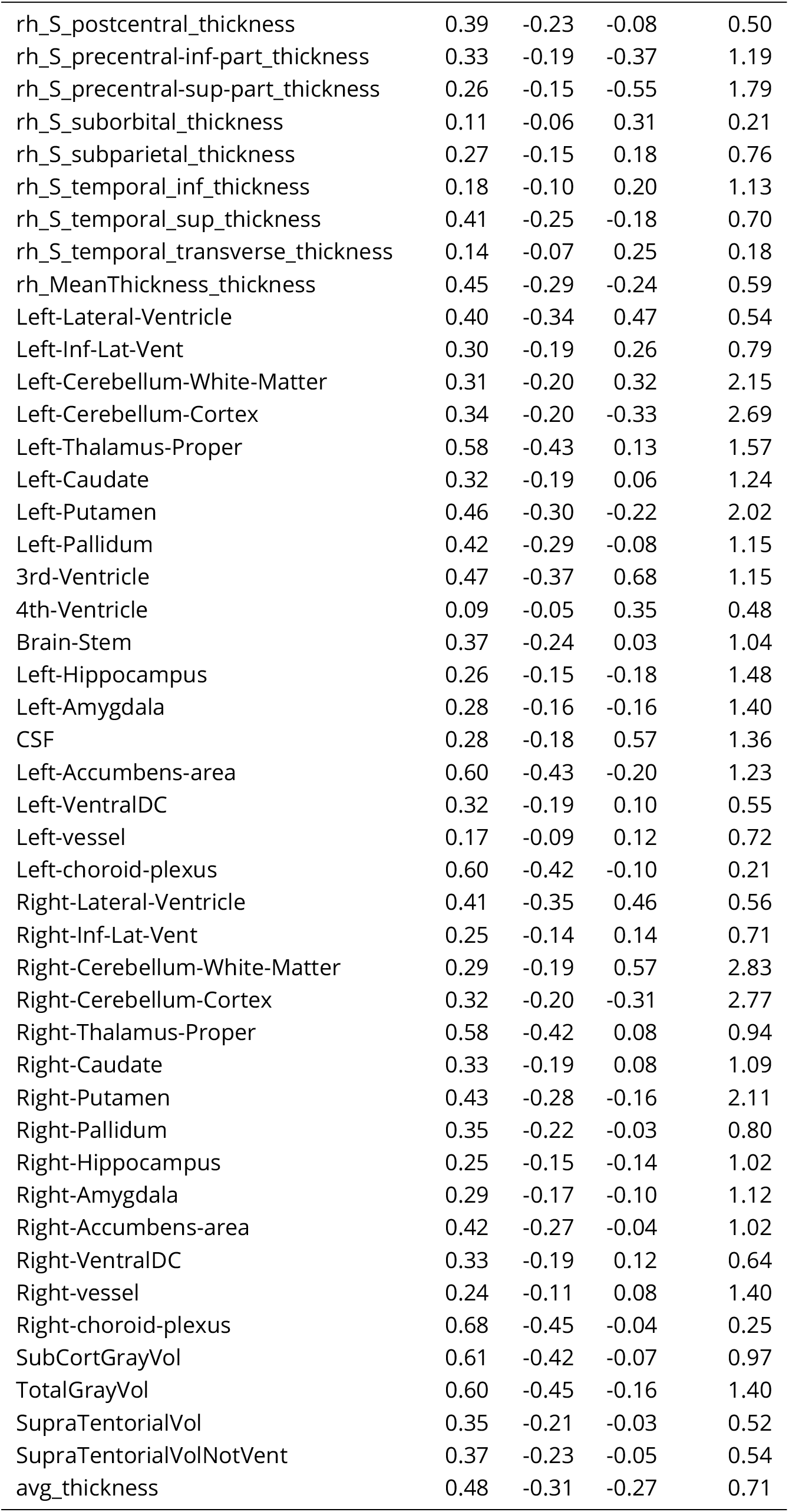
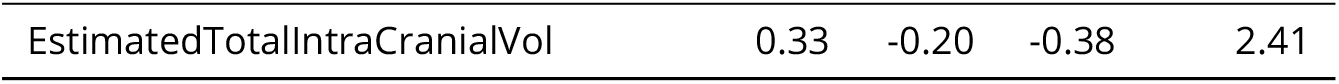
Full Test Set Evaluation Metrics

| ROI | EV | MSLL | Skew | Kurtosis |
| --- | --- | --- | --- | --- |
| lh_G&S_frontomargin_thickness | 0.27 | -0.16 | 0.07 | 0.92 |
| lh_G&S_occipital_inf_thickness | 0.21 | -0.11 | 0.10 | 0.30 |
| lh_G&S_paracentral_thickness | 0.21 | -0.10 | -0.05 | 0.25 |
| Continued on next page |  |  |  |  |

**Appendix 0 Table 5.** Full Test Set Evaluation Metrics
| ROI | EV | MSLL | Skew | Kurtosis |
| --- | --- | --- | --- | --- |
| lh_G&S_subcentral_thickness | 0.26 | -0.13 | 0.02 | 0.35 |
| lh_G&S_transv_frontopol_thickness | 0.25 | -0.13 | 0.20 | 1.09 |
| lh_G&S_cingul-Ant_thickness | 0.28 | -0.16 | 0.11 | 0.80 |
| lh_G&S_cingul-Mid-Ant_thickness | 0.23 | -0.11 | -0.44 | 0.90 |
| lh_G&S_cingul-Mid-Post_thickness | 0.33 | -0.20 | -0.13 | 0.64 |
| lh_G_cingul-Post-dorsal_thickness | 0.30 | -0.15 | -0.10 | 0.65 |
| lh_G_cingul-Post-ventral_thickness | 0.35 | -0.21 | 0.02 | 0.18 |
| lh_G_cuneus_thickness | 0.39 | -0.25 | 0.45 | 1.61 |
| lh_G_front_inf-Opercular_thickness | 0.32 | -0.17 | -0.14 | 0.65 |
| lh_G_front_inf-Orbital_thickness | 0.27 | -0.14 | -0.08 | 0.82 |
| lh_G_front_inf-Triangul_thickness | 0.35 | -0.20 | -0.05 | 0.68 |
| lh_G_front_middle_thickness | 0.39 | -0.25 | -0.26 | 0.82 |
| lh_G_front_sup_thickness | 0.39 | -0.23 | -0.18 | 0.62 |
| lh_G_Ins_Ig&S_cent_ins_thickness | 0.13 | -0.07 | 0.05 | 0.15 |
| lh_G_insular_short_thickness | 0.12 | -0.06 | -0.22 | 0.36 |
| lh_G_occipital_middle_thickness | 0.23 | -0.13 | -0.24 | 0.98 |
| lh_G_occipital_sup_thickness | 0.22 | -0.12 | 0.17 | 0.41 |
| lh_G_oc-temp_lat-fusifor_thickness | 0.26 | -0.12 | -0.22 | 0.66 |
| lh_G_oc-temp_med-Lingual_thickness | 0.49 | -0.35 | 0.19 | 0.57 |
| lh_G_oc-temp_med-Parahip_thickness | 0.14 | -0.07 | -0.21 | 0.24 |
| lh_G_orbital_thickness | 0.32 | -0.19 | -0.09 | 0.70 |
| lh_G_pariet_inf-Angular_thickness | 0.37 | -0.23 | -0.32 | 1.15 |
| lh_G_pariet_inf-Supramar_thickness | 0.34 | -0.20 | -0.21 | 0.65 |
| lh_G_parietal_sup_thickness | 0.35 | -0.22 | -0.14 | 0.55 |
| lh_G_postcentral_thickness | 0.22 | -0.12 | 0.11 | 0.38 |
| lh_G_precentral_thickness | 0.23 | -0.12 | -0.50 | 1.04 |
| lh_G_precuneus_thickness | 0.34 | -0.21 | -0.09 | 0.43 |
| lh_G_rectus_thickness | 0.24 | -0.11 | -0.11 | 0.99 |
| lh_G_subcallosal_thickness | 0.23 | -0.13 | 0.06 | 0.41 |
| lh_G_temp_sup-G_T_transv_thickness | 0.24 | -0.13 | 0.16 | 0.24 |
| lh_G_temp_sup-Lateral_thickness | 0.23 | -0.11 | -0.08 | 0.36 |
| lh_G_temp_sup-Plan_polar_thickness | 0.14 | -0.08 | -0.27 | 0.41 |
| lh_G_temp_sup-Plan_tempo_thickness | 0.25 | -0.12 | -0.04 | 0.33 |
| lh_G_temporal_inf_thickness | 0.29 | -0.14 | -0.09 | 0.51 |
| lh_G_temporal_middle_thickness | 0.29 | -0.17 | -0.24 | 0.78 |
| lh_Lat_Fis-ant-Horizont_thickness | 0.16 | -0.07 | 0.88 | 2.04 |
| lh_Lat_Fis-ant-Vertical_thickness | 0.21 | -0.10 | 0.46 | 1.22 |
| lh_Lat_Fis-post_thickness | 0.30 | -0.17 | 0.07 | 0.73 |
| lh_Pole_occipital_thickness | 0.15 | -0.07 | 0.16 | 0.63 |
| lh_Pole_temporal_thickness | 0.18 | -0.09 | -0.31 | 0.77 |
| lh_S_calcarine_thickness | 0.51 | -0.36 | 0.30 | 0.97 |
| lh_S_central_thickness | 0.30 | -0.16 | -0.03 | 0.63 |
| lh_S_cingul-Marginalis_thickness | 0.36 | -0.22 | -0.10 | 0.66 |
| lh_S_circular_insula_ant_thickness | 0.23 | -0.10 | 0.02 | 0.60 |
| lh_S_circular_insula_inf_thickness | 0.28 | -0.14 | 0.05 | 0.28 |
| lh_S_circular_insula_sup_thickness | 0.39 | -0.23 | -0.22 | 0.97 |
| lh_S_collat_transv_ant_thickness | 0.11 | -0.04 | 0.27 | 1.02 |
| lh_S_collat_transv_post_thickness | 0.19 | -0.10 | 0.45 | 0.86 |
| lh_S_front_inf_thickness | 0.42 | -0.25 | -0.18 | 1.03 |
| lh_S_front_middle_thickness | 0.39 | -0.23 | 0.08 | 1.40 |
| lh_S_front_sup_thickness | 0.43 | -0.27 | -0.29 | 1.11 |
| lh_S_interm_prim-Jensen_thickness | 0.16 | -0.08 | 1.08 | 2.10 |
| lh_S_intrapariet&P_trans_thickness | 0.41 | -0.25 | -0.19 | 1.08 |
| lh_S_oc_middle&Lunatus_thickness | 0.27 | -0.15 | 0.23 | 1.56 |
| lh_S_oc_sup&transversal_thickness | 0.32 | -0.18 | -0.06 | 0.67 |
| lh_S_occipital_ant_thickness | 0.19 | -0.10 | 0.17 | 0.55 |
| lh_S_oc-temp_lat_thickness | 0.19 | -0.10 | 0.22 | 0.77 |
| lh_S_oc-temp_med&Lingual_thickness | 0.32 | -0.19 | 0.05 | 0.31 |
| lh_S_orbital_lateral_thickness | 0.24 | -0.11 | 0.70 | 2.36 |
| lh_S_orbital_med-olfact_thickness | 0.20 | -0.11 | 0.76 | 3.94 |
| lh_S_orbital-H_Shaped_thickness | 0.35 | -0.19 | 0.11 | 0.75 |
| lh_S_parieto_occipital_thickness | 0.38 | -0.22 | -0.03 | 0.52 |
| lh_S_pericallosal_thickness | 0.10 | -0.05 | 0.67 | 0.88 |
| lh_S_postcentral_thickness | 0.41 | -0.25 | -0.10 | 0.68 |
| lh_S_precentral-inf-part_thickness | 0.36 | -0.21 | -0.28 | 0.92 |
| lh_S_precentral-sup-part_thickness | 0.29 | -0.17 | -0.29 | 0.85 |
| lh_S_suborbital_thickness | 0.13 | -0.06 | 0.64 | 1.53 |
| lh_S_subparietal_thickness | 0.29 | -0.16 | 0.11 | 0.97 |
| lh_S_temporal_inf_thickness | 0.20 | -0.11 | 0.14 | 0.79 |
| lh_S_temporal_sup_thickness | 0.41 | -0.25 | -0.22 | 0.68 |
| lh_S_temporal_transverse_thickness | 0.13 | -0.07 | 0.49 | 0.20 |
| lh_MeanThickness_thickness | 0.46 | -0.29 | -0.30 | 0.62 |
| rh_G&S_frontomargin_thickness | 0.27 | -0.13 | 0.36 | 1.60 |
| rh_G&S_occipital_inf_thickness | 0.12 | -0.04 | 0.12 | 0.23 |
| rh_G&S_paracentral_thickness | 0.21 | -0.10 | 0.05 | 0.40 |
| rh_G&S_subcentral_thickness | 0.25 | -0.12 | 0.05 | 0.27 |
| rh_G&S_transv_frontopol_thickness | 0.29 | -0.15 | 0.25 | 1.11 |
| rh_G&S_cingul-Ant_thickness | 0.36 | -0.21 | 0.06 | 1.17 |
| rh_G&S_cingul-Mid-Ant_thickness | 0.25 | -0.13 | -0.36 | 0.92 |
| rh_G&S_cingul-Mid-Post_thickness | 0.31 | -0.18 | -0.16 | 0.63 |
| rh_G_cingul-Post-dorsal_thickness | 0.30 | -0.17 | -0.39 | 1.59 |
| rh_G_cingul-Post-ventral_thickness | 0.25 | -0.14 | 0.00 | 0.31 |
| rh_G_cuneus_thickness | 0.50 | -0.36 | 0.52 | 2.00 |
| rh_G_front_inf-Opercular_thickness | 0.33 | -0.18 | -0.14 | 0.55 |
| rh_G_front_inf-Orbital_thickness | 0.25 | -0.13 | 0.00 | 0.66 |
| rh_G_front_inf-Triangul_thickness | 0.35 | -0.19 | -0.06 | 0.65 |
| rh_G_front_middle_thickness | 0.39 | -0.25 | -0.24 | 0.96 |
| rh_G_front_sup_thickness | 0.38 | -0.24 | -0.23 | 0.69 |
| rh_G_Ins_Ig&S_cent_ins_thickness | 0.09 | -0.04 | 0.02 | 0.02 |
| rh_G_insular_short_thickness | 0.13 | -0.07 | -0.01 | 0.11 |
| rh_G_occipital_middle_thickness | 0.25 | -0.14 | -0.21 | 1.03 |
| rh_G_occipital_sup_thickness | 0.26 | -0.15 | 0.08 | 0.47 |
| rh_G_oc-temp_lat-fusifor_thickness | 0.20 | -0.08 | -0.11 | 0.47 |
| rh_G_oc-temp_med-Lingual_thickness | 0.54 | -0.41 | 0.20 | 0.59 |
| rh_G_oc-temp_med-Parahip_thickness | 0.15 | -0.07 | -0.20 | 0.29 |
| rh_G_orbital_thickness | 0.29 | -0.15 | -0.04 | 0.47 |
| rh_G_pariet_inf-Angular_thickness | 0.33 | -0.20 | -0.30 | 0.91 |
| rh_G_pariet_inf-Supramar_thickness | 0.32 | -0.19 | -0.28 | 0.71 |
| rh_G_parietal_sup_thickness | 0.34 | -0.20 | -0.08 | 0.46 |
| rh_G_postcentral_thickness | 0.20 | -0.11 | 0.09 | 0.49 |
| rh_G_precentral_thickness | 0.21 | -0.10 | -0.58 | 1.14 |
| rh_G_precuneus_thickness | 0.33 | -0.20 | 0.00 | 0.42 |
| rh_G_rectus_thickness | 0.21 | -0.09 | 0.07 | 1.04 |
| rh_G_subcallosal_thickness | 0.11 | -0.05 | 0.39 | 0.48 |
| rh_G_temp_sup-G_T_transv_thickness | 0.21 | -0.11 | 0.10 | 0.38 |
| rh_G_temp_sup-Lateral_thickness | 0.25 | -0.12 | -0.13 | 0.58 |
| rh_G_temp_sup-Plan_polar_thickness | 0.15 | -0.07 | -0.13 | 0.19 |
| rh_G_temp_sup-Plan_tempo_thickness | 0.27 | -0.13 | -0.02 | 0.40 |
| rh_G_temporal_inf_thickness | 0.25 | -0.14 | -0.17 | 0.50 |
| rh_G_temporal_middle_thickness | 0.31 | -0.17 | -0.26 | 0.95 |
| rh_Lat_Fis-ant-Horizont_thickness | 0.22 | -0.10 | 0.46 | 1.08 |
| rh_Lat_Fis-ant-Vertical_thickness | 0.17 | -0.08 | 0.68 | 1.71 |
| rh_Lat_Fis-post_thickness | 0.33 | -0.19 | 0.04 | 1.10 |
| rh_Pole_occipital_thickness | 0.19 | -0.10 | 0.20 | 0.40 |
| rh_Pole_temporal_thickness | 0.17 | -0.08 | -0.31 | 0.86 |
| rh_S_calcarine_thickness | 0.51 | -0.36 | 0.32 | 1.14 |
| rh_S_central_thickness | 0.29 | -0.16 | 0.04 | 0.70 |
| rh_S_cingul-Marginalis_thickness | 0.36 | -0.21 | -0.13 | 0.75 |
| rh_S_circular_insula_ant_thickness | 0.21 | -0.09 | 0.12 | 0.42 |
| rh_S_circular_insula_inf_thickness | 0.26 | -0.13 | 0.15 | 0.41 |
| rh_S_circular_insula_sup_thickness | 0.38 | -0.23 | -0.19 | 0.91 |
| rh_S_collat_transv_ant_thickness | 0.14 | -0.05 | 0.57 | 1.52 |
| rh_S_collat_transv_post_thickness | 0.20 | -0.10 | 0.28 | 0.73 |
| rh_S_front_inf_thickness | 0.41 | -0.25 | -0.26 | 1.47 |
| rh_S_front_middle_thickness | 0.43 | -0.26 | -0.13 | 1.32 |
| rh_S_front_sup_thickness | 0.42 | -0.25 | -0.28 | 1.26 |
| rh_S_interm_prim-Jensen_thickness | 0.25 | -0.12 | 0.47 | 1.29 |
| rh_S_intrapariet&P_trans_thickness | 0.41 | -0.25 | -0.14 | 0.85 |
| rh_S_oc_middle&Lunatus_thickness | 0.23 | -0.12 | 0.32 | 1.20 |
| rh_S_oc_sup&transversal_thickness | 0.31 | -0.17 | 0.05 | 0.98 |
| rh_S_occipital_ant_thickness | 0.17 | -0.09 | 0.24 | 0.39 |
| rh_S_oc-temp_lat_thickness | 0.17 | -0.08 | 0.41 | 1.05 |
| rh_S_oc-temp_med&Lingual_thickness | 0.33 | -0.20 | 0.08 | 0.59 |
| rh_S_orbital_lateral_thickness | 0.29 | -0.16 | 0.48 | 2.19 |
| rh_S_orbital_med-olfact_thickness | 0.24 | -0.13 | 0.49 | 2.50 |
| rh_S_orbital-H_Shaped_thickness | 0.39 | -0.21 | 0.09 | 0.91 |
| rh_S_parieto_occipital_thickness | 0.34 | -0.20 | -0.05 | 0.53 |
| rh_S_pericallosal_thickness | 0.10 | -0.05 | 0.82 | 1.07 |
| rh_S_postcentral_thickness | 0.39 | -0.23 | -0.08 | 0.50 |
| rh_S_precentral-inf-part_thickness | 0.33 | -0.19 | -0.37 | 1.19 |
| rh_S_precentral-sup-part_thickness | 0.26 | -0.15 | -0.55 | 1.79 |
| rh_S_suborbital_thickness | 0.11 | -0.06 | 0.31 | 0.21 |
| rh_S_subparietal_thickness | 0.27 | -0.15 | 0.18 | 0.76 |
| rh_S_temporal_inf_thickness | 0.18 | -0.10 | 0.20 | 1.13 |
| rh_S_temporal_sup_thickness | 0.41 | -0.25 | -0.18 | 0.70 |
| rh_S_temporal_transverse_thickness | 0.14 | -0.07 | 0.25 | 0.18 |
| rh_MeanThickness_thickness | 0.45 | -0.29 | -0.24 | 0.59 |
| Left-Lateral-Ventricle | 0.40 | -0.34 | 0.47 | 0.54 |
| Left-Inf-Lat-Vent | 0.30 | -0.19 | 0.26 | 0.79 |
| Left-Cerebellum-White-Matter | 0.31 | -0.20 | 0.32 | 2.15 |
| Left-Cerebellum-Cortex | 0.34 | -0.20 | -0.33 | 2.69 |
| Left-Thalamus-Proper | 0.58 | -0.43 | 0.13 | 1.57 |
| Left-Caudate | 0.32 | -0.19 | 0.06 | 1.24 |
| Left-Putamen | 0.46 | -0.30 | -0.22 | 2.02 |
| Left-Pallidum | 0.42 | -0.29 | -0.08 | 1.15 |
| 3rd-Ventricle | 0.47 | -0.37 | 0.68 | 1.15 |
| 4th-Ventricle | 0.09 | -0.05 | 0.35 | 0.48 |
| Brain-Stem | 0.37 | -0.24 | 0.03 | 1.04 |
| Left-Hippocampus | 0.26 | -0.15 | -0.18 | 1.48 |
| Left-Amygdala | 0.28 | -0.16 | -0.16 | 1.40 |
| CSF | 0.28 | -0.18 | 0.57 | 1.36 |
| Left-Accumbens-area | 0.60 | -0.43 | -0.20 | 1.23 |
| Left-VentralDC | 0.32 | -0.19 | 0.10 | 0.55 |
| Left-vessel | 0.17 | -0.09 | 0.12 | 0.72 |
| Left-choroid-plexus | 0.60 | -0.42 | -0.10 | 0.21 |
| Right-Lateral-Ventricle | 0.41 | -0.35 | 0.46 | 0.56 |
| Right-Inf-Lat-Vent | 0.25 | -0.14 | 0.14 | 0.71 |
| Right-Cerebellum-White-Matter | 0.29 | -0.19 | 0.57 | 2.83 |
| Right-Cerebellum-Cortex | 0.32 | -0.20 | -0.31 | 2.77 |
| Right-Thalamus-Proper | 0.58 | -0.42 | 0.08 | 0.94 |
| Right-Caudate | 0.33 | -0.19 | 0.08 | 1.09 |
| Right-Putamen | 0.43 | -0.28 | -0.16 | 2.11 |
| Right-Pallidum | 0.35 | -0.22 | -0.03 | 0.80 |
| Right-Hippocampus | 0.25 | -0.15 | -0.14 | 1.02 |
| Right-Amygdala | 0.29 | -0.17 | -0.10 | 1.12 |
| Right-Accumbens-area | 0.42 | -0.27 | -0.04 | 1.02 |
| Right-VentralDC | 0.33 | -0.19 | 0.12 | 0.64 |
| Right-vessel | 0.24 | -0.11 | 0.08 | 1.40 |
| Right-choroid-plexus | 0.68 | -0.45 | -0.04 | 0.25 |
| SubCortGrayVol | 0.61 | -0.42 | -0.07 | 0.97 |
| TotalGrayVol | 0.60 | -0.45 | -0.16 | 1.40 |
| SupraTentorialVol | 0.35 | -0.21 | -0.03 | 0.52 |
| SupraTentorialVolNotVent | 0.37 | -0.23 | -0.05 | 0.54 |
| avg_thickness | 0.48 | -0.31 | -0.27 | 0.71 |
| EstimatedTotalIntraCranialVol | 0.33 | -0.20 | -0.38 | 2.41 |

**Appendix 0 Table 6.**
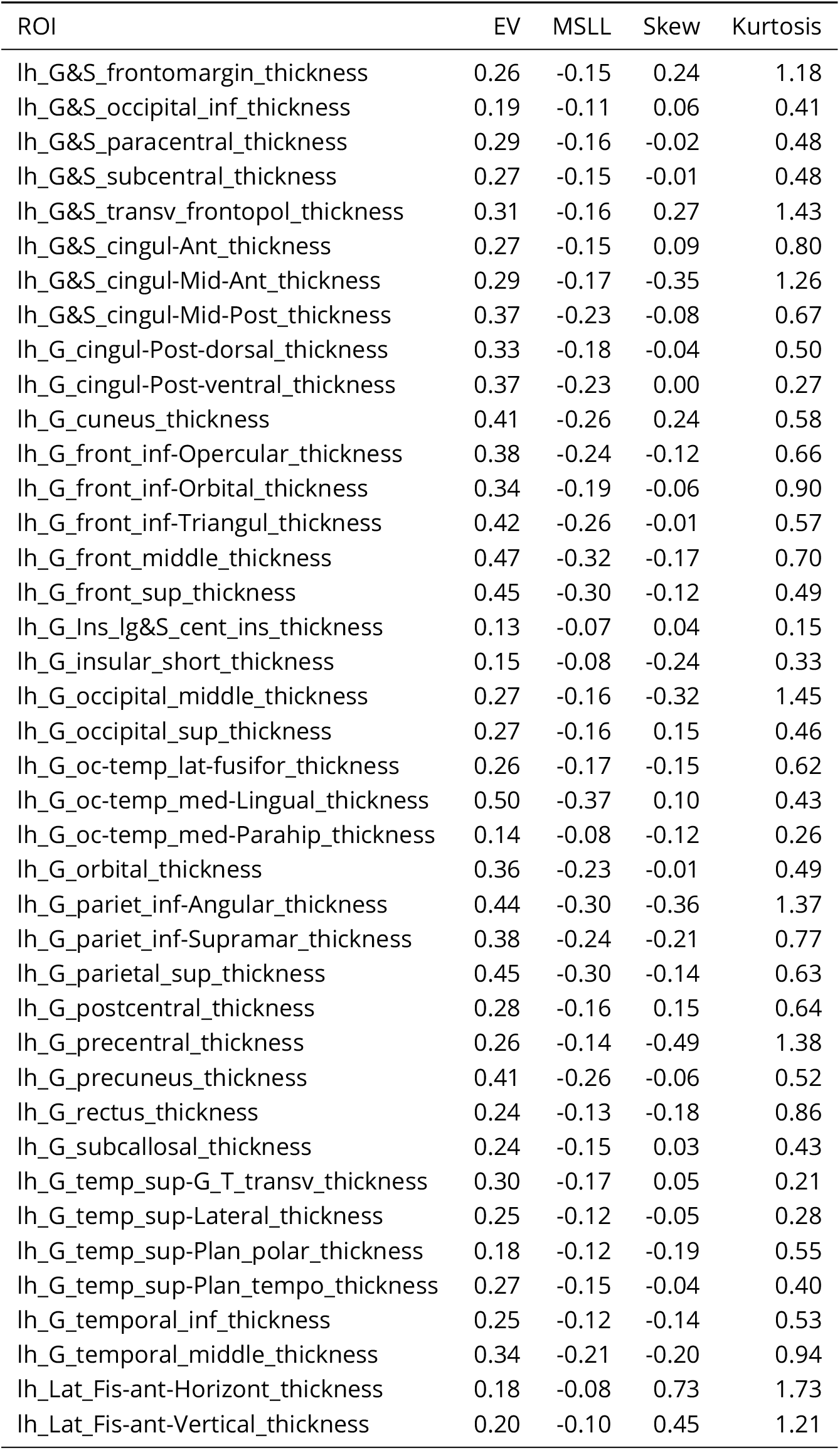
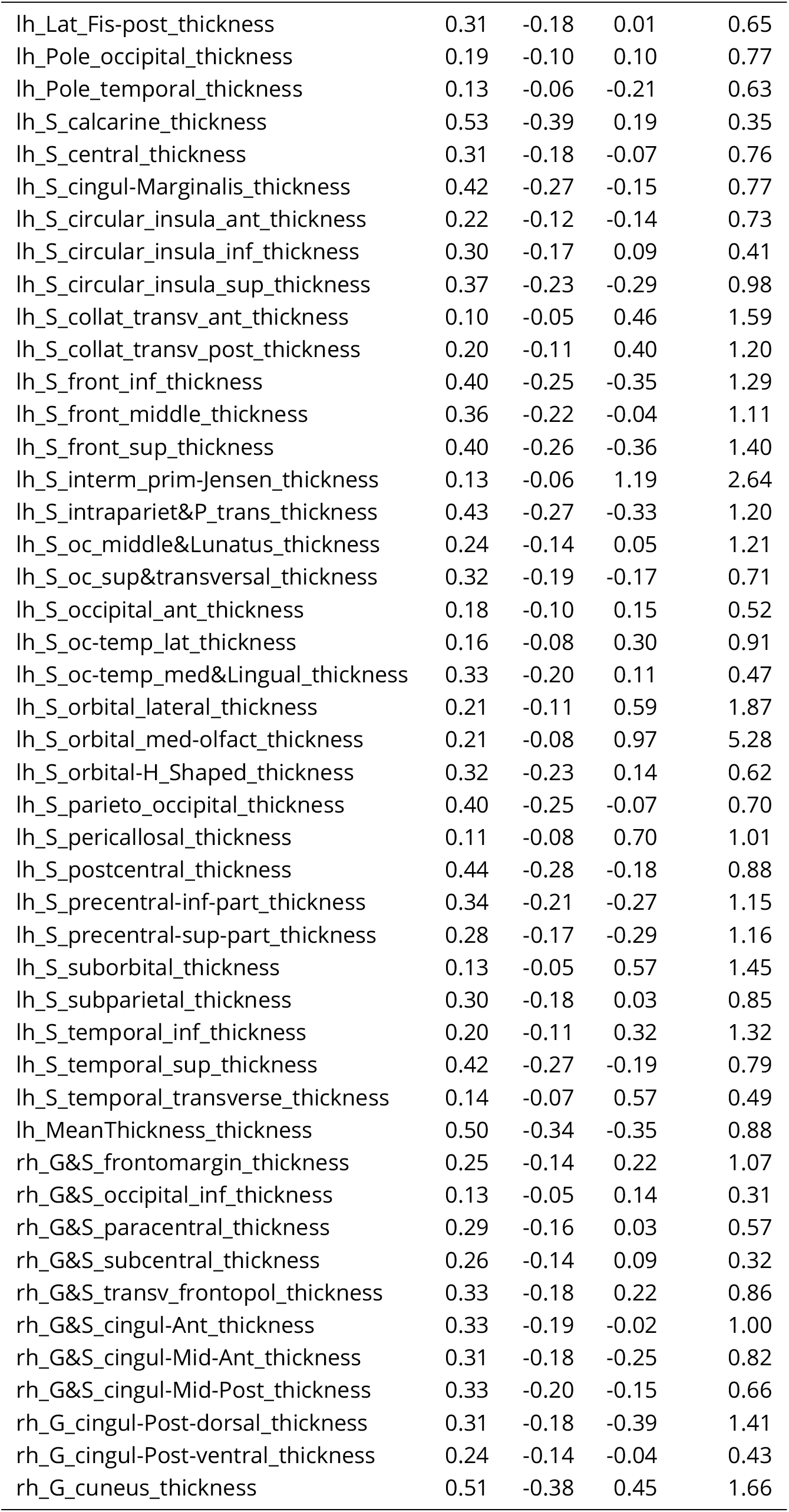
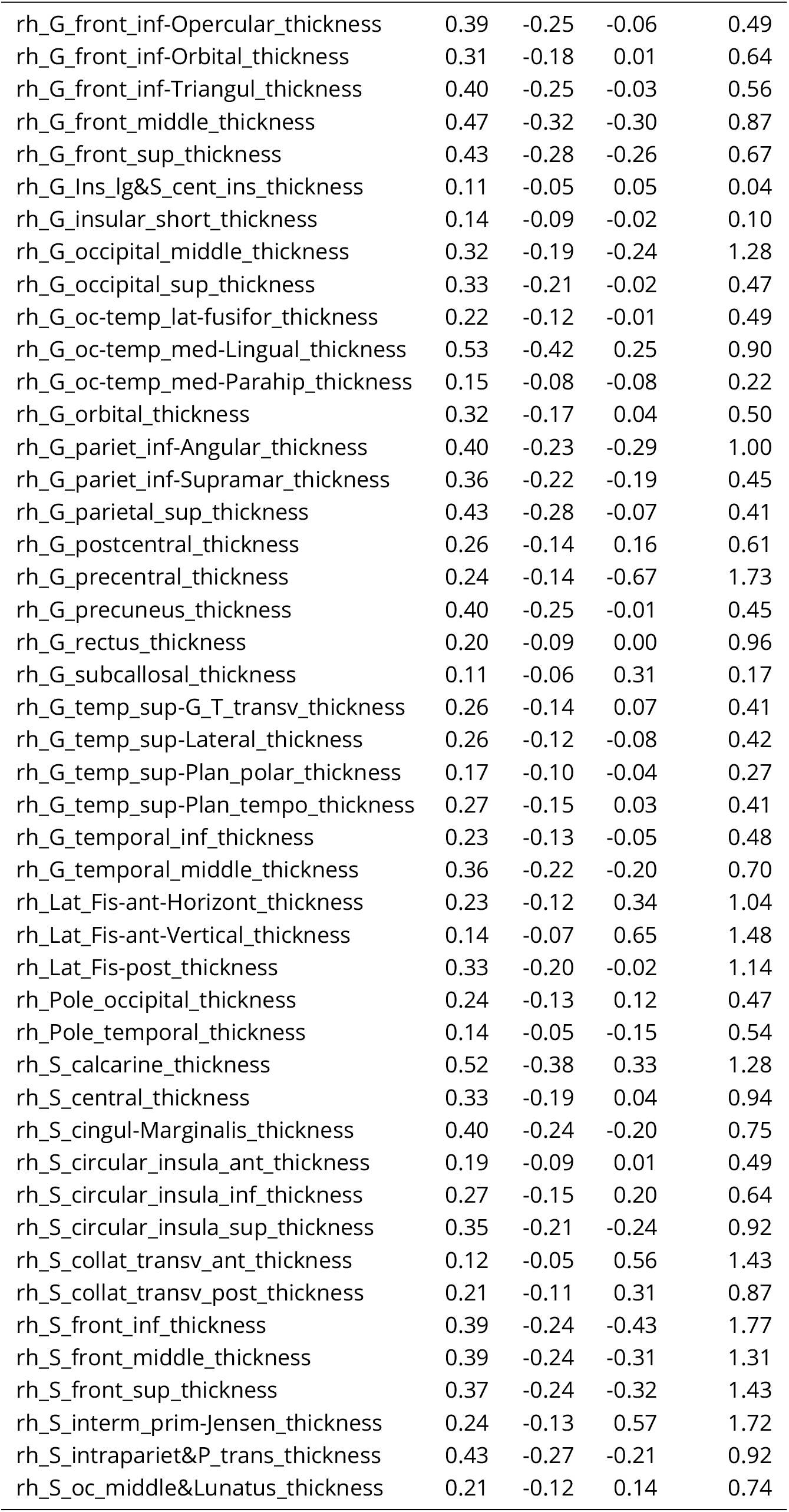
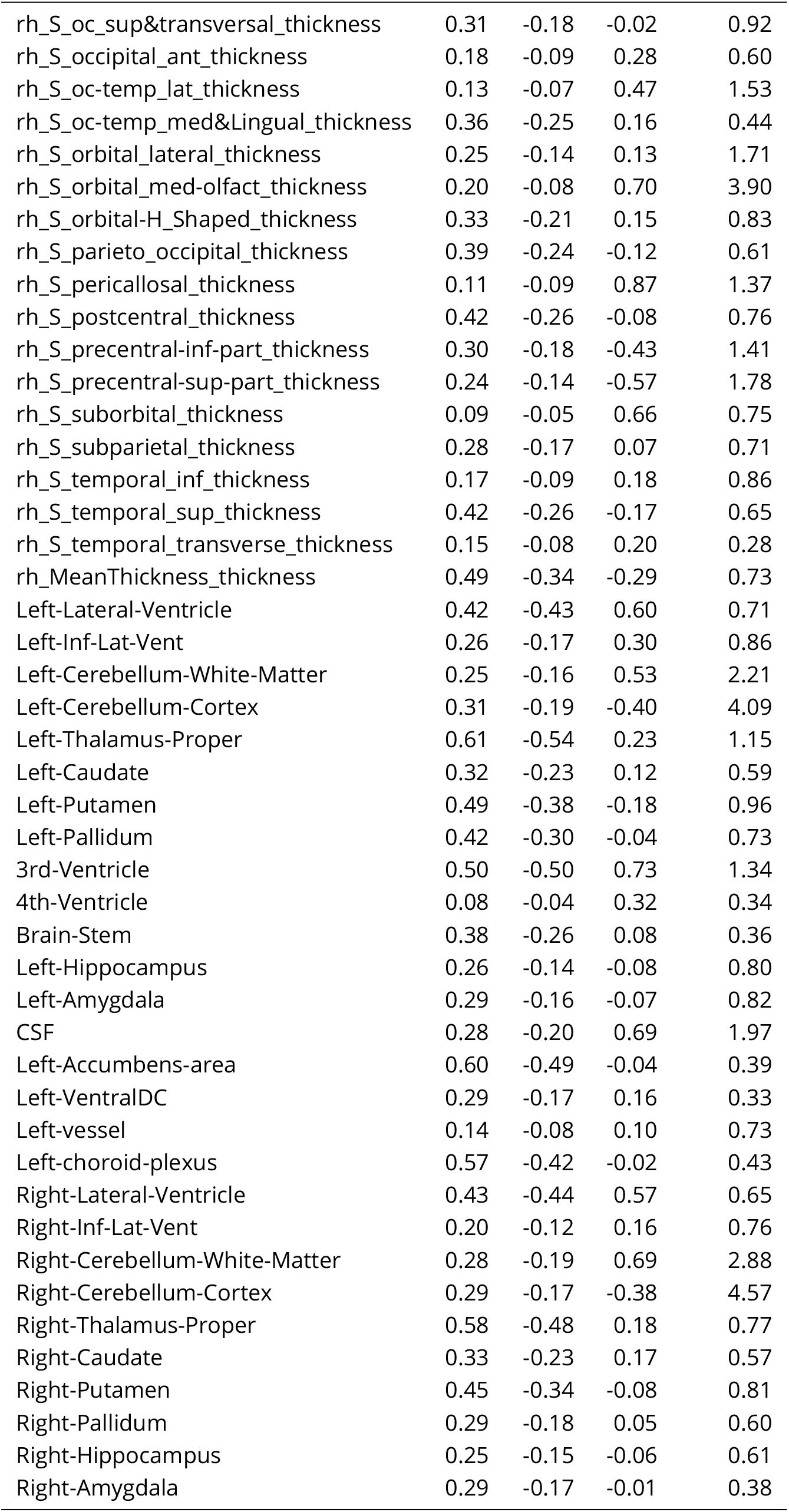
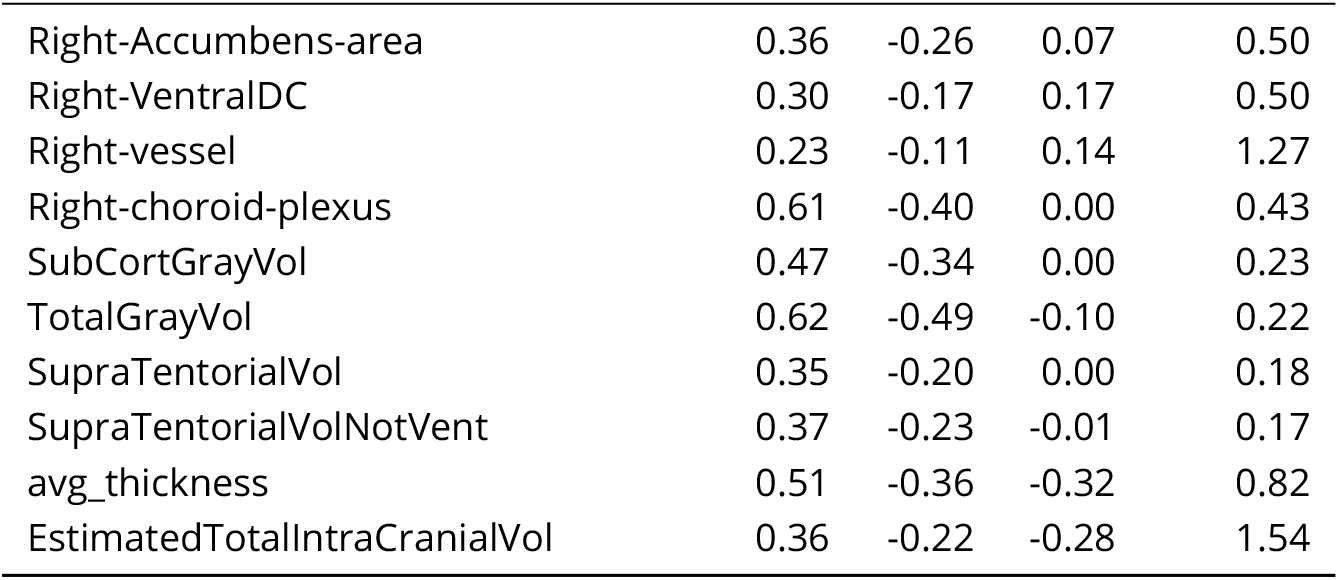
mQC Test Set Evaluation Metrics

| ROI | EV | MSLL | Skew | Kurtosis |
| --- | --- | --- | --- | --- |
| lh_G&S_frontomargin_thickness | 0.26 | -0.15 | 0.24 | 1.18 |
| lh_G&S_occipital_inf_thickness | 0.19 | -0.11 | 0.06 | 0.41 |
| lh_G&S_paracentral_thickness | 0.29 | -0.16 | -0.02 | 0.48 |
| lh_G&S_subcentral_thickness | 0.27 | -0.15 | -0.01 | 0.48 |
| lh_G&S_transv_frontopol_thickness | 0.31 | -0.16 | 0.27 | 1.43 |
| lh_G&S_cingul-Ant_thickness | 0.27 | -0.15 | 0.09 | 0.80 |
| lh_G&S_cingul-Mid-Ant_thickness | 0.29 | -0.17 | -0.35 | 1.26 |
| lh_G&S_cingul-Mid-Post_thickness | 0.37 | -0.23 | -0.08 | 0.67 |
| lh_G_cingul-Post-dorsal_thickness | 0.33 | -0.18 | -0.04 | 0.50 |
| lh_G_cingul-Post-ventral_thickness | 0.37 | -0.23 | 0.00 | 0.27 |
| lh_G_cuneus_thickness | 0.41 | -0.26 | 0.24 | 0.58 |
| lh_G_front_inf-Opercular_thickness | 0.38 | -0.24 | -0.12 | 0.66 |
| lh_G_front_inf-Orbital_thickness | 0.34 | -0.19 | -0.06 | 0.90 |
| lh_G_front_inf-Triangul_thickness | 0.42 | -0.26 | -0.01 | 0.57 |
| lh_G_front_middle_thickness | 0.47 | -0.32 | -0.17 | 0.70 |
| lh_G_front_sup_thickness | 0.45 | -0.30 | -0.12 | 0.49 |
| lh_G_Ins_Ig&S_cent_ins_thickness | 0.13 | -0.07 | 0.04 | 0.15 |
| lh_G_insular_short_thickness | 0.15 | -0.08 | -0.24 | 0.33 |
| lh_G_occipital_middle_thickness | 0.27 | -0.16 | -0.32 | 1.45 |
| lh_G_occipital_sup_thickness | 0.27 | -0.16 | 0.15 | 0.46 |
| lh_G_oc-temp_lat-fusifor_thickness | 0.26 | -0.17 | -0.15 | 0.62 |
| lh_G_oc-temp_med-Lingual_thickness | 0.50 | -0.37 | 0.10 | 0.43 |
| lh_G_oc-temp_med-Parahip_thickness | 0.14 | -0.08 | -0.12 | 0.26 |
| lh_G_orbital_thickness | 0.36 | -0.23 | -0.01 | 0.49 |
| lh_G_pariet_inf-Angular_thickness | 0.44 | -0.30 | -0.36 | 1.37 |
| lh_G_pariet_inf-Supramar_thickness | 0.38 | -0.24 | -0.21 | 0.77 |
| lh_G_parietal_sup_thickness | 0.45 | -0.30 | -0.14 | 0.63 |
| lh_G_postcentral_thickness | 0.28 | -0.16 | 0.15 | 0.64 |
| lh_G_precentral_thickness | 0.26 | -0.14 | -0.49 | 1.38 |
| lh_G_precuneus_thickness | 0.41 | -0.26 | -0.06 | 0.52 |
| lh_G_rectus_thickness | 0.24 | -0.13 | -0.18 | 0.86 |
| lh_G_subcallosal_thickness | 0.24 | -0.15 | 0.03 | 0.43 |
| lh_G_temp_sup-G_T_transv_thickness | 0.30 | -0.17 | 0.05 | 0.21 |
| lh_G_temp_sup-Lateral_thickness | 0.25 | -0.12 | -0.05 | 0.28 |
| lh_G_temp_sup-Plan_polar_thickness | 0.18 | -0.12 | -0.19 | 0.55 |
| lh_G_temp_sup-Plan_tempo_thickness | 0.27 | -0.15 | -0.04 | 0.40 |
| lh_G_temporal_inf_thickness | 0.25 | -0.12 | -0.14 | 0.53 |
| lh_G_temporal_middle_thickness | 0.34 | -0.21 | -0.20 | 0.94 |
| lh_Lat_Fis-ant-Horizont_thickness | 0.18 | -0.08 | 0.73 | 1.73 |
| lh_Lat_Fis-ant-Vertical_thickness | 0.20 | -0.10 | 0.45 | 1.21 |
| lh_Lat_Fis-post_thickness | 0.31 | -0.18 | 0.01 | 0.65 |
| lh_Pole_occipital_thickness | 0.19 | -0.10 | 0.10 | 0.77 |
| lh_Pole_temporal_thickness | 0.13 | -0.06 | -0.21 | 0.63 |
| lh_S_calcarine_thickness | 0.53 | -0.39 | 0.19 | 0.35 |
| lh_S_central_thickness | 0.31 | -0.18 | -0.07 | 0.76 |
| lh_S_cingul-Marginalis_thickness | 0.42 | -0.27 | -0.15 | 0.77 |
| lh_S_circular_insula_ant_thickness | 0.22 | -0.12 | -0.14 | 0.73 |
| lh_S_circular_insula_inf_thickness | 0.30 | -0.17 | 0.09 | 0.41 |
| lh_S_circular_insula_sup_thickness | 0.37 | -0.23 | -0.29 | 0.98 |
| lh_S_collat_transv_ant_thickness | 0.10 | -0.05 | 0.46 | 1.59 |
| lh_S_collat_transv_post_thickness | 0.20 | -0.11 | 0.40 | 1.20 |
| lh_S_front_inf_thickness | 0.40 | -0.25 | -0.35 | 1.29 |
| lh_S_front_middle_thickness | 0.36 | -0.22 | -0.04 | 1.11 |
| lh_S_front_sup_thickness | 0.40 | -0.26 | -0.36 | 1.40 |
| lh_S_interm_prim-Jensen_thickness | 0.13 | -0.06 | 1.19 | 2.64 |
| lh_S_intrapariet&P_trans_thickness | 0.43 | -0.27 | -0.33 | 1.20 |
| lh_S_oc_middle&Lunatus_thickness | 0.24 | -0.14 | 0.05 | 1.21 |
| lh_S_oc_sup&transversal_thickness | 0.32 | -0.19 | -0.17 | 0.71 |
| lh_S_occipital_ant_thickness | 0.18 | -0.10 | 0.15 | 0.52 |
| lh_S_oc-temp_lat_thickness | 0.16 | -0.08 | 0.30 | 0.91 |
| lh_S_oc-temp_med&Lingual_thickness | 0.33 | -0.20 | 0.11 | 0.47 |
| lh_S_orbital_lateral_thickness | 0.21 | -0.11 | 0.59 | 1.87 |
| lh_S_orbital_med-olfact_thickness | 0.21 | -0.08 | 0.97 | 5.28 |
| lh_S_orbital-H-Shaped_thickness | 0.32 | -0.23 | 0.14 | 0.62 |
| lh_S_parieto_occipital_thickness | 0.40 | -0.25 | -0.07 | 0.70 |
| lh_S_pericallosal_thickness | 0.11 | -0.08 | 0.70 | 1.01 |
| lh_S_postcentral_thickness | 0.44 | -0.28 | -0.18 | 0.88 |
| lh_S_precentral-inf-part_thickness | 0.34 | -0.21 | -0.27 | 1.15 |
| lh_S_precentral-sup-part_thickness | 0.28 | -0.17 | -0.29 | 1.16 |
| lh_S_suborbital_thickness | 0.13 | -0.05 | 0.57 | 1.45 |
| lh_S_subparietal_thickness | 0.30 | -0.18 | 0.03 | 0.85 |
| lh_S_temporal_inf_thickness | 0.20 | -0.11 | 0.32 | 1.32 |
| lh_S_temporal_sup_thickness | 0.42 | -0.27 | -0.19 | 0.79 |
| lh_S_temporal_transverse_thickness | 0.14 | -0.07 | 0.57 | 0.49 |
| lh_MeanThickness_thickness | 0.50 | -0.34 | -0.35 | 0.88 |
| rh_G&S_frontomargin_thickness | 0.25 | -0.14 | 0.22 | 1.07 |
| rh_G&S_occipital_inf_thickness | 0.13 | -0.05 | 0.14 | 0.31 |
| rh_G&S_paracentral_thickness | 0.29 | -0.16 | 0.03 | 0.57 |
| rh_G&S_subcentral_thickness | 0.26 | -0.14 | 0.09 | 0.32 |
| rh_G&S_transv_frontopol_thickness | 0.33 | -0.18 | 0.22 | 0.86 |
| rh_G&S_cingul-Ant_thickness | 0.33 | -0.19 | -0.02 | 1.00 |
| rh_G&S_cingul-Mid-Ant_thickness | 0.31 | -0.18 | -0.25 | 0.82 |
| rh_G&S_cingul-Mid-Post_thickness | 0.33 | -0.20 | -0.15 | 0.66 |
| rh_G_cingul-Post-dorsal_thickness | 0.31 | -0.18 | -0.39 | 1.41 |
| rh_G_cingul-Post-ventral_thickness | 0.24 | -0.14 | -0.04 | 0.43 |
| rh_G_cuneus_thickness | 0.51 | -0.38 | 0.45 | 1.66 |
| rh_G_front_inf-Opercular_thickness | 0.39 | -0.25 | -0.06 | 0.49 |
| rh_G_front_inf-Orbital_thickness | 0.31 | -0.18 | 0.01 | 0.64 |
| rh_G_front_inf-Triangul_thickness | 0.40 | -0.25 | -0.03 | 0.56 |
| rh_G_front_middle_thickness | 0.47 | -0.32 | -0.30 | 0.87 |
| rh_G_front_sup_thickness | 0.43 | -0.28 | -0.26 | 0.67 |
| rh_G_Ins_Ig&S_cent_ins_thickness | 0.11 | -0.05 | 0.05 | 0.04 |
| rh_G_insular_short_thickness | 0.14 | -0.09 | -0.02 | 0.10 |
| rh_G_occipital_middle_thickness | 0.32 | -0.19 | -0.24 | 1.28 |
| rh_G_occipital_sup_thickness | 0.33 | -0.21 | -0.02 | 0.47 |
| rh_G_oc-temp_lat-fusifor_thickness | 0.22 | -0.12 | -0.01 | 0.49 |
| rh_G_oc-temp_med-Lingual_thickness | 0.53 | -0.42 | 0.25 | 0.90 |
| rh_G_oc-temp_med-Parahip_thickness | 0.15 | -0.08 | -0.08 | 0.22 |
| rh_G_orbital_thickness | 0.32 | -0.17 | 0.04 | 0.50 |
| rh_G_pariet_inf-Angular_thickness | 0.40 | -0.23 | -0.29 | 1.00 |
| rh_G_pariet_inf-Supramar_thickness | 0.36 | -0.22 | -0.19 | 0.45 |
| rh_G_parietal_sup_thickness | 0.43 | -0.28 | -0.07 | 0.41 |
| rh_G_postcentral_thickness | 0.26 | -0.14 | 0.16 | 0.61 |
| rh_G_precentral_thickness | 0.24 | -0.14 | -0.67 | 1.73 |
| rh_G_precuneus_thickness | 0.40 | -0.25 | -0.01 | 0.45 |
| rh_G_rectus_thickness | 0.20 | -0.09 | 0.00 | 0.96 |
| rh_G_subcallosal_thickness | 0.11 | -0.06 | 0.31 | 0.17 |
| rh_G_temp_sup-G_T_transv_thickness | 0.26 | -0.14 | 0.07 | 0.41 |
| rh_G_temp_sup-Lateral_thickness | 0.26 | -0.12 | -0.08 | 0.42 |
| rh_G_temp_sup-Plan_polar_thickness | 0.17 | -0.10 | -0.04 | 0.27 |
| rh_G_temp_sup-Plan_tempo_thickness | 0.27 | -0.15 | 0.03 | 0.41 |
| rh_G_temporal_inf_thickness | 0.23 | -0.13 | -0.05 | 0.48 |
| rh_G_temporal_middle_thickness | 0.36 | -0.22 | -0.20 | 0.70 |
| rh_Lat_Fis-ant-Horizont_thickness | 0.23 | -0.12 | 0.34 | 1.04 |
| rh_Lat_Fis-ant-Vertical_thickness | 0.14 | -0.07 | 0.65 | 1.48 |
| rh_Lat_Fis-post_thickness | 0.33 | -0.20 | -0.02 | 1.14 |
| rh_Pole_occipital_thickness | 0.24 | -0.13 | 0.12 | 0.47 |
| rh_Pole_temporal_thickness | 0.14 | -0.05 | -0.15 | 0.54 |
| rh_S_calcarine_thickness | 0.52 | -0.38 | 0.33 | 1.28 |
| rh_S_central_thickness | 0.33 | -0.19 | 0.04 | 0.94 |
| rh_S_cingul-Marginalis_thickness | 0.40 | -0.24 | -0.20 | 0.75 |
| rh_S_circular_insula_ant_thickness | 0.19 | -0.09 | 0.01 | 0.49 |
| rh_S_circular_insula_inf_thickness | 0.27 | -0.15 | 0.20 | 0.64 |
| rh_S_circular_insula_sup_thickness | 0.35 | -0.21 | -0.24 | 0.92 |
| rh_S_collat_transv_ant_thickness | 0.12 | -0.05 | 0.56 | 1.43 |
| rh_S_collat_transv_post_thickness | 0.21 | -0.11 | 0.31 | 0.87 |
| rh_S_front_inf_thickness | 0.39 | -0.24 | -0.43 | 1.77 |
| rh_S_front_middle_thickness | 0.39 | -0.24 | -0.31 | 1.31 |
| rh_S_front_sup_thickness | 0.37 | -0.24 | -0.32 | 1.43 |
| rh_S_interm_prim-Jensen_thickness | 0.24 | -0.13 | 0.57 | 1.72 |
| rh_S_intrapariet&P_trans_thickness | 0.43 | -0.27 | -0.21 | 0.92 |
| rh_S_oc_middle&Lunatus_thickness | 0.21 | -0.12 | 0.14 | 0.74 |
| rh_S_oc_sup&transversal_thickness | 0.31 | -0.18 | -0.02 | 0.92 |
| rh_S_occipital_ant_thickness | 0.18 | -0.09 | 0.28 | 0.60 |
| rh_S_oc-temp_lat_thickness | 0.13 | -0.07 | 0.47 | 1.53 |
| rh_S_oc-temp_med&Lingual_thickness | 0.36 | -0.25 | 0.16 | 0.44 |
| rh_S_orbital_lateral_thickness | 0.25 | -0.14 | 0.13 | 1.71 |
| rh_S_orbital_med-olfact_thickness | 0.20 | -0.08 | 0.70 | 3.90 |
| rh_S_orbital-H_Shaped_thickness | 0.33 | -0.21 | 0.15 | 0.83 |
| rh_S_parieto_occipital_thickness | 0.39 | -0.24 | -0.12 | 0.61 |
| rh_S_pericallosal_thickness | 0.11 | -0.09 | 0.87 | 1.37 |
| rh_S_postcentral_thickness | 0.42 | -0.26 | -0.08 | 0.76 |
| rh_S_precentral-inf-part_thickness | 0.30 | -0.18 | -0.43 | 1.41 |
| rh_S_precentral-sup-part_thickness | 0.24 | -0.14 | -0.57 | 1.78 |
| rh_S_suborbital_thickness | 0.09 | -0.05 | 0.66 | 0.75 |
| rh_S_subparietal_thickness | 0.28 | -0.17 | 0.07 | 0.71 |
| rh_S_temporal_inf_thickness | 0.17 | -0.09 | 0.18 | 0.86 |
| rh_S_temporal_sup_thickness | 0.42 | -0.26 | -0.17 | 0.65 |
| rh_S_temporal_transverse_thickness | 0.15 | -0.08 | 0.20 | 0.28 |
| rh_MeanThickness_thickness | 0.49 | -0.34 | -0.29 | 0.73 |
| Left-Lateral-Ventricle | 0.42 | -0.43 | 0.60 | 0.71 |
| Left-Inf-Lat-Vent | 0.26 | -0.17 | 0.30 | 0.86 |
| Left-Cerebellum-White-Matter | 0.25 | -0.16 | 0.53 | 2.21 |
| Left-Cerebellum-Cortex | 0.31 | -0.19 | -0.40 | 4.09 |
| Left-Thalamus-Proper | 0.61 | -0.54 | 0.23 | 1.15 |
| Left-Caudate | 0.32 | -0.23 | 0.12 | 0.59 |
| Left-Putamen | 0.49 | -0.38 | -0.18 | 0.96 |
| Left-Pallidum | 0.42 | -0.30 | -0.04 | 0.73 |
| 3rd-Ventricle | 0.50 | -0.50 | 0.73 | 1.34 |
| 4th-Ventricle | 0.08 | -0.04 | 0.32 | 0.34 |
| Brain-Stem | 0.38 | -0.26 | 0.08 | 0.36 |
| Left-Hippocampus | 0.26 | -0.14 | -0.08 | 0.80 |
| Left-Amygdala | 0.29 | -0.16 | -0.07 | 0.82 |
| CSF | 0.28 | -0.20 | 0.69 | 1.97 |
| Left-Accumbens-area | 0.60 | -0.49 | -0.04 | 0.39 |
| Left-VentralDC | 0.29 | -0.17 | 0.16 | 0.33 |
| Left-vessel | 0.14 | -0.08 | 0.10 | 0.73 |
| Left-choroid-plexus | 0.57 | -0.42 | -0.02 | 0.43 |
| Right-Lateral-Ventricle | 0.43 | -0.44 | 0.57 | 0.65 |
| Right-Inf-Lat-Vent | 0.20 | -0.12 | 0.16 | 0.76 |
| Right-Cerebellum-White-Matter | 0.28 | -0.19 | 0.69 | 2.88 |
| Right-Cerebellum-Cortex | 0.29 | -0.17 | -0.38 | 4.57 |
| Right-Thalamus-Proper | 0.58 | -0.48 | 0.18 | 0.77 |
| Right-Caudate | 0.33 | -0.23 | 0.17 | 0.57 |
| Right-Putamen | 0.45 | -0.34 | -0.08 | 0.81 |
| Right-Pallidum | 0.29 | -0.18 | 0.05 | 0.60 |
| Right-Hippocampus | 0.25 | -0.15 | -0.06 | 0.61 |
| Right-Amygdala | 0.29 | -0.17 | -0.01 | 0.38 |
| Right-Accumbens-area | 0.36 | -0.26 | 0.07 | 0.50 |
| Right-VentralDC | 0.30 | -0.17 | 0.17 | 0.50 |
| Right-vessel | 0.23 | -0.11 | 0.14 | 1.27 |
| Right-choroid-plexus | 0.61 | -0.40 | 0.00 | 0.43 |
| SubCortGrayVol | 0.47 | -0.34 | 0.00 | 0.23 |
| TotalGrayVol | 0.62 | -0.49 | -0.10 | 0.22 |
| SupraTentorialVol | 0.35 | -0.20 | 0.00 | 0.18 |
| SupraTentorialVolNotVent | 0.37 | -0.23 | -0.01 | 0.17 |
| avg_thickness | 0.51 | -0.36 | -0.32 | 0.82 |
| EstimatedTotalIntraCranialVol | 0.36 | -0.22 | -0.28 | 1.54 |

**Appendix 0 Table 7.**
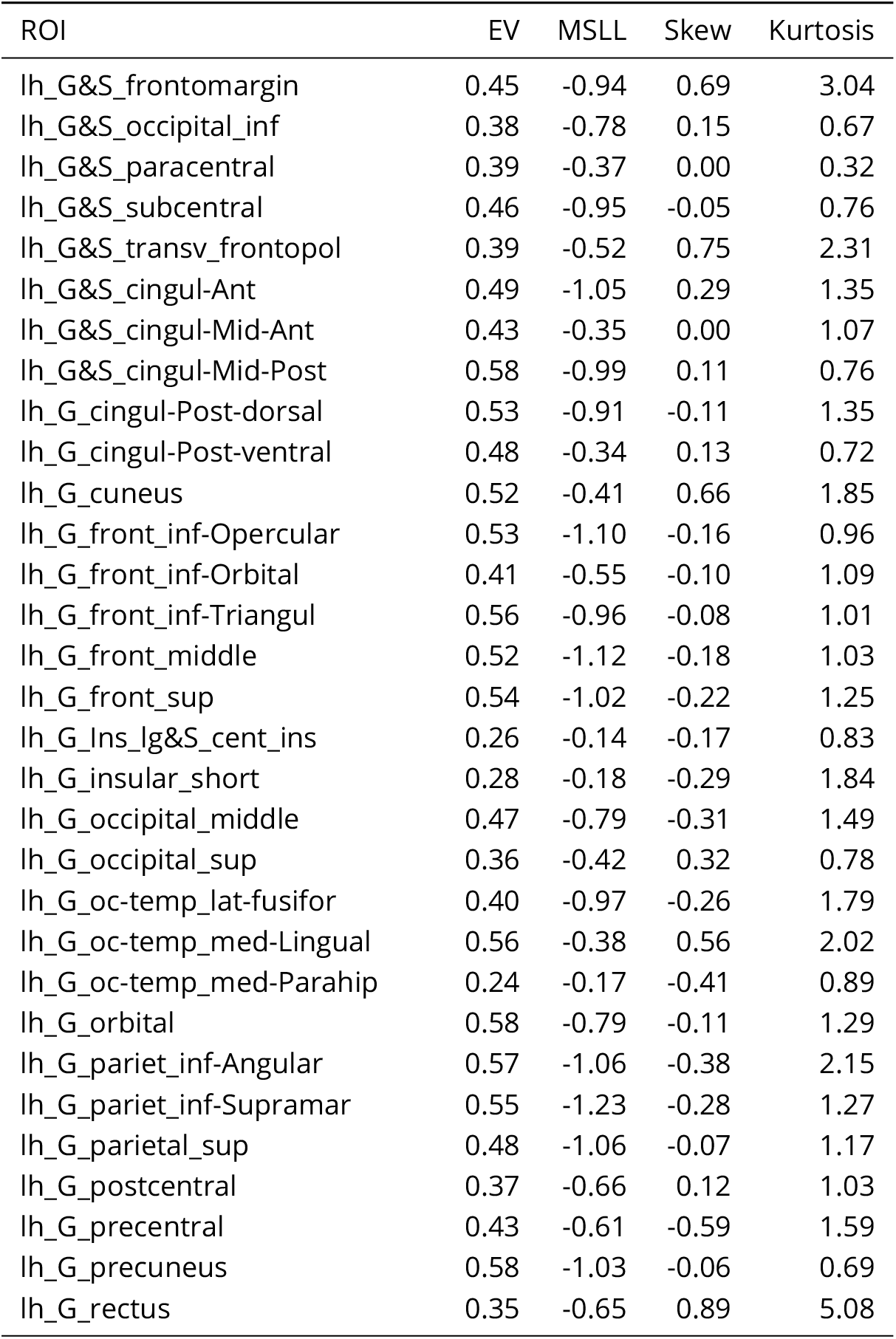
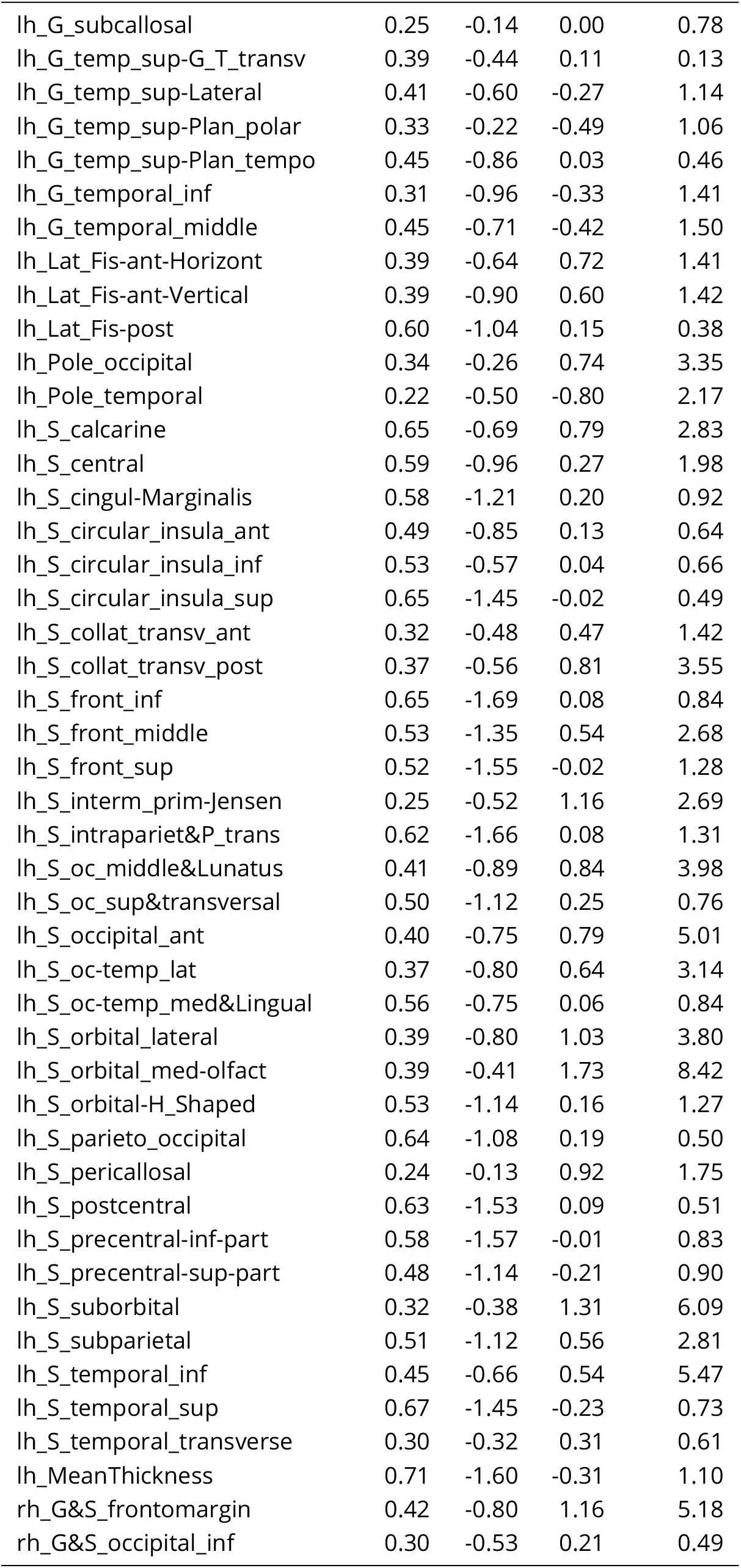
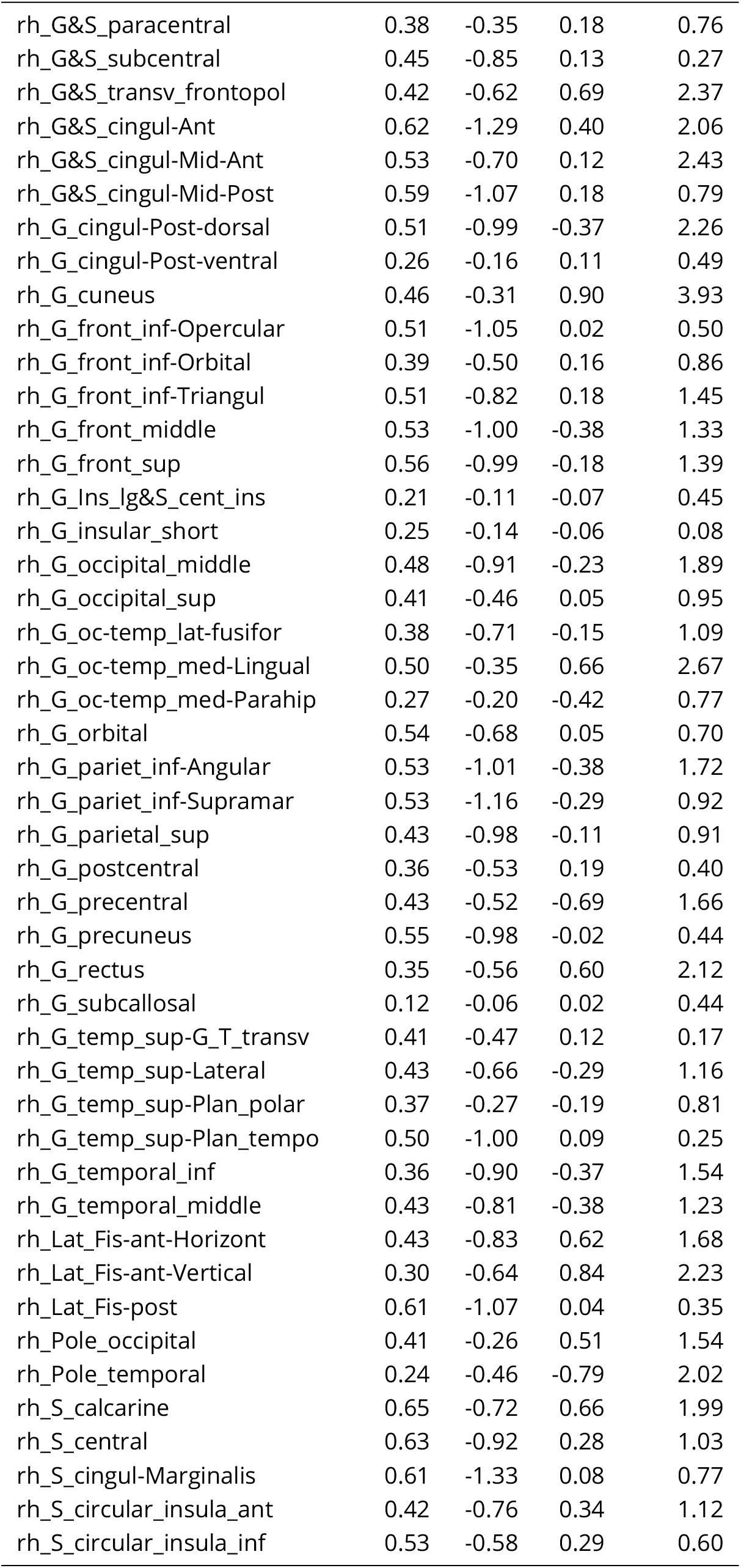
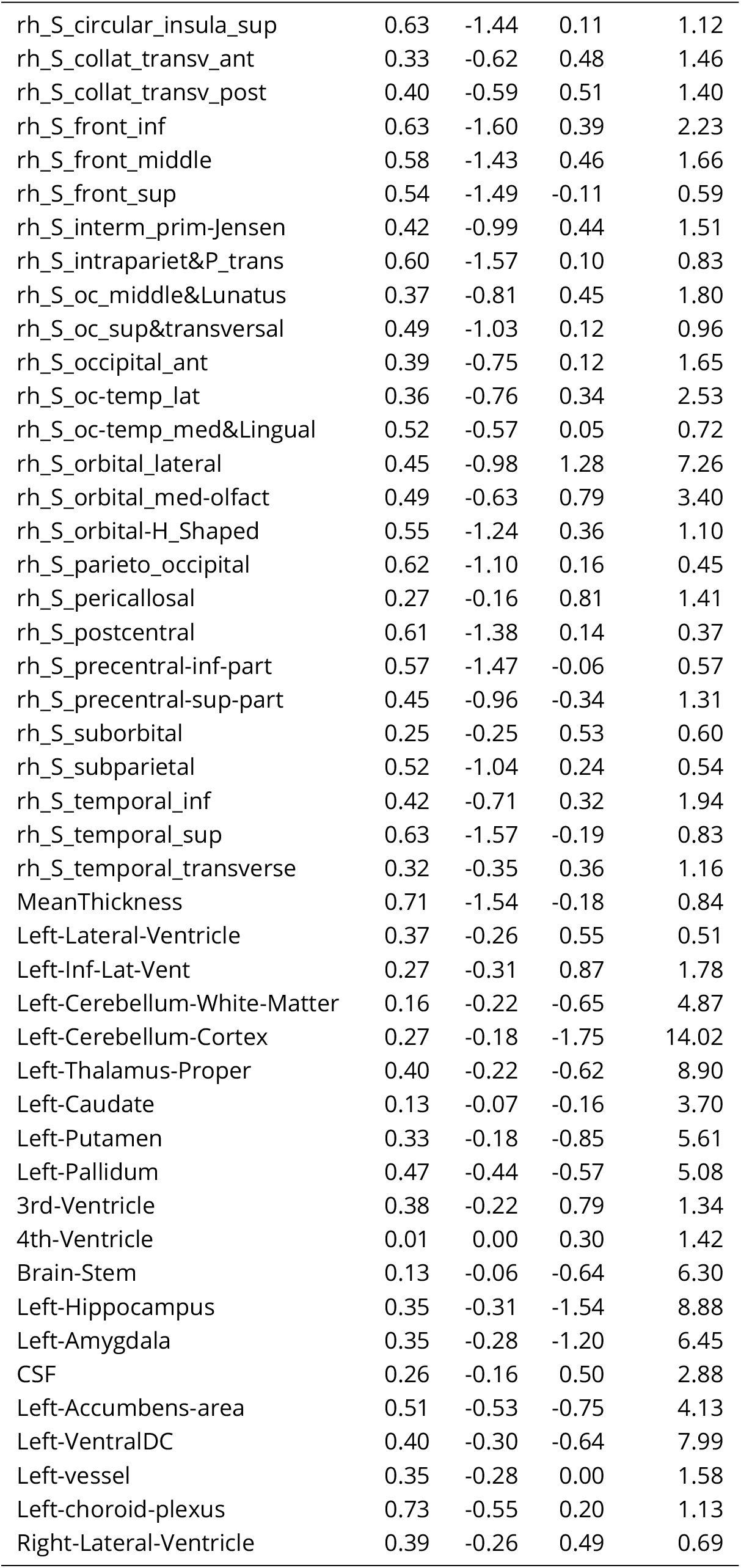
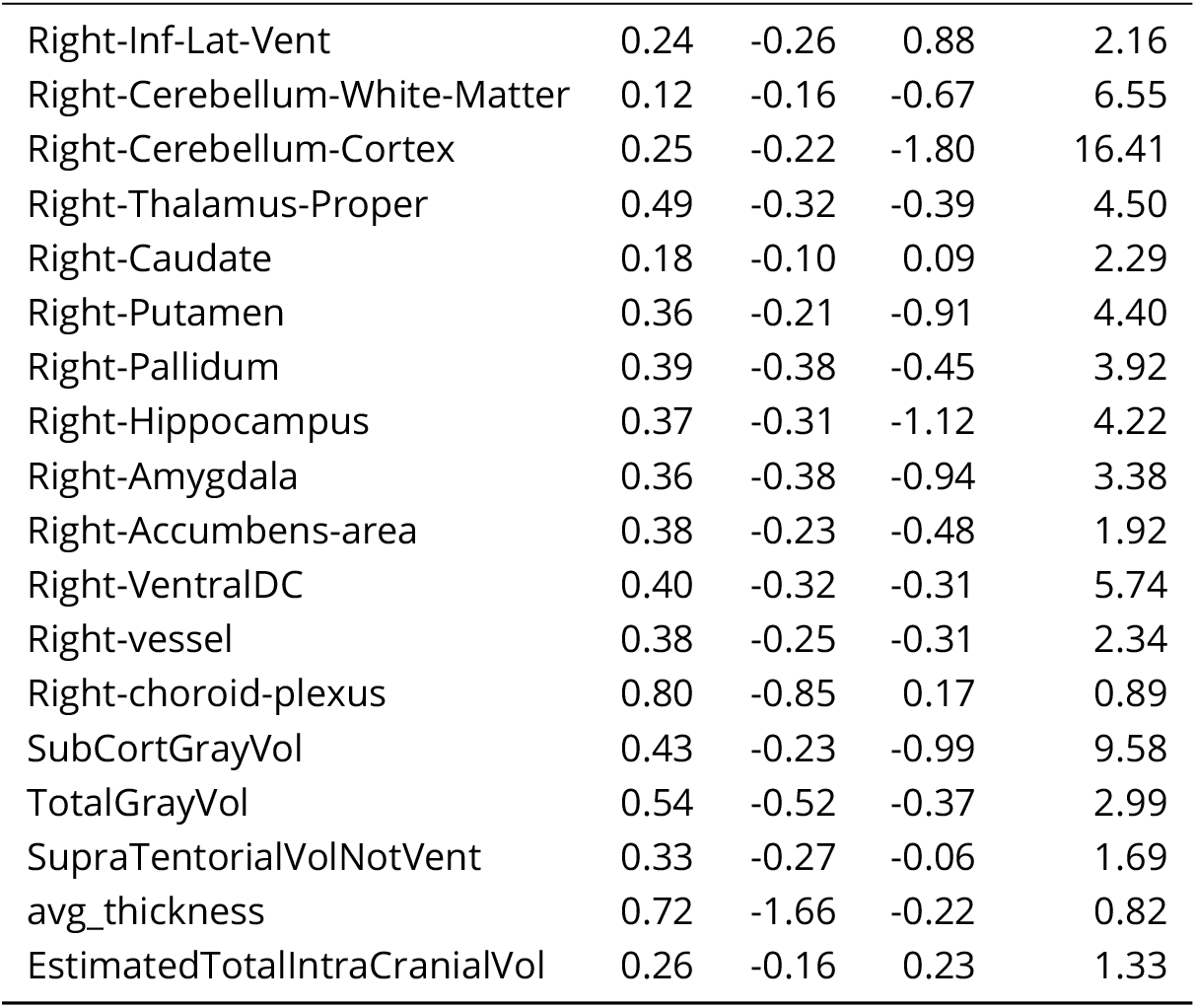
Patient Test Set Evaluation Metrics

| ROI | EV | MSLL | Skew | Kurtosis |
| --- | --- | --- | --- | --- |
| lh_G&S_frontomargin | 0.45 | -0.94 | 0.69 | 3.04 |
| lh_G&S_occipital_inf | 0.38 | -0.78 | 0.15 | 0.67 |
| lh_G&S_paracentral | 0.39 | -0.37 | 0.00 | 0.32 |
| lh_G&S_subcentral | 0.46 | -0.95 | -0.05 | 0.76 |
| lh_G&S_transv_frontopol | 0.39 | -0.52 | 0.75 | 2.31 |
| lh_G&S_cingul-Ant | 0.49 | -1.05 | 0.29 | 1.35 |
| lh_G&S_cingul-Mid-Ant | 0.43 | -0.35 | 0.00 | 1.07 |
| lh_G&S_cingul-Mid-Post | 0.58 | -0.99 | 0.11 | 0.76 |
| lh_G_cingul-Post-dorsal | 0.53 | -0.91 | -0.11 | 1.35 |
| lh_G_cingul-Post-ventral | 0.48 | -0.34 | 0.13 | 0.72 |
| lh_G_cuneus | 0.52 | -0.41 | 0.66 | 1.85 |
| lh_G_front_inf-Opercular | 0.53 | -1.10 | -0.16 | 0.96 |
| lh_G_front_inf-Orbital | 0.41 | -0.55 | -0.10 | 1.09 |
| lh_G_front_inf-Triangul | 0.56 | -0.96 | -0.08 | 1.01 |
| lh_G_front_middle | 0.52 | -1.12 | -0.18 | 1.03 |
| lh_G_front_sup | 0.54 | -1.02 | -0.22 | 1.25 |
| lh_G_Ins_lg&S_cent_ins | 0.26 | -0.14 | -0.17 | 0.83 |
| lh_G_insular_short | 0.28 | -0.18 | -0.29 | 1.84 |
| lh_G_occipital_middle | 0.47 | -0.79 | -0.31 | 1.49 |
| lh_G_occipital_sup | 0.36 | -0.42 | 0.32 | 0.78 |
| lh_G_oc-temp_lat-fusifor | 0.40 | -0.97 | -0.26 | 1.79 |
| lh_G_oc-temp_med-Lingual | 0.56 | -0.38 | 0.56 | 2.02 |
| lh_G_oc-temp_med-Parahip | 0.24 | -0.17 | -0.41 | 0.89 |
| lh_G_orbital | 0.58 | -0.79 | -0.11 | 1.29 |
| lh_G_pariet_inf-Angular | 0.57 | -1.06 | -0.38 | 2.15 |
| lh_G_pariet_inf-Supramar | 0.55 | -1.23 | -0.28 | 1.27 |
| lh_G_parietal_sup | 0.48 | -1.06 | -0.07 | 1.17 |
| lh_G_postcentral | 0.37 | -0.66 | 0.12 | 1.03 |
| lh_G_precentral | 0.43 | -0.61 | -0.59 | 1.59 |
| lh_G_precuneus | 0.58 | -1.03 | -0.06 | 0.69 |
| lh_G_rectus | 0.35 | -0.65 | 0.89 | 5.08 |
| lh_G_subcallosal | 0.25 | -0.14 | 0.00 | 0.78 |
| lh_G_temp_sup-G_T_transv | 0.39 | -0.44 | 0.11 | 0.13 |
| lh_G_temp_sup-Lateral | 0.41 | -0.60 | -0.27 | 1.14 |
| lh_G_temp_sup-Plan_polar | 0.33 | -0.22 | -0.49 | 1.06 |
| lh_G_temp_sup-Plan_tempo | 0.45 | -0.86 | 0.03 | 0.46 |
| lh_G_temporal_inf | 0.31 | -0.96 | -0.33 | 1.41 |
| lh_G_temporal_middle | 0.45 | -0.71 | -0.42 | 1.50 |
| lh_Lat_Fis-ant-Horizont | 0.39 | -0.64 | 0.72 | 1.41 |
| lh_Lat_Fis-ant-Vertical | 0.39 | -0.90 | 0.60 | 1.42 |
| lh_Lat_Fis-post | 0.60 | -1.04 | 0.15 | 0.38 |
| lh_Pole_occipital | 0.34 | -0.26 | 0.74 | 3.35 |
| lh_Pole_temporal | 0.22 | -0.50 | -0.80 | 2.17 |
| lh_S_calcarine | 0.65 | -0.69 | 0.79 | 2.83 |
| lh_S_central | 0.59 | -0.96 | 0.27 | 1.98 |
| lh_S_cingul-Marginalis | 0.58 | -1.21 | 0.20 | 0.92 |
| lh_S_circular_insula_ant | 0.49 | -0.85 | 0.13 | 0.64 |
| lh_S_circular_insula_inf | 0.53 | -0.57 | 0.04 | 0.66 |
| lh_S_circular_insula_sup | 0.65 | -1.45 | -0.02 | 0.49 |
| lh_S_collat_transv_ant | 0.32 | -0.48 | 0.47 | 1.42 |
| lh_S_collat_transv_post | 0.37 | -0.56 | 0.81 | 3.55 |
| lh_S_front_inf | 0.65 | -1.69 | 0.08 | 0.84 |
| lh_S_front_middle | 0.53 | -1.35 | 0.54 | 2.68 |
| lh_S_front_sup | 0.52 | -1.55 | -0.02 | 1.28 |
| lh_S_interm_prim-Jensen | 0.25 | -0.52 | 1.16 | 2.69 |
| lh_S_intrapariet&P_trans | 0.62 | -1.66 | 0.08 | 1.31 |
| lh_S_oc_middle&Lunatus | 0.41 | -0.89 | 0.84 | 3.98 |
| lh_S_oc_sup&transversal | 0.50 | -1.12 | 0.25 | 0.76 |
| lh_S_occipital_ant | 0.40 | -0.75 | 0.79 | 5.01 |
| lh_S_oc-temp_lat | 0.37 | -0.80 | 0.64 | 3.14 |
| lh_S_oc-temp_med&Lingual | 0.56 | -0.75 | 0.06 | 0.84 |
| lh_S_orbital_lateral | 0.39 | -0.80 | 1.03 | 3.80 |
| lh_S_orbital_med-olfact | 0.39 | -0.41 | 1.73 | 8.42 |
| lh_S_orbital-H_Shaped | 0.53 | -1.14 | 0.16 | 1.27 |
| lh_S_parieto_occipital | 0.64 | -1.08 | 0.19 | 0.50 |
| lh_S_pericallosal | 0.24 | -0.13 | 0.92 | 1.75 |
| lh_S_postcentral | 0.63 | -1.53 | 0.09 | 0.51 |
| lh_S_precentral-inf-part | 0.58 | -1.57 | -0.01 | 0.83 |
| lh_S_precentral-sup-part | 0.48 | -1.14 | -0.21 | 0.90 |
| lh_S_suborbital | 0.32 | -0.38 | 1.31 | 6.09 |
| lh_S_subparietal | 0.51 | -1.12 | 0.56 | 2.81 |
| lh_S_temporal_inf | 0.45 | -0.66 | 0.54 | 5.47 |
| lh_S_temporal_sup | 0.67 | -1.45 | -0.23 | 0.73 |
| lh_S_temporal_transverse | 0.30 | -0.32 | 0.31 | 0.61 |
| lh_MeanThickness | 0.71 | -1.60 | -0.31 | 1.10 |
| rh_G&S_frontomargin | 0.42 | -0.80 | 1.16 | 5.18 |
| rh_G&S_occipital_inf | 0.30 | -0.53 | 0.21 | 0.49 |
| rh_G&S_paracentral | 0.38 | -0.35 | 0.18 | 0.76 |
| rh_G&S_subcentral | 0.45 | -0.85 | 0.13 | 0.27 |
| rh_G&S_transv_frontopol | 0.42 | -0.62 | 0.69 | 2.37 |
| rh_G&S_cingul-Ant | 0.62 | -1.29 | 0.40 | 2.06 |
| rh_G&S_cingul-Mid-Ant | 0.53 | -0.70 | 0.12 | 2.43 |
| rh_G&S_cingul-Mid-Post | 0.59 | -1.07 | 0.18 | 0.79 |
| rh_G_cingul-Post-dorsal | 0.51 | -0.99 | -0.37 | 2.26 |
| rh_G_cingul-Post-ventral | 0.26 | -0.16 | 0.11 | 0.49 |
| rh_G_cuneus | 0.46 | -0.31 | 0.90 | 3.93 |
| rh_G_front_inf-Opercular | 0.51 | -1.05 | 0.02 | 0.50 |
| rh_G_front_inf-Orbital | 0.39 | -0.50 | 0.16 | 0.86 |
| rh_G_front_inf-Triangul | 0.51 | -0.82 | 0.18 | 1.45 |
| rh_G_front_middle | 0.53 | -1.00 | -0.38 | 1.33 |
| rh_G_front_sup | 0.56 | -0.99 | -0.18 | 1.39 |
| rh_G_Ins_lg&S_cent_ins | 0.21 | -0.11 | -0.07 | 0.45 |
| rh_G_insular_short | 0.25 | -0.14 | -0.06 | 0.08 |
| rh_G_occipital_middle | 0.48 | -0.91 | -0.23 | 1.89 |
| rh_G_occipital_sup | 0.41 | -0.46 | 0.05 | 0.95 |
| rh_G_oc-temp_lat-fusifor | 0.38 | -0.71 | -0.15 | 1.09 |
| rh_G_oc-temp_med-Lingual | 0.50 | -0.35 | 0.66 | 2.67 |
| rh_G_oc-temp_med-Parahip | 0.27 | -0.20 | -0.42 | 0.77 |
| rh_G_orbital | 0.54 | -0.68 | 0.05 | 0.70 |
| rh_G_pariet_inf-Angular | 0.53 | -1.01 | -0.38 | 1.72 |
| rh_G_pariet_inf-Supramar | 0.53 | -1.16 | -0.29 | 0.92 |
| rh_G_parietal_sup | 0.43 | -0.98 | -0.11 | 0.91 |
| rh_G_postcentral | 0.36 | -0.53 | 0.19 | 0.40 |
| rh_G_precentral | 0.43 | -0.52 | -0.69 | 1.66 |
| rh_G_precuneus | 0.55 | -0.98 | -0.02 | 0.44 |
| rh_G_rectus | 0.35 | -0.56 | 0.60 | 2.12 |
| rh_G_subcallosal | 0.12 | -0.06 | 0.02 | 0.44 |
| rh_G_temp_sup-G_T_transv | 0.41 | -0.47 | 0.12 | 0.17 |
| rh_G_temp_sup-Lateral | 0.43 | -0.66 | -0.29 | 1.16 |
| rh_G_temp_sup-Plan_polar | 0.37 | -0.27 | -0.19 | 0.81 |
| rh_G_temp_sup-Plan_tempo | 0.50 | -1.00 | 0.09 | 0.25 |
| rh_G_temporal_inf | 0.36 | -0.90 | -0.37 | 1.54 |
| rh_G_temporal_middle | 0.43 | -0.81 | -0.38 | 1.23 |
| rh_Lat_Fis-ant-Horizont | 0.43 | -0.83 | 0.62 | 1.68 |
| rh_Lat_Fis-ant-Vertical | 0.30 | -0.64 | 0.84 | 2.23 |
| rh_Lat_Fis-post | 0.61 | -1.07 | 0.04 | 0.35 |
| rh_Pole_occipital | 0.41 | -0.26 | 0.51 | 1.54 |
| rh_Pole_temporal | 0.24 | -0.46 | -0.79 | 2.02 |
| rh_S_calcarine | 0.65 | -0.72 | 0.66 | 1.99 |
| rh_S_central | 0.63 | -0.92 | 0.28 | 1.03 |
| rh_S_cingul-Marginalis | 0.61 | -1.33 | 0.08 | 0.77 |
| rh_S_circular_insula_ant | 0.42 | -0.76 | 0.34 | 1.12 |
| rh_S_circular_insula_inf | 0.53 | -0.58 | 0.29 | 0.60 |
| rh_S_circular_insula_sup | 0.63 | -1.44 | 0.11 | 1.12 |
| rh_S_collat_transv_ant | 0.33 | -0.62 | 0.48 | 1.46 |
| rh_S_collat_transv_post | 0.40 | -0.59 | 0.51 | 1.40 |
| rh_S_front_inf | 0.63 | -1.60 | 0.39 | 2.23 |
| rh_S_front_middle | 0.58 | -1.43 | 0.46 | 1.66 |
| rh_S_front_sup | 0.54 | -1.49 | -0.11 | 0.59 |
| rh_S_interm_prim-Jensen | 0.42 | -0.99 | 0.44 | 1.51 |
| rh_S_intrapariet&P_trans | 0.60 | -1.57 | 0.10 | 0.83 |
| rh_S_oc_middle&Lunatus | 0.37 | -0.81 | 0.45 | 1.80 |
| rh_S_oc_sup&transversal | 0.49 | -1.03 | 0.12 | 0.96 |
| rh_S_occipital_ant | 0.39 | -0.75 | 0.12 | 1.65 |
| rh_S_oc-temp_lat | 0.36 | -0.76 | 0.34 | 2.53 |
| rh_S_oc-temp_med&Lingual | 0.52 | -0.57 | 0.05 | 0.72 |
| rh_S_orbital_lateral | 0.45 | -0.98 | 1.28 | 7.26 |
| rh_S_orbital_med-olfact | 0.49 | -0.63 | 0.79 | 3.40 |
| rh_S_orbital-H_Shaped | 0.55 | -1.24 | 0.36 | 1.10 |
| rh_S_parieto_occipital | 0.62 | -1.10 | 0.16 | 0.45 |
| rh_S_pericallosal | 0.27 | -0.16 | 0.81 | 1.41 |
| rh_S_postcentral | 0.61 | -1.38 | 0.14 | 0.37 |
| rh_S_precentral-inf-part | 0.57 | -1.47 | -0.06 | 0.57 |
| rh_S_precentral-sup-part | 0.45 | -0.96 | -0.34 | 1.31 |
| rh_S_suborbital | 0.25 | -0.25 | 0.53 | 0.60 |
| rh_S_subparietal | 0.52 | -1.04 | 0.24 | 0.54 |
| rh_S_temporal_inf | 0.42 | -0.71 | 0.32 | 1.94 |
| rh_S_temporal_sup | 0.63 | -1.57 | -0.19 | 0.83 |
| rh_S_temporal_transverse | 0.32 | -0.35 | 0.36 | 1.16 |
| MeanThickness | 0.71 | -1.54 | -0.18 | 0.84 |
| Left-Lateral-Ventricle | 0.37 | -0.26 | 0.55 | 0.51 |
| Left-Inf-Lat-Vent | 0.27 | -0.31 | 0.87 | 1.78 |
| Left-Cerebellum-White-Matter | 0.16 | -0.22 | -0.65 | 4.87 |
| Left-Cerebellum-Cortex | 0.27 | -0.18 | -1.75 | 14.02 |
| Left-Thalamus-Proper | 0.40 | -0.22 | -0.62 | 8.90 |
| Left-Caudate | 0.13 | -0.07 | -0.16 | 3.70 |
| Left-Putamen | 0.33 | -0.18 | -0.85 | 5.61 |
| Left-Pallidum | 0.47 | -0.44 | -0.57 | 5.08 |
| 3rd-Ventricle | 0.38 | -0.22 | 0.79 | 1.34 |
| 4th-Ventricle | 0.01 | 0.00 | 0.30 | 1.42 |
| Brain-Stem | 0.13 | -0.06 | -0.64 | 6.30 |
| Left-Hippocampus | 0.35 | -0.31 | -1.54 | 8.88 |
| Left-Amygdala | 0.35 | -0.28 | -1.20 | 6.45 |
| CSF | 0.26 | -0.16 | 0.50 | 2.88 |
| Left-Accumbens-area | 0.51 | -0.53 | -0.75 | 4.13 |
| Left-VentralDC | 0.40 | -0.30 | -0.64 | 7.99 |
| Left-vessel | 0.35 | -0.28 | 0.00 | 1.58 |
| Left-choroid-plexus | 0.73 | -0.55 | 0.20 | 1.13 |
| Right-Lateral-Ventricle | 0.39 | -0.26 | 0.49 | 0.69 |

**Appendix 0 Table 7. Patient Test Set Evaluation Metrics**
| ROI | EV | MSLL | Skew | Kurtosis |
| --- | --- | --- | --- | --- |
| Right-Inf-Lat-Vent | 0.24 | -0.26 | 0.88 | 2.16 |
| Right-Cerebellum-White-Matter | 0.12 | -0.16 | -0.67 | 6.55 |
| Right-Cerebellum-Cortex | 0.25 | -0.22 | -1.80 | 16.41 |
| Right-Thalamus-Proper | 0.49 | -0.32 | -0.39 | 4.50 |
| Right-Caudate | 0.18 | -0.10 | 0.09 | 2.29 |
| Right-Putamen | 0.36 | -0.21 | -0.91 | 4.40 |
| Right-Pallidum | 0.39 | -0.38 | -0.45 | 3.92 |
| Right-Hippocampus | 0.37 | -0.31 | -1.12 | 4.22 |
| Right-Amygdala | 0.36 | -0.38 | -0.94 | 3.38 |
| Right-Accumbens-area | 0.38 | -0.23 | -0.48 | 1.92 |
| Right-VentralDC | 0.40 | -0.32 | -0.31 | 5.74 |
| Right-vessel | 0.38 | -0.25 | -0.31 | 2.34 |
| Right-choroid-plexus | 0.80 | -0.85 | 0.17 | 0.89 |
| SubCortGrayVol | 0.43 | -0.23 | -0.99 | 9.58 |
| TotalGrayVol | 0.54 | -0.52 | -0.37 | 2.99 |
| SupraTentorialVolNotVent | 0.33 | -0.27 | -0.06 | 1.69 |
| avg_thickness | 0.72 | -1.66 | -0.22 | 0.82 |
| EstimatedTotalIntraCranialVol | 0.26 | -0.16 | 0.23 | 1.33 |

**Appendix 0 Table 8.**
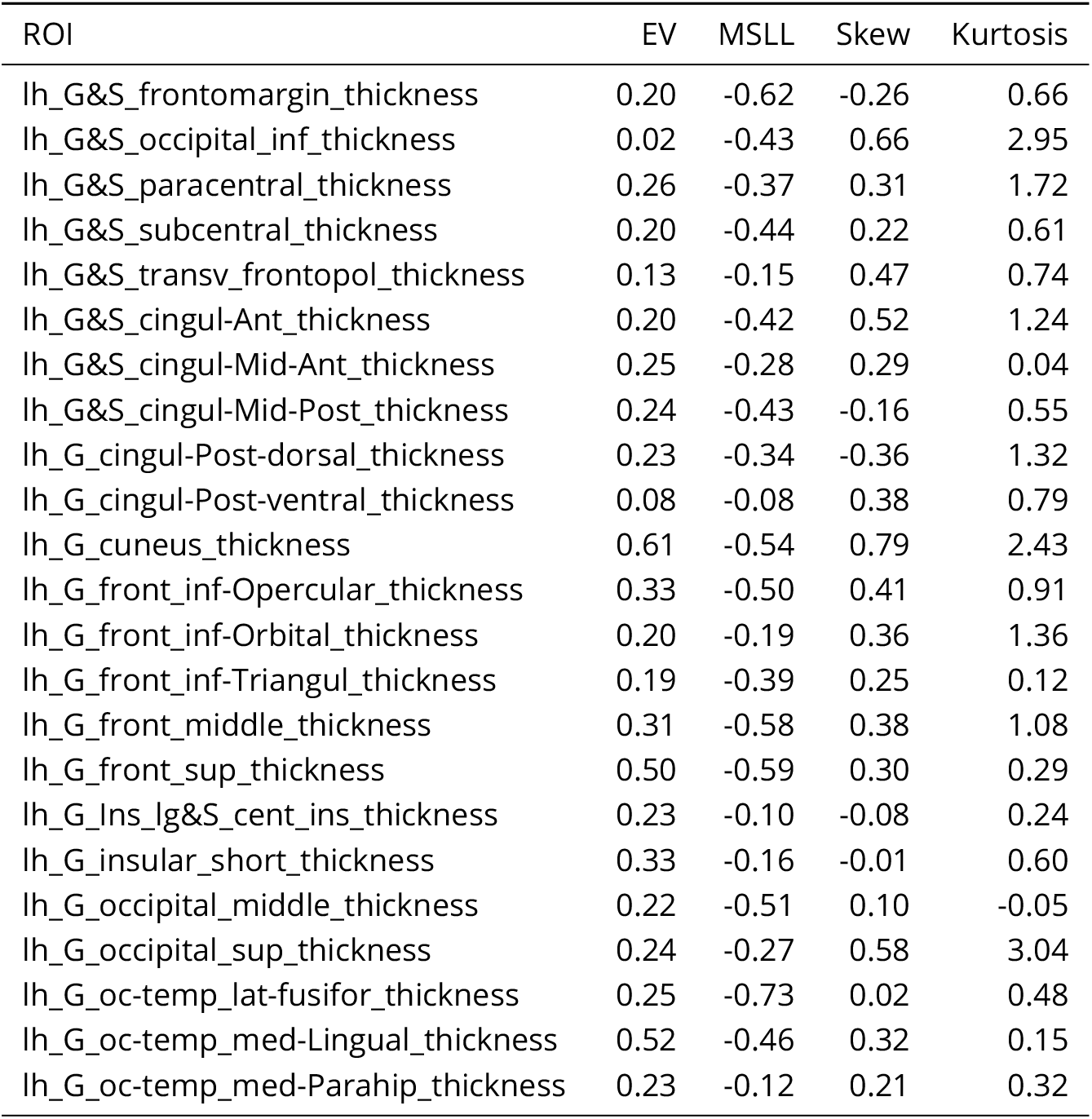
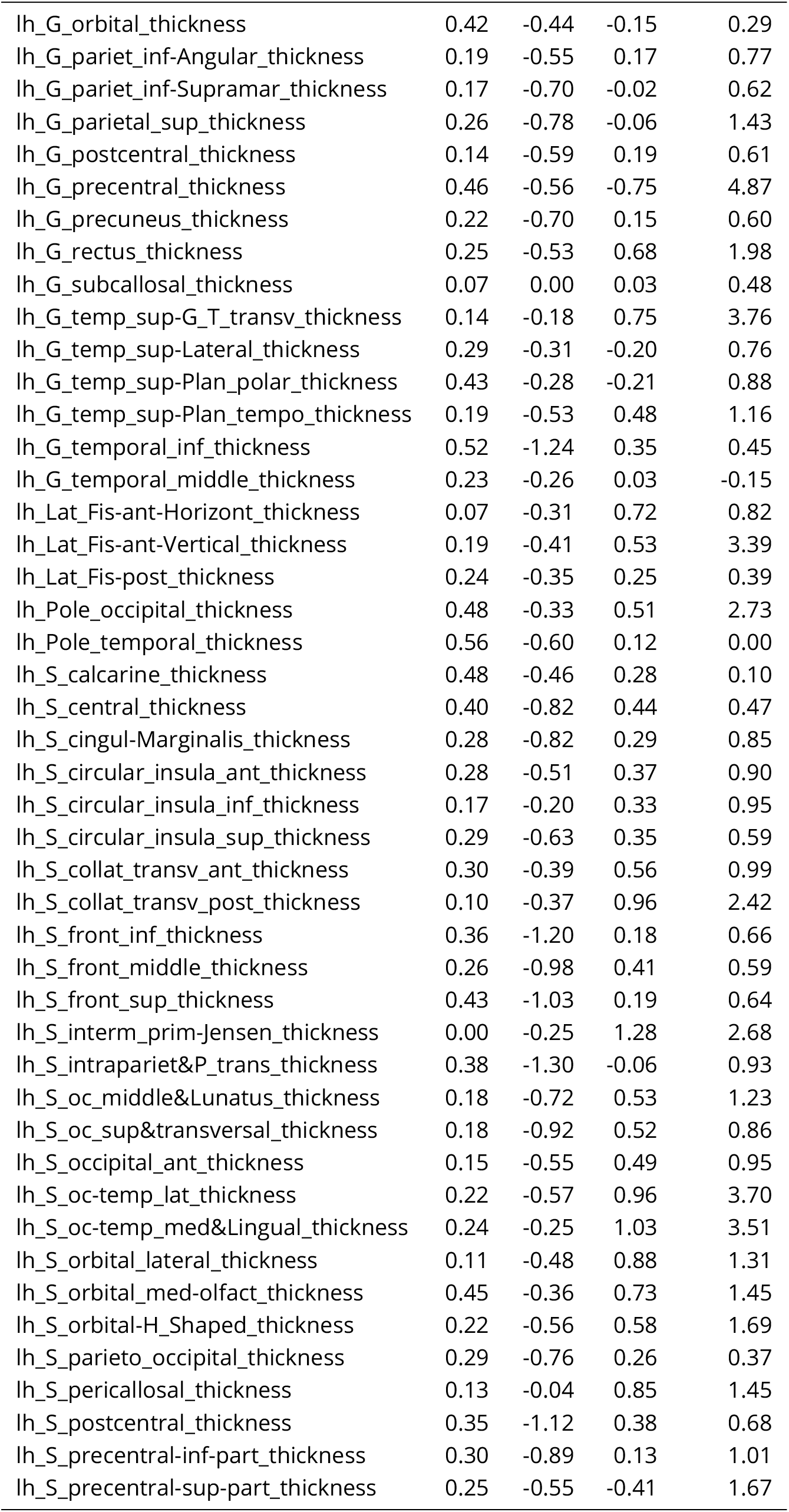
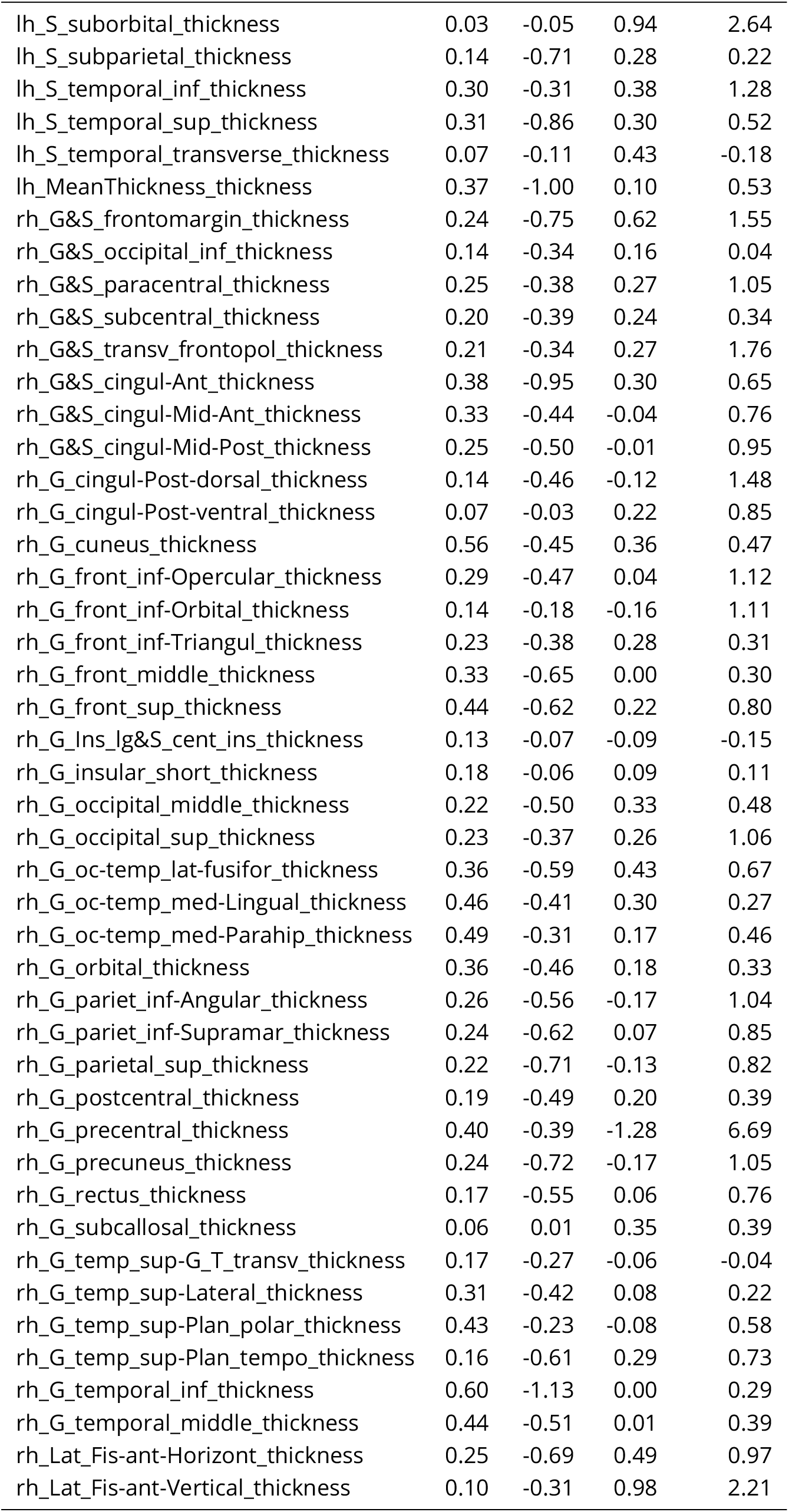
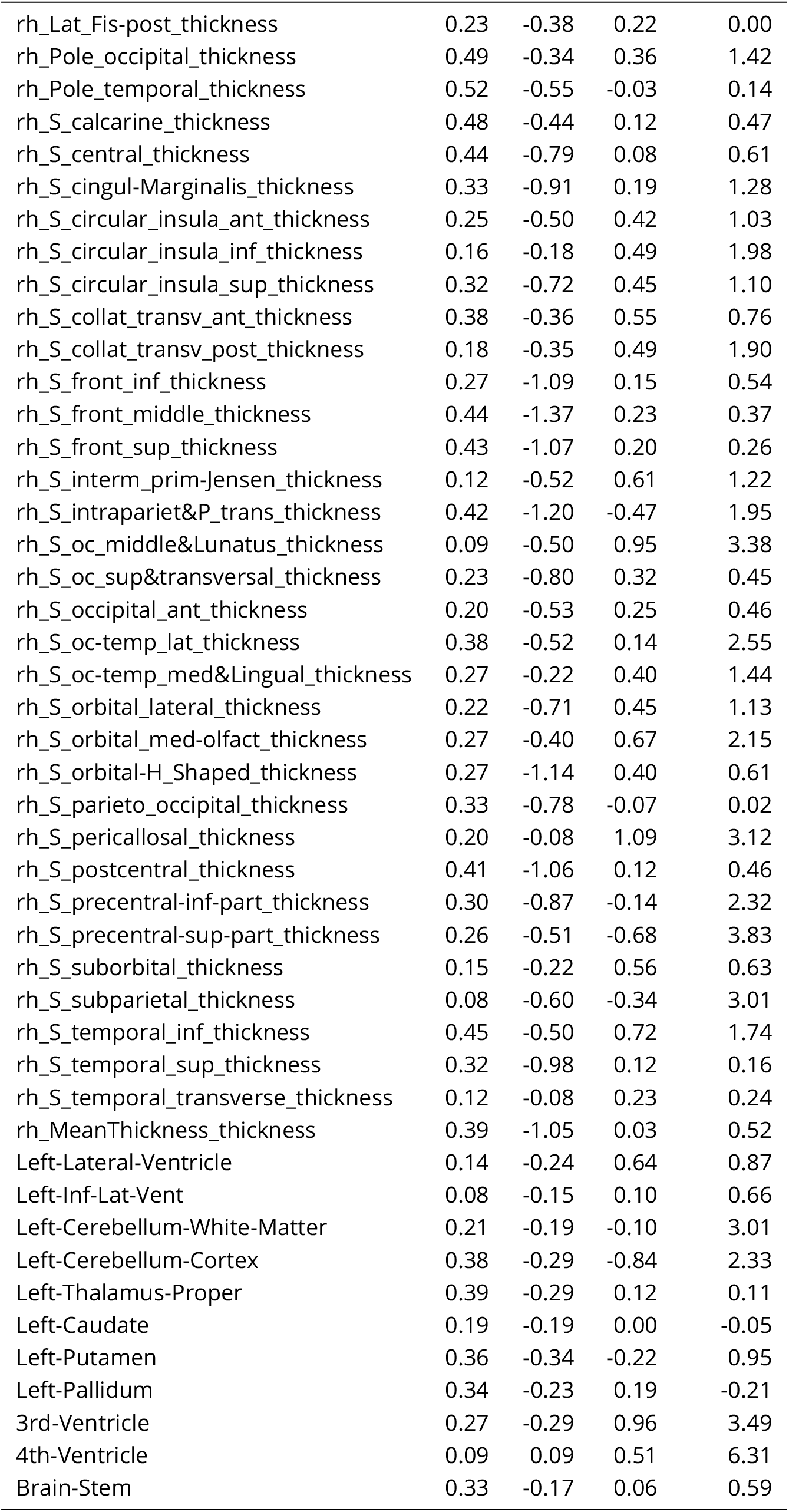
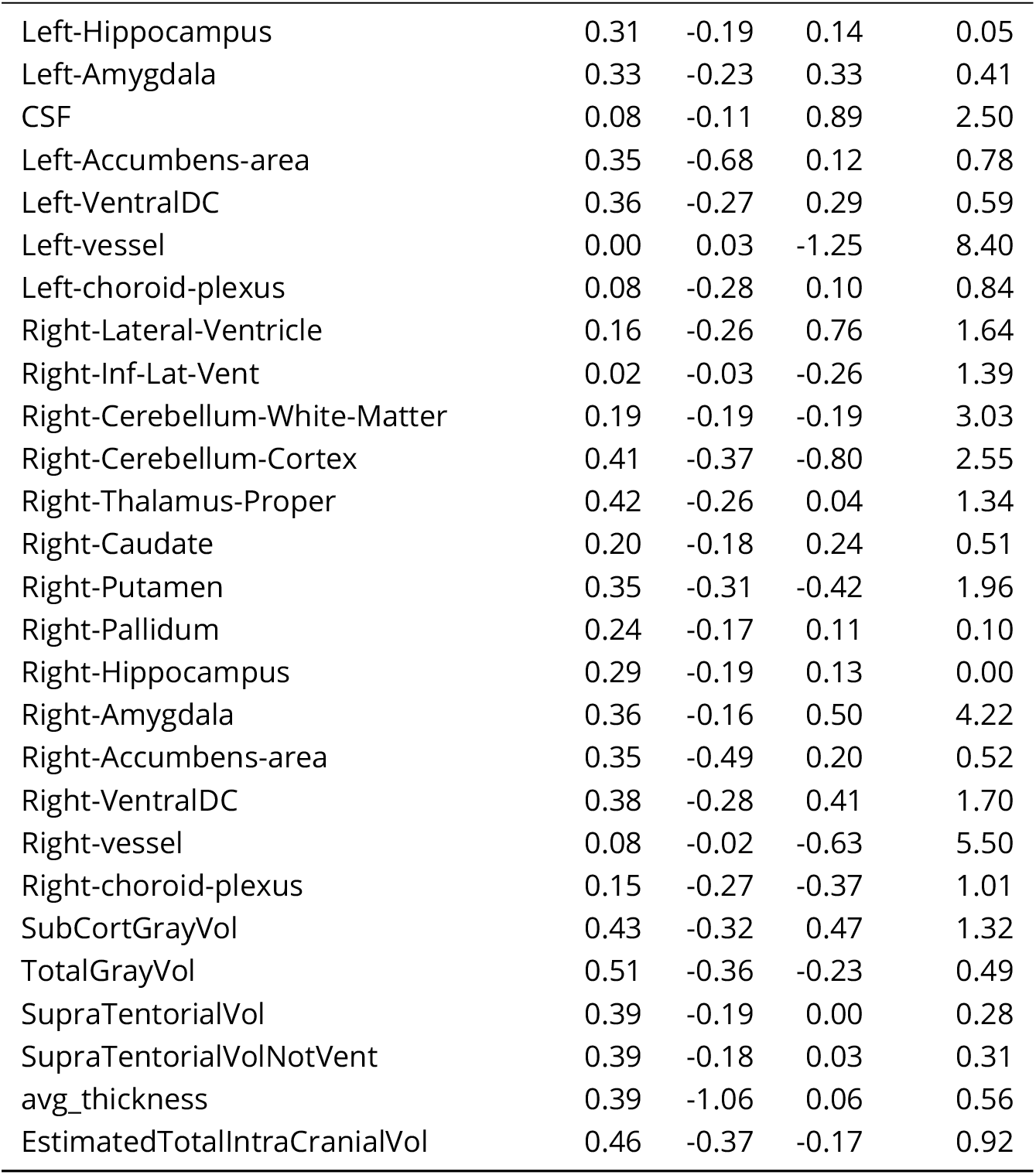
Transfer Test Set Evaluation Metrics

| ROI | EV | MSLL | Skew | Kurtosis |
| --- | --- | --- | --- | --- |
| lh_G&S_frontomargin_thickness | 0.20 | -0.62 | -0.26 | 0.66 |
| lh_G&S_occipital_inf_thickness | 0.02 | -0.43 | 0.66 | 2.95 |
| lh_G&S_paracentral_thickness | 0.26 | -0.37 | 0.31 | 1.72 |
| lh_G&S_subcentral_thickness | 0.20 | -0.44 | 0.22 | 0.61 |
| lh_G&S_transv_frontopol_thickness | 0.13 | -0.15 | 0.47 | 0.74 |
| lh_G&S_cingul-Ant_thickness | 0.20 | -0.42 | 0.52 | 1.24 |
| lh_G&S_cingul-Mid-Ant_thickness | 0.25 | -0.28 | 0.29 | 0.04 |
| lh_G&S_cingul-Mid-Post_thickness | 0.24 | -0.43 | -0.16 | 0.55 |
| lh_G_cingul-Post-dorsal_thickness | 0.23 | -0.34 | -0.36 | 1.32 |
| lh_G_cingul-Post-ventral_thickness | 0.08 | -0.08 | 0.38 | 0.79 |
| lh_G_cuneus_thickness | 0.61 | -0.54 | 0.79 | 2.43 |
| lh_G_front_inf-Opercular_thickness | 0.33 | -0.50 | 0.41 | 0.91 |
| lh_G_front_inf-Orbital_thickness | 0.20 | -0.19 | 0.36 | 1.36 |
| lh_G_front_inf-Triangul_thickness | 0.19 | -0.39 | 0.25 | 0.12 |
| lh_G_front_middle_thickness | 0.31 | -0.58 | 0.38 | 1.08 |
| lh_G_front_sup_thickness | 0.50 | -0.59 | 0.30 | 0.29 |
| lh_G_Ins_Ig&S_cent_ins_thickness | 0.23 | -0.10 | -0.08 | 0.24 |
| lh_G_insular_short_thickness | 0.33 | -0.16 | -0.01 | 0.60 |
| lh_G_occipital_middle_thickness | 0.22 | -0.51 | 0.10 | -0.05 |
| lh_G_occipital_sup_thickness | 0.24 | -0.27 | 0.58 | 3.04 |
| lh_G_oc-temp_lat-fusifor_thickness | 0.25 | -0.73 | 0.02 | 0.48 |
| lh_G_oc-temp_med-Lingual_thickness | 0.52 | -0.46 | 0.32 | 0.15 |
| lh_G_oc-temp_med-Parahip_thickness | 0.23 | -0.12 | 0.21 | 0.32 |
| lh_G_orbital_thickness | 0.42 | -0.44 | -0.15 | 0.29 |
| lh_G_pariet_inf-Angular_thickness | 0.19 | -0.55 | 0.17 | 0.77 |
| lh_G_pariet_inf-Supramar_thickness | 0.17 | -0.70 | -0.02 | 0.62 |
| lh_G_parietal_sup_thickness | 0.26 | -0.78 | -0.06 | 1.43 |
| lh_G_postcentral_thickness | 0.14 | -0.59 | 0.19 | 0.61 |
| lh_G_precentral_thickness | 0.46 | -0.56 | -0.75 | 4.87 |
| lh_G_precuneus_thickness | 0.22 | -0.70 | 0.15 | 0.60 |
| lh_G_rectus_thickness | 0.25 | -0.53 | 0.68 | 1.98 |
| lh_G_subcallosal_thickness | 0.07 | 0.00 | 0.03 | 0.48 |
| lh_G_temp_sup-G_T_transv_thickness | 0.14 | -0.18 | 0.75 | 3.76 |
| lh_G_temp_sup-Lateral_thickness | 0.29 | -0.31 | -0.20 | 0.76 |
| lh_G_temp_sup-Plan_polar_thickness | 0.43 | -0.28 | -0.21 | 0.88 |
| lh_G_temp_sup-Plan_tempo_thickness | 0.19 | -0.53 | 0.48 | 1.16 |
| lh_G_temporal_inf_thickness | 0.52 | -1.24 | 0.35 | 0.45 |
| lh_G_temporal_middle_thickness | 0.23 | -0.26 | 0.03 | -0.15 |
| lh_Lat_Fis-ant-Horizont_thickness | 0.07 | -0.31 | 0.72 | 0.82 |
| lh_Lat_Fis-ant-Vertical_thickness | 0.19 | -0.41 | 0.53 | 3.39 |
| lh_Lat_Fis-post_thickness | 0.24 | -0.35 | 0.25 | 0.39 |
| lh_Pole_occipital_thickness | 0.48 | -0.33 | 0.51 | 2.73 |
| lh_Pole_temporal_thickness | 0.56 | -0.60 | 0.12 | 0.00 |
| lh_S_calcarine_thickness | 0.48 | -0.46 | 0.28 | 0.10 |
| lh_S_central_thickness | 0.40 | -0.82 | 0.44 | 0.47 |
| lh_S_cingul-Marginalis_thickness | 0.28 | -0.82 | 0.29 | 0.85 |
| lh_S_circular_insula_ant_thickness | 0.28 | -0.51 | 0.37 | 0.90 |
| lh_S_circular_insula_inf_thickness | 0.17 | -0.20 | 0.33 | 0.95 |
| lh_S_circular_insula_sup_thickness | 0.29 | -0.63 | 0.35 | 0.59 |
| lh_S_collat_transv_ant_thickness | 0.30 | -0.39 | 0.56 | 0.99 |
| lh_S_collat_transv_post_thickness | 0.10 | -0.37 | 0.96 | 2.42 |
| lh_S_front_inf_thickness | 0.36 | -1.20 | 0.18 | 0.66 |
| lh_S_front_middle_thickness | 0.26 | -0.98 | 0.41 | 0.59 |
| lh_S_front_sup_thickness | 0.43 | -1.03 | 0.19 | 0.64 |
| lh_S_interm_prim-Jensen_thickness | 0.00 | -0.25 | 1.28 | 2.68 |
| lh_S_intrapariet&P_trans_thickness | 0.38 | -1.30 | -0.06 | 0.93 |
| lh_S_oc_middle&Lunatus_thickness | 0.18 | -0.72 | 0.53 | 1.23 |
| lh_S_oc_sup&transversal_thickness | 0.18 | -0.92 | 0.52 | 0.86 |
| lh_S_occipital_ant_thickness | 0.15 | -0.55 | 0.49 | 0.95 |
| lh_S_oc-temp_lat_thickness | 0.22 | -0.57 | 0.96 | 3.70 |
| lh_S_oc-temp_med&Lingual_thickness | 0.24 | -0.25 | 1.03 | 3.51 |
| lh_S_orbital_lateral_thickness | 0.11 | -0.48 | 0.88 | 1.31 |
| lh_S_orbital_med-olfact_thickness | 0.45 | -0.36 | 0.73 | 1.45 |
| lh_S_orbital-H_Shaped_thickness | 0.22 | -0.56 | 0.58 | 1.69 |
| lh_S_parieto_occipital_thickness | 0.29 | -0.76 | 0.26 | 0.37 |
| lh_S_pericallosal_thickness | 0.13 | -0.04 | 0.85 | 1.45 |
| lh_S_postcentral_thickness | 0.35 | -1.12 | 0.38 | 0.68 |
| lh_S_precentral-inf-part_thickness | 0.30 | -0.89 | 0.13 | 1.01 |
| lh_S_precentral-sup-part_thickness | 0.25 | -0.55 | -0.41 | 1.67 |
| lh_S_suborbital_thickness | 0.03 | -0.05 | 0.94 | 2.64 |
| lh_S_subparietal_thickness | 0.14 | -0.71 | 0.28 | 0.22 |
| lh_S_temporal_inf_thickness | 0.30 | -0.31 | 0.38 | 1.28 |
| lh_S_temporal_sup_thickness | 0.31 | -0.86 | 0.30 | 0.52 |
| lh_S_temporal_transverse_thickness | 0.07 | -0.11 | 0.43 | -0.18 |
| lh_MeanThickness_thickness | 0.37 | -1.00 | 0.10 | 0.53 |
| rh_G&S_frontomargin_thickness | 0.24 | -0.75 | 0.62 | 1.55 |
| rh_G&S_occipital_inf_thickness | 0.14 | -0.34 | 0.16 | 0.04 |
| rh_G&S_paracentral_thickness | 0.25 | -0.38 | 0.27 | 1.05 |
| rh_G&S_subcentral_thickness | 0.20 | -0.39 | 0.24 | 0.34 |
| rh_G&S_transv_frontopol_thickness | 0.21 | -0.34 | 0.27 | 1.76 |
| rh_G&S_cingul-Ant_thickness | 0.38 | -0.95 | 0.30 | 0.65 |
| rh_G&S_cingul-Mid-Ant_thickness | 0.33 | -0.44 | -0.04 | 0.76 |
| rh_G&S_cingul-Mid-Post_thickness | 0.25 | -0.50 | -0.01 | 0.95 |
| rh_G_cingul-Post-dorsal_thickness | 0.14 | -0.46 | -0.12 | 1.48 |
| rh_G_cingul-Post-ventral_thickness | 0.07 | -0.03 | 0.22 | 0.85 |
| rh_G_cuneus_thickness | 0.56 | -0.45 | 0.36 | 0.47 |
| rh_G_front_inf-Opercular_thickness | 0.29 | -0.47 | 0.04 | 1.12 |
| rh_G_front_inf-Orbital_thickness | 0.14 | -0.18 | -0.16 | 1.11 |
| rh_G_front_inf-Triangul_thickness | 0.23 | -0.38 | 0.28 | 0.31 |
| rh_G_front_middle_thickness | 0.33 | -0.65 | 0.00 | 0.30 |
| rh_G_front_sup_thickness | 0.44 | -0.62 | 0.22 | 0.80 |
| rh_G_Ins_Ig&S_cent_ins_thickness | 0.13 | -0.07 | -0.09 | -0.15 |
| rh_G_insular_short_thickness | 0.18 | -0.06 | 0.09 | 0.11 |
| rh_G_occipital_middle_thickness | 0.22 | -0.50 | 0.33 | 0.48 |
| rh_G_occipital_sup_thickness | 0.23 | -0.37 | 0.26 | 1.06 |
| rh_G_oc-temp_lat-fusifor_thickness | 0.36 | -0.59 | 0.43 | 0.67 |
| rh_G_oc-temp_med-Lingual_thickness | 0.46 | -0.41 | 0.30 | 0.27 |
| rh_G_oc-temp_med-Parahip_thickness | 0.49 | -0.31 | 0.17 | 0.46 |
| rh_G_orbital_thickness | 0.36 | -0.46 | 0.18 | 0.33 |
| rh_G_pariet_inf-Angular_thickness | 0.26 | -0.56 | -0.17 | 1.04 |
| rh_G_pariet_inf-Supramar_thickness | 0.24 | -0.62 | 0.07 | 0.85 |
| rh_G_parietal_sup_thickness | 0.22 | -0.71 | -0.13 | 0.82 |
| rh_G_postcentral_thickness | 0.19 | -0.49 | 0.20 | 0.39 |
| rh_G_precentral_thickness | 0.40 | -0.39 | -1.28 | 6.69 |
| rh_G_precuneus_thickness | 0.24 | -0.72 | -0.17 | 1.05 |
| rh_G_rectus_thickness | 0.17 | -0.55 | 0.06 | 0.76 |
| rh_G_subcallosal_thickness | 0.06 | 0.01 | 0.35 | 0.39 |
| rh_G_temp_sup-G_T_transv_thickness | 0.17 | -0.27 | -0.06 | -0.04 |
| rh_G_temp_sup-Lateral_thickness | 0.31 | -0.42 | 0.08 | 0.22 |
| rh_G_temp_sup-Plan_polar_thickness | 0.43 | -0.23 | -0.08 | 0.58 |
| rh_G_temp_sup-Plan_tempo_thickness | 0.16 | -0.61 | 0.29 | 0.73 |
| rh_G_temporal_inf_thickness | 0.60 | -1.13 | 0.00 | 0.29 |
| rh_G_temporal_middle_thickness | 0.44 | -0.51 | 0.01 | 0.39 |
| rh_Lat_Fis-ant-Horizont_thickness | 0.25 | -0.69 | 0.49 | 0.97 |
| rh_Lat_Fis-ant-Vertical_thickness | 0.10 | -0.31 | 0.98 | 2.21 |
| rh_Lat_Fis-post_thickness | 0.23 | -0.38 | 0.22 | 0.00 |
| rh_Pole_occipital_thickness | 0.49 | -0.34 | 0.36 | 1.42 |
| rh_Pole_temporal_thickness | 0.52 | -0.55 | -0.03 | 0.14 |
| rh_S_calcarine_thickness | 0.48 | -0.44 | 0.12 | 0.47 |
| rh_S_central_thickness | 0.44 | -0.79 | 0.08 | 0.61 |
| rh_S_cingul-Marginalis_thickness | 0.33 | -0.91 | 0.19 | 1.28 |
| rh_S_circular_insula_ant_thickness | 0.25 | -0.50 | 0.42 | 1.03 |
| rh_S_circular_insula_inf_thickness | 0.16 | -0.18 | 0.49 | 1.98 |
| rh_S_circular_insula_sup_thickness | 0.32 | -0.72 | 0.45 | 1.10 |
| rh_S_collat_transv_ant_thickness | 0.38 | -0.36 | 0.55 | 0.76 |
| rh_S_collat_transv_post_thickness | 0.18 | -0.35 | 0.49 | 1.90 |
| rh_S_front_inf_thickness | 0.27 | -1.09 | 0.15 | 0.54 |
| rh_S_front_middle_thickness | 0.44 | -1.37 | 0.23 | 0.37 |
| rh_S_front_sup_thickness | 0.43 | -1.07 | 0.20 | 0.26 |
| rh_S_interm_prim-Jensen_thickness | 0.12 | -0.52 | 0.61 | 1.22 |
| rh_S_intrapariet&P_trans_thickness | 0.42 | -1.20 | -0.47 | 1.95 |
| rh_S_oc_middle&Lunatus_thickness | 0.09 | -0.50 | 0.95 | 3.38 |
| rh_S_oc_sup&transversal_thickness | 0.23 | -0.80 | 0.32 | 0.45 |
| rh_S_occipital_ant_thickness | 0.20 | -0.53 | 0.25 | 0.46 |
| rh_S_oc-temp_lat_thickness | 0.38 | -0.52 | 0.14 | 2.55 |
| rh_S_oc-temp_med&Lingual_thickness | 0.27 | -0.22 | 0.40 | 1.44 |
| rh_S_orbital_lateral_thickness | 0.22 | -0.71 | 0.45 | 1.13 |
| rh_S_orbital_med-olfact_thickness | 0.27 | -0.40 | 0.67 | 2.15 |
| rh_S_orbital-H_Shaped_thickness | 0.27 | -1.14 | 0.40 | 0.61 |
| rh_S_parieto_occipital_thickness | 0.33 | -0.78 | -0.07 | 0.02 |
| rh_S_pericallosal_thickness | 0.20 | -0.08 | 1.09 | 3.12 |
| rh_S_postcentral_thickness | 0.41 | -1.06 | 0.12 | 0.46 |
| rh_S_precentral-inf-part_thickness | 0.30 | -0.87 | -0.14 | 2.32 |
| rh_S_precentral-sup-part_thickness | 0.26 | -0.51 | -0.68 | 3.83 |
| rh_S_suborbital_thickness | 0.15 | -0.22 | 0.56 | 0.63 |
| rh_S_subparietal_thickness | 0.08 | -0.60 | -0.34 | 3.01 |
| rh_S_temporal_inf_thickness | 0.45 | -0.50 | 0.72 | 1.74 |
| rh_S_temporal_sup_thickness | 0.32 | -0.98 | 0.12 | 0.16 |
| rh_S_temporal_transverse_thickness | 0.12 | -0.08 | 0.23 | 0.24 |
| rh_MeanThickness_thickness | 0.39 | -1.05 | 0.03 | 0.52 |
| Left-Lateral-Ventricle | 0.14 | -0.24 | 0.64 | 0.87 |
| Left-Inf-Lat-Vent | 0.08 | -0.15 | 0.10 | 0.66 |
| Left-Cerebellum-White-Matter | 0.21 | -0.19 | -0.10 | 3.01 |
| Left-Cerebellum-Cortex | 0.38 | -0.29 | -0.84 | 2.33 |
| Left-Thalamus-Proper | 0.39 | -0.29 | 0.12 | 0.11 |
| Left-Caudate | 0.19 | -0.19 | 0.00 | -0.05 |
| Left-Putamen | 0.36 | -0.34 | -0.22 | 0.95 |
| Left-Pallidum | 0.34 | -0.23 | 0.19 | -0.21 |
| 3rd-Ventricle | 0.27 | -0.29 | 0.96 | 3.49 |
| 4th-Ventricle | 0.09 | 0.09 | 0.51 | 6.31 |
| Brain-Stem | 0.33 | -0.17 | 0.06 | 0.59 |
| Left-Hippocampus | 0.31 | -0.19 | 0.14 | 0.05 |
| Left-Amygdala | 0.33 | -0.23 | 0.33 | 0.41 |
| CSF | 0.08 | -0.11 | 0.89 | 2.50 |
| Left-Accumbens-area | 0.35 | -0.68 | 0.12 | 0.78 |
| Left-VentralDC | 0.36 | -0.27 | 0.29 | 0.59 |
| Left-vessel | 0.00 | 0.03 | -1.25 | 8.40 |
| Left-choroid-plexus | 0.08 | -0.28 | 0.10 | 0.84 |
| Right-Lateral-Ventricle | 0.16 | -0.26 | 0.76 | 1.64 |
| Right-Inf-Lat-Vent | 0.02 | -0.03 | -0.26 | 1.39 |
| Right-Cerebellum-White-Matter | 0.19 | -0.19 | -0.19 | 3.03 |
| Right-Cerebellum-Cortex | 0.41 | -0.37 | -0.80 | 2.55 |
| Right-Thalamus-Proper | 0.42 | -0.26 | 0.04 | 1.34 |
| Right-Caudate | 0.20 | -0.18 | 0.24 | 0.51 |
| Right-Putamen | 0.35 | -0.31 | -0.42 | 1.96 |
| Right-Pallidum | 0.24 | -0.17 | 0.11 | 0.10 |
| Right-Hippocampus | 0.29 | -0.19 | 0.13 | 0.00 |
| Right-Amygdala | 0.36 | -0.16 | 0.50 | 4.22 |
| Right-Accumbens-area | 0.35 | -0.49 | 0.20 | 0.52 |
| Right-VentralDC | 0.38 | -0.28 | 0.41 | 1.70 |
| Right-vessel | 0.08 | -0.02 | -0.63 | 5.50 |
| Right-choroid-plexus | 0.15 | -0.27 | -0.37 | 1.01 |
| SubCortGrayVol | 0.43 | -0.32 | 0.47 | 1.32 |
| TotalGrayVol | 0.51 | -0.36 | -0.23 | 0.49 |
| SupraTentorialVol | 0.39 | -0.19 | 0.00 | 0.28 |
| SupraTentorialVolNotVent | 0.39 | -0.18 | 0.03 | 0.31 |
| avg_thickness | 0.39 | -1.06 | 0.06 | 0.56 |
| EstimatedTotalIntraCranialVol | 0.46 | -0.37 | -0.17 | 0.92 |

